# Single-cell multimodal glioma analyses reveal epigenetic regulators of cellular plasticity and environmental stress response

**DOI:** 10.1101/2020.07.22.215335

**Authors:** Kevin C. Johnson, Kevin J. Anderson, Elise T. Courtois, Floris P. Barthel, Frederick S. Varn, Diane Luo, Martine Seignon, Eunhee Yi, Hoon Kim, Marcos RH Estecio, Ming Tang, Nicholas E. Navin, Rahul Maurya, Chew Yee Ngan, Niels Verburg, Philip C De Witt Hamer, Ketan Bulsara, Michael L. Samuels, Sunit Das, Paul Robson, Roel GW Verhaak

## Abstract

Glioma intratumoral heterogeneity enables adaptation to challenging microenvironments and contributes to universal therapeutic resistance. Here, we integrated 914 single-cell DNA methylomes, 55,284 single-cell transcriptomes, and bulk multi-omic profiles across 11 adult IDH-mutant or IDH-wild-type gliomas to delineate sources of intratumoral heterogeneity. We found that local DNA methylation instability, or epimutation burden, was elevated in more aggressive tumors, reflected intratumoral variability, linked with transcriptional disruption, and associated with environmental stress response. We show that the activation of cell-state specific transcription factors is impacted by epimutations and that loosened epigenetic control may facilitate cellular plasticity. Our analyses support that somatic copy number alterations (SCNAs) promote epigenetic instability and that SCNAs largely precede epigenetic and transcriptomic diversification during glioma evolution. We confirmed the link between genetic and epigenetic instability by analyzing larger cohorts of bulk longitudinally collected and spatially separated DNA methylation data. Increased DNA methylation instability was associated with accelerated disease progression, and recurrently selected DNA methylation changes were enriched for environmental stress response pathways. Our work provides an integrative framework to better understand glioma evolution and highlights the importance of epigenetic heterogeneity in shaping therapeutic response.

## INTRODUCTION

Diffuse gliomas are the most common malignant brain tumors in adults and remain incurable. Extensive molecular characterization of glioma has defined genomic drivers and clinically relevant subtypes based on the presence of *IDH1/2* gene mutations (i.e., IDH-mutant and IDH-wild-type) (Cancer Genome Atlas Research et al., 2015; Ceccarelli et al., 2016; Louis et al., 2016). Inter- and intra-tumoral heterogeneity are salient features across glioma subtypes that contribute to the universal therapeutic resistance. The heterogeneity observed in surgical resection specimens reflects each tumor’s evolutionary path that is driven by competition between subpopulations harboring diverse genetic, epigenetic, and transcriptional aberrations (Barthel et al., 2019; Klughammer et al., 2018; Korber et al., 2019; Mazor et al., 2015; Wang et al., 2017). Thus, understanding how these different layers of heterogeneity integrate to define clonal lineages and drive glioma evolution may provide insights into treatment failure.

The study of tumor heterogeneity is complicated by cellular plasticity that enables cancer cells to reversibly transition between distinct cellular states in response to genetic, microenvironmental, and therapeutic stimuli (Flavahan et al., 2017). Single-cell RNA sequencing studies have previously identified such dynamic cellular states in IDH-wild-type gliomas (Bhaduri et al., 2020; Neftel et al., 2019; Wang et al., 2019; Yuan et al., 2018). Cell states of IDH-mutant gliomas were found to display a more restricted plasticity along a hierarchical differentiation axis (Tirosh et al., 2016; Venteicher et al., 2017). Epigenetic modifications, such as DNA methylation at cytosine followed by guanine dinucleotides (i.e., CpGs), are mitotically heritable marks and regulate cellular states (Easwaran et al., 2014). For example, the transition from a differentiated-like state to an undifferentiated, or stem-like, state following chemotherapy in glioma was accompanied by epigenetic reprogramming (Liau et al., 2017). However, the epigenetic mechanisms that enable cellular plasticity and regulate glioma cell states remain poorly understood.

Epimutation is aberrant DNA methylation resulting from errors in the placement or removal of epigenetic marks. These stochastic errors in DNA methylation replication can accumulate in cancer cells as passenger events or be evolutionarily selected by destabilizing gene expression programs. Accordingly, epimutations provide genetically identical tumor cells with greater plasticity to respond to environmental stressors (Flavahan et al., 2017). Previous studies of glioma have demonstrated associations between bulk tumor epigenetic heterogeneity metrics and clinical outcomes (Ceccarelli et al., 2016; Klughammer et al., 2018). Together, these findings suggest that stochastic DNA methylation alterations contribute to tumor heterogeneity and cellular plasticity and may drive clonal evolution of treatment-resistant phenotypes.

Single-cell DNA methylation technologies have recently emerged as tools to further dissect heterogeneous cell populations (Angermueller et al., 2016; Argelaguet et al., 2019; Farlik et al., 2016; Zhu et al., 2018b) and define epigenetic states that contribute to tumor evolution (Bian et al., 2018; Gaiti et al., 2019). Here, we integrated single-cell DNA methylomes, single-cell transcriptomes, and single-cell copy number profiles with bulk genomic profiles across a cohort of 11 glioma patient samples to deconstruct the sources of glioma heterogeneity. These analyses identified the gene regulatory regions most susceptible to stochastic DNA methylation alterations, the epigenetic modulation of transcriptional networks involved in glioma cellular identity, and that genetic driver events largely precede DNA methylation diversification during glioma evolution. We confirmed these single-cell findings through the association of DNA methylation instability across spatially separated and longitudinally collected bulk glioma tissue samples. Collectively, our work provides insights into the sources of intratumoral heterogeneity that fuel glioma evolution.

## RESULTS

### Single-cell DNA methylation highlights inter- and intratumoral heterogeneity at gene regulatory regions

To investigate glioma heterogeneity we performed single-cell DNA methylation, single-cell gene expression, and accompanying bulk tumor profiling in 11 adult patients with glioma (Figure 1A). This cohort was representative of two principal molecular subtypes (IDH-mutant and IDH-wild-type) and captured distinct clinical time points (i.e., unmatched initial and recurrent tumors, Table S1 and Figure S1). We mechanically dissected tumor specimens from the same geographic region dissociating tissue for single-cell protocols and flash freezing tissue for bulk genomic assays (Figure 1A). We implemented an established single-cell DNA methylation protocol, reduced representation bisulfite sequencing (scRRBS), and 10X Genomics’ single-cell gene expression protocol on cells from the same dissociation (Figure S2A) (Guo et al., 2015; Guo et al., 2013). Viable CD45^-^ (i.e., pan-immune cell marker) cells were plated for scRBBS, while single-cell transcriptomics was performed on all viable cells deriving a set of 914 single-cell methylomes and 55,284 single-cell transcriptomes (Methods). On average, ∼150,000 unique CpG dinucleotides covering representative chromosomal regions were measured per cell (Figure S2B-E), and expression was measured on an average of 2,340 genes per cell. Tumor cells were defined based on the detection of inferred copy number alterations in both datasets resulting in a final set of 844 tumor cells for single-cell DNA methylation and 30,831 tumor cells for single-cell transcriptomics (Methods, Figure S3A-I).

**Figure 1.**
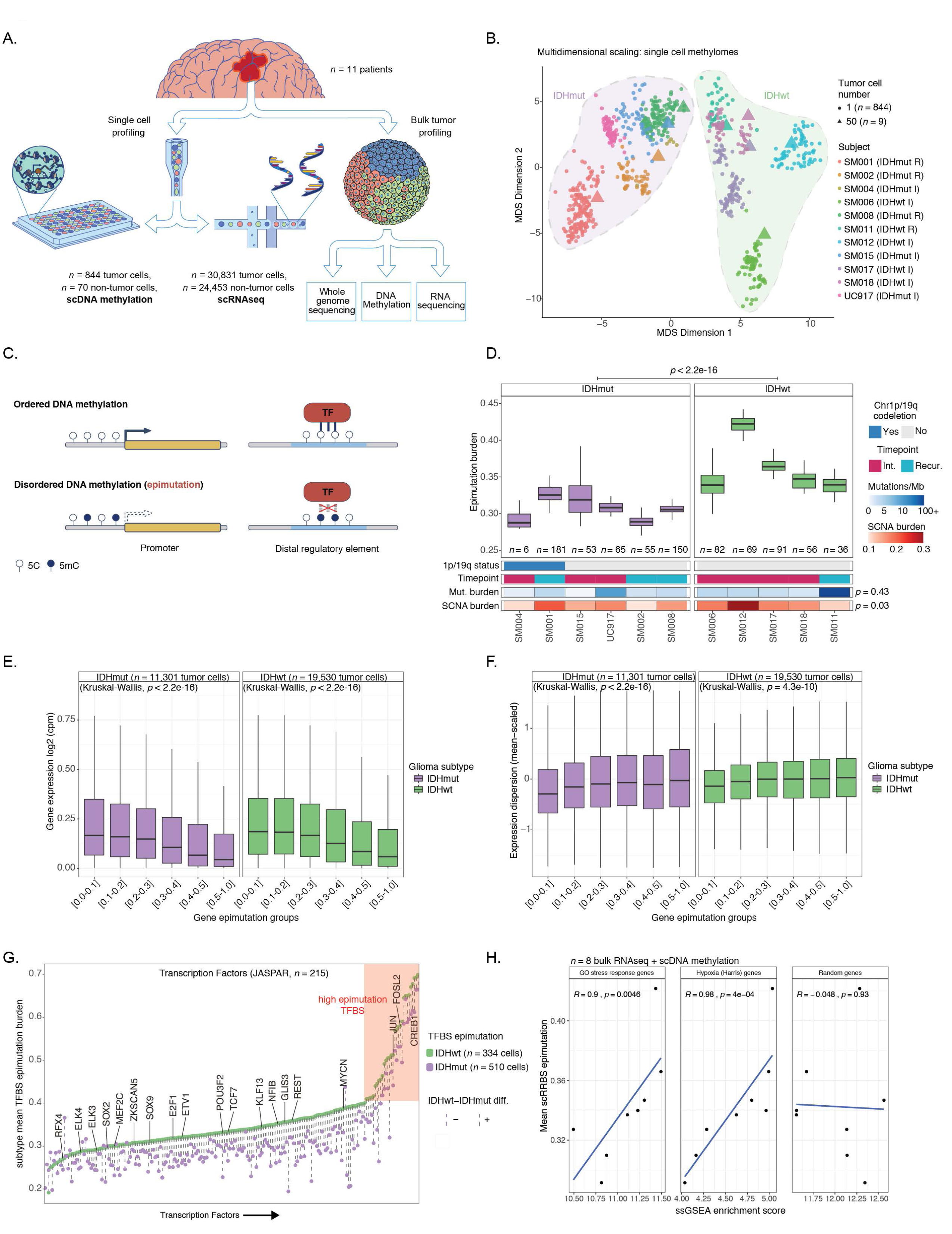
Single-cell DNA methylation sequencing highlights intratumoral heterogeneity and disruption of epigenetic regulatory mechanisms. (A) Schematic diagram detailing tumor sample processing and molecular profiling of single cells and bulk tumor samples (*n* = 11 subjects). (B) Multidimensional scaling (MDS) analysis using pairwise individual CpG distance metrics calculated between individual cells. Shapes represent whether a sample was a single tumor cell (*n* = 844 cells) or 50-tumor cells, *n* = 9/11 subjects). Colors indicate individual subjects, shaded regions indicated *IDH1*-mutation status of tumor, and annotation is provided indicating clinical timepoint (I = initial, R = recurrence). (C) Schematic depiction of local DNA methylation disorder in different genomic contexts. Left panel demonstrates epimutation, or local DNA methylation disorder, at the promoter region, where gene expression is disrupted by epimutation. The right panel provides an example of disrupted transcription factor binding due to epimutation. (D) Boxplots of tumor cell epimutation burden grouped by sample. Each boxplot spans the interquartile range with the whiskers representing the absolute range, excluding outliers. Wilcoxon rank sum *p*-value represents comparison between IDHmut and IDHwt epimutation burden. Each sample is annotated with clinical and molecular metrics with *p*-values indicating the relationship between sample mean epimutation burden and whole-genome sequencing derived somatic mutation burden or somatic alteration burden (Spearman correlation). (E) Boxplots of gene expression values, as log2 (counts per million), from single-cell RNAseq data across different gene epimutation groups. Gene epimutation groups are defined by the determining the mean epimutation value across a single gene. Color indicates *IDH1* mutation status. (F) Boxplots of gene expression dispersion. Expression profiles were mean-expression scaled to account for expression level-dependent variability across the same gene epimutation groups defined in panel E. (G) Scatterplot of the mean single-cell epimutation burden metric calculated across transcription factor binding sites (TFBSs) within a subtype, ordered by IDHwt TFBS epimutation. Each column represents a single transcription factor (TF) with a colored dotted line connecting IDHmut and IDHwt values. Names of TFs previously indicated to confer fitness advantages to glioma cells (MacLeod et al.) are listed above their TFBS epimutation burden estimate. (H) Scatterplot depicting the association between average single-cell epimutation burden estimate and single-sample Gene Set Enrichment Score for stress response, hypoxia, and random genes from bulk RNAseq data. Spearman correlation coefficient and *p*-values are indicated.

Unsupervised clustering and multidimensional scaling of the pairwise distances between single-cell genome-wide DNA methylation patterns grouped tumor cells by *IDH1* mutation status consistent with IDH-mutant tumors displaying greater genome-wide DNA methylation levels (Figure 1B and Figure S4A, Wilcoxon *p* < 2.2e-16). The co-localization of cells from different patients suggested some shared epigenetic states, while the isolated patient-specific grouping of 1 of 6 IDH-mutant and 2 out of 5 IDH-wild-type tumors suggested that genetic intertumoral heterogeneity identified by whole genome sequencing was also observed at the epigenetic level (Figure S1 and Figure 1B).

We next sought to determine the extent of intratumoral epigenetic heterogeneity by quantifying stochastic DNA methylation alterations in each single cell. In a non-diseased gene regulatory context, there is a general DNA methylation congruence in nearby CpGs reflecting tightly ordered gene regulation (Figure 1C top panel) (Kelsey et al., 2017). Epimutations reflect local DNA methylation disorder and may disrupt both local and distant gene regulation (Figure 1C bottom panel) (Easwaran et al., 2014). We constructed an epimutation burden metric per cell measured by the proportion of sequencing reads discordant for DNA methylation status as previously described (Gaiti et al., 2019; Klughammer et al., 2018; Landau et al., 2014). Cell-to-cell variation in epimutation burden was tumor dependent (Figure 1D) and was increased in tumor cells compared with non-tumor cells across IDH-mutant and IDH-wild-type glioma subtypes (Wilcoxon *p* < 0.0001, Figure S4B). The mean epimutation burden across a tumor’s single cells was not associated with the total somatic single nucleotide variant burden inferred through whole genome sequencing (Spearman correlation rho = 0.26, *p* = 0.43), independent of sequence context (Figure S4C). However, there was a positive association between the fraction of the genome with somatic copy number alterations (SCNA burden) and epimutation burden (Spearman correlation rho = 0.66, *p* = 0.03). Mutation burden reflects patient age (Alexandrov et al., 2020) and mutational processes (Figure S4D), while SCNA burden is associated with severed cell cycle checkpoints that compromises a cell’s ability to correct mis-segregations (Zhu et al., 2018a). The stronger relationship with SCNA burden suggested that epimutation burden increases with advanced disease rather than being elevated in the tumor cell of origin.

We next examined whether stochastic DNA methylation changes might impact levels of DNA methylation and transcriptional output. First, we determined both the epimutation and DNA methylation levels per gene, observing significant positive correlations across gene regions (Spearman correlation *p* < 2.2e-16, Figure S5A-B). An increase in stochastic DNA methylation might exert its functional impact by disrupting transcription programs (Gaiti et al., 2019). We leveraged companion single-cell RNAseq data to examine the association between epimutation burden, gene expression, and transcriptional variability. We observed both a reduction in mean expression (Kruskal Wallis *p* < 2.2e-16, Figure 1E) and an increase in gene dispersion (expression variability normalized for mean expression level) with increasing levels of epimutation burden in both IDH-mutant and IDH-wild-type tumors (Kruskal Wallis *p* < 2.2e-16, Figure 1F, Figure S5C-D), implying that epimutation contributes to gene expression dysregulation. To understand how aberrant DNA methylation could rewire broader regulatory networks, we performed a Gene Ontology enrichment analysis on genes with high epimutation (i.e., epimutation burden > 0.5) and genes with low epimutation (i.e., epimutation burden 0.0-0.1) (Methods), revealing that high epimutation genes were associated with processes involved in cellular differentiation and low epimutation genes were related with critical metabolic processes (Fisher’s Exact test, *p* < 0.05, Figure S5E-F). The enrichment results were consistent when using epimutation burden groupings from the promoter or gene body (Figure S5G-H). Together, these results suggest that cells may acquire adaptive cell states through the accumulation of epimutations that impair normal transcriptional and differentiation programs.

Beyond proximal dysregulation of gene expression, epimutations may impact the binding of key transcription factors, as changes in DNA methylation at DNA-binding motifs can positively or negatively impact transcription factor binding (Yin et al., 2017). To identify regulatory elements more prone to stochastic DNA methylation changes we determined the epimutation burden across the transcription factor binding sites (TFBS) listed in the JASPAR database (Methods, Figure 1G). Consistent with observed subtype-specific differences in methylation disorder, the majority of transcription factor binding sites had increased epimutation burden in IDH-wild-type compared with IDH-mutant cells. In both subtypes, transcription factors shown to be essential for glioma stem cell maintenance (e.g., SOX2, SOX9, etc. (MacLeod et al., 2019)) had lower than the median binding site epimutation burden suggesting that selection may act to deplete stochastic DNA methylation at these target regions (Figure 1G). In contrast, transcription factors that displayed high epimutation levels (Methods, Figure S6A) were associated with response to extracellular stimuli (Fisher’s Exact test, *p* < 0.05, Figure S6B). These findings suggest that increased epimutation levels at these environmental stress response regulators may facilitate an adaptive response to stressors such as hypoxia, which is commonly observed in glioma (Jin et al., 2017). To substantiate this association in the bulk glioma tissues, we performed single-sample Gene Set Enrichment Analyses (ssGSEA, Methods) using bulk RNAseq data and demonstrated robust associations between tumor average epimutation and positive stress response regulation (Spearman correlation rho = 0.9, *p* < 0.01) or cellular response to hypoxia (Spearman correlation rho = 0.98, p < 0.001 respectively), but not randomly selected genes (Spearman correlation rho = −0.05, *p* > 0.05, Figure 1H). Taken together, these results suggest that intratumoral variability in single-cell DNA methylation disorder may facilitate the adoption of distinct phenotypic states in response to stress stimuli.

### Integrative single-cell gene expression and DNA methylation analyses nominate epigenetic regulators of glioma cell states and stress response

To further examine the association between DNA methylation, stress response, and cellular states, we defined each tumor’s cellular composition from the single-cell transcriptional profiles. We performed unsupervised clustering analysis of all single cells and annotated clusters using marker genes (Figure 2A, Figure S7A-D) that revealed glial, immune, stromal, and malignant populations previously identified in glioma (Bhaduri et al., 2020; Wang et al., 2019). Malignant cells were broadly distributed over three cell states that all expressed canonical stem cell marker *SOX2* (Figure S7B). These pan-glioma states exist across both IDH-mutant and IDH-wild-type tumors, which we labelled as 1. differentiated-like, 2. stem-like, and 3. proliferating stem-like tumor cells, on the basis of marker gene expression (Figure 2A, Figure S7B, Table S2). Enumerating the proportion of pan-glioma malignant states by tumor of origin showed that IDH-mutant gliomas contained high fractions of stem-like cells (median 61%), while IDH-wild-type gliomas were dominated by differentiated-like cells (median 83%) and significantly higher fractions of proliferating stem-like cells (16% IDH-wild-type vs. 2% IDH-mutant, Wilcoxon *p* = 0.01, Figure 2B). Malignant cell state diversity (Shannon diversity index) was not associated with epigenetic (Spearman correlation rho = 0.12, *p* > 0.05) or genetic burden metrics (Spearman correlation rho = −0.18, *p* > 0.05, Figure 2B). Previously described malignant signatures of IDH-mutant glioma included Astrocyte-like and Oligodendrocyte-like cell types (Venteicher et al., 2017), which correspond to “differentiated-like” cells here. IDH-wild-type glioma cellular states (Neftel et al., 2019) included the “Astrocyte-like” and “Mesenchymal-like”, which were identified as “differentiated-like” in our clustering (Figure 2B, Figure S7D-F). The “proliferating stem-like” and “stem-like” states in our pan-glioma classification align closely with the “Undifferentiated” cells in IDH-mutant tumors and “Oligodendrocyte progenitor-like” and “Neural progenitor-like” in the IDH-wild-type tumors (Figure S7D-F), thus highlighting consistency of these pan-glioma signatures with previously reported IDH-subtype specific signatures (Neftel et al., 2019; Venteicher et al., 2017).

**Figure 2.**
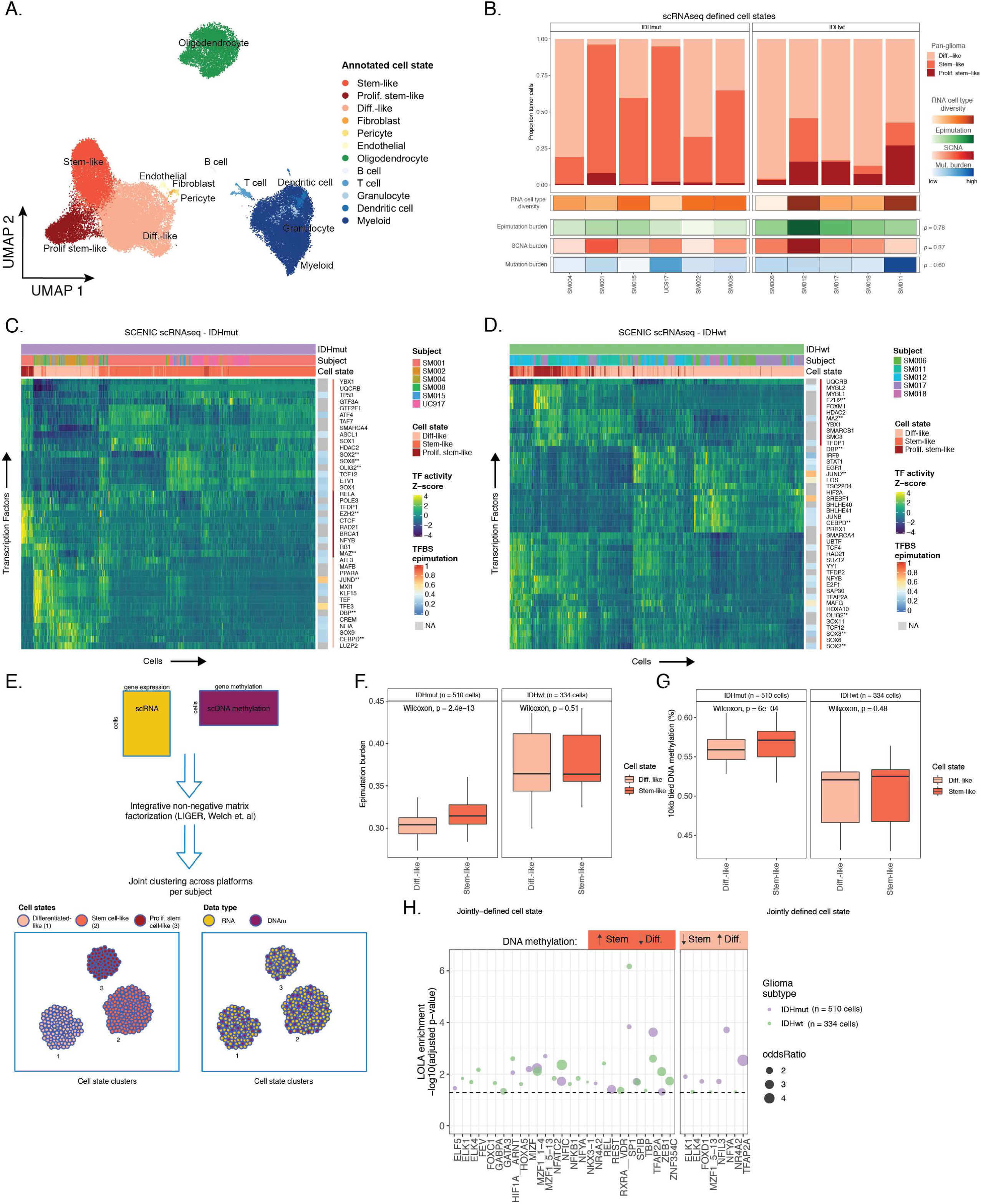
Integrative single-cell gene expression and DNA methylation analyses nominate epigenetic regulators of glioma cell state variability. (A) Uniform Manifold Approximation and Projection (UMAP) dimensionality reduction plot of scRNAseq data (*n* = 55,284 tumor cells, *n* = 11 subjects) showing the clustering of cell populations by transcriptionally defined cell state (point color) and labelled according to marker gene expression (Figure S6B). (B) Stacked bar plots representing the proportion of cellular states per tumor for pan-glioma classification. Each sample is annotated with molecular metrics with *p*-values indicating the relationship between cell type diversity, measured by Shannon’s entropy, and sample mean epimutation burden, whole-genome sequencing derived somatic alteration burden, or whole-genome sequencing derived somatic mutation burden (Spearman correlation). (C-D) Enriched transcription factor activity across pan-glioma cellular states determined using SCENIC algorithm and displayed as a heatmap of z-score enrichment values. Visualization is presented for the hierarchical clustering of 5,000 randomly selected tumor cells in both (C) IDHmut and (D) IDHwt tumors. (E) Schematic diagram representing LIGER workflow to jointly cluster single-cell RNAseq and DNA methylation data generated from the same tumor dissociation. (F) Boxplots representing the average epimutation burden in differentiated-like and stem-like populations in IDHmut (left panel) and IDHwt (right) tumors. (G) Boxplots representing the 10-kb tiled DNA methylation levels in differentiated-like and stem-like populations in IDHmut (left panel) and IDHwt (right) tumors. (H) Region set enrichment analysis for 10-kb tiles with higher DNA methylation in Stem-like (left panel) or Differentiated-like cells (right panel). Enrichment was determined by Locus Overlap Analysis (LOLA). Individual points represent enrichment of specific TFs in differentially methylated regions, color indicates results for specific IDH-mutant subtype, point size indicates log-odds ratio, and dotted line represents the statistical significance threshold (adjusted *p*-value < 0.05).

We next inferred gene regulatory networks from IDH-mutant and IDH-wild-type single-cell gene expression data to identify transcription factors (TFs) governing cell states (Methods) (Aibar et al., 2017). The inferred TF activity demonstrated that the three pan-glioma cell states are each regulated by a small set of TFs (Figure 2C-D). For example, stem-like tumor cells demonstrated the highest activity for known stem-cell regulators such as SOX2, SOX8, and OLIG2 in both the IDH-mutant and IDH-wild-type tumors (Figure 2C-D). In addition to high activity for these transcription factors, proliferating stem-like cells also had an overrepresentation of gene regulatory networks involved in chromatin remodeling and DNA repair such as those directed by EZH2 and BRCA1 (Figure 2C-D). In contrast, differentiated-like cells had higher transcription factor activities involved in astrocyte differentiation (i.e., SOX9) and response to stress stimuli (i.e., JUND, PPARA, HIF2A). We then tested whether the epimutation burden differed between cell state-specific transcription factors and did not find significant differences (Kolmogorov-Smirnov test *p* > 0.05 Figure S7G-H). However, several transcription factors associated with the differentiated-like cell state (e.g., JUND, TFE3, and SREBF1) were characterized by high epimutation levels, nominating them as regulators of cellular fitness (Figure 2C-D).

To define the epigenetic states of stem-like and differentiated-like cells in glioma, we used the linked inference of genomic experimental relationships (LIGER) method to identify shared properties between single-cell gene expression and DNA methylation data (Methods, Figure 2E) (Welch et al., 2019). We found that the distribution of tumor cell states within each sample was consistent between the two methods, as expected from the same tissue dissociation (Figure S8). We next investigated whether there were different levels of DNA methylation and epimutation between the two broad cell state classifications of stem-like (combining stem-like and proliferating stem-like) and differentiated-like. In IDH-mutant tumors, stem-like cells had significantly higher levels of both epimutation burden (*p* = 2.4e-13; Figure 2F left panel, Figure S9A) and DNA methylation (*p* = 6.0e-04, Figure 2G left panel, Figure S9B) likely reflecting elevated DNA methylation at genes responsible for cellular differentiation (Figure S5H). In IDH-wild-type tumors, which are marked by higher levels of epimutation and lower levels of DNA methylation compared with IDH-mutants, the differences between differentiated-like and stem-like cell populations demonstrated greater variability in both epimutation (*p* = 0.51; Figure 2F, Figure S9C) and DNA methylation (*p* = 0.48; Figure 2G and Figure S9D) suggesting loosened epigenetic control over cell states. To identify changes in DNA methylation between differentiated-like and stem-like cells in both IDH-wild-type and IDH-mutant glioma, we compared DNA methylation between cell states using a linear mixed effect model with tumor of origin as the random effect (Methods). Regions with increases in DNA methylation in stem-like cells were enriched for binding sites of SP1 and TFAP2A, two transcription factors that frequently cooperate in regulation of development associated genes (Figure 2H) (Orso et al., 2010). In addition, the analysis identified enrichment of increased DNA methylation at binding sites of the HIF1A/ARNT master transcriptional regulator of hypoxic response, in stem-like cells (Figure 2H). As increased DNA methylation at binding sites may result in reduced transcription factor binding efficiency, these results suggest that elevated cell stress transcription factor activity in differentiated-like cells may occur via dynamic epigenetic remodeling (Figure 2H). We found only a few regions where there was an increase in DNA methylation in differentiated-like cells (Figure 2H). Together, these results suggest that perturbing epigenetic control via epimutation may promote the adaptive cell states necessary to tolerate diverse stressful microenvironments, including hypoxia (Li et al., 2009) and therapy (Liau et al., 2017; Shaffer et al., 2017; Sharma et al., 2010).

### Somatic copy number alterations are associated with stochastic DNA methylation changes during disease evolution

We next investigated whether genetic stimuli could help explain the variability of epimutation burden across glioma cells. The fraction of genome with SCNAs was associated with epimutation burden at the bulk level (Spearman correlation rho = 0.66, *p* = 0.03, Figure 1D) and we confirmed this broad observation at the single-cell level (Spearman correlation rho = 0.50, *p* < 2.2e-16 IDH-mutant and rho = 0.72, *p* < 2.2e-16 IDH-wild-type, Figure 3A). To determine whether this relationship was driven by greater epimutation burden in copy number altered regions, we calculated the epimutation burden for each cell in copy number altered and non-altered regions. We did not observe a consistent relationship between epimutation burden and the copy number status in single-cell DNA methylation data (paired Wilcoxon *p* > 0.05, Figure S10A). Instead, most genomic regions displayed a similar epimutation burden independent of copy number status (e.g., SM012, Figure S10A) suggesting that aneuploid regions do not directly account for increases in epigenetic diversity, but that these somatic events are likely shaped by similar biological processes (e.g., replication errors). Late replicating regions of the genome tend to accumulate more DNA mutations and structural rearrangements (Koren et al., 2012), and we discovered that single-cell epimutation burden across both promoter and gene body regions was positively associated with later replication regions in both subtypes (Kruskal Wallis *p* < 1e-04, Figure 3B). Late replicating genomic regions may have reduced capacity to correct epimutations leading to their accumulation in a largely stochastic manner.

**Figure 3.**
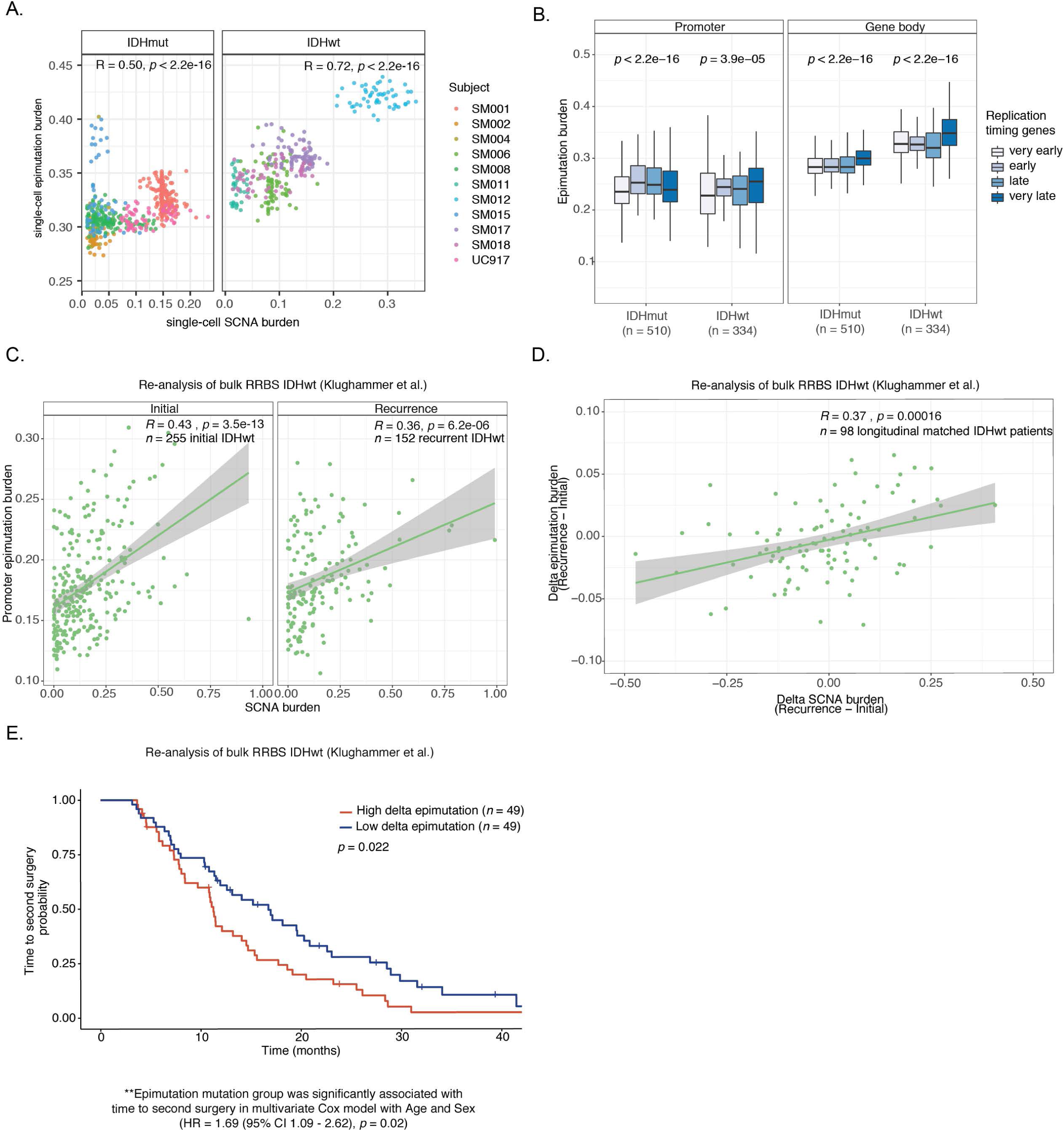
Somatic copy number alterations are associated with stochastic DNA methylation changes during disease evolution. (A) Scatterplot depicting the association between single-cell (*n* = 844 tumor cells) somatic copy number alteration (SCNA) and epimutation burden estimates by IDHmut (left panel) and IDHwt (right panel) subtypes. Points are colored by patient. Spearman correlation coefficients represent subtype-specific estimates. (B) Boxplots of epimutation burden calculated across the promoter (left panel) and gene body regions (right panel) based on different DNA replication times in IDHmut (*n* = 510) and IDHwt (*n* = 334) single cells. Kruskal-Wallis *p*-values indicate a test for differences across the replication time groupings separately for IDHmut and IDHwt cells (C) Scatterplot depicting the re-analysis of bulk promoter epimutation burden and SCNA burden in IDHwt initial (*n* = 255) and recurrent (*n* = 152) tumors (Klughammer et al.). Spearman correlation coefficients and *p*-values are presented for each independent timepoint. (D) Scatterplot depicting the association between bulk delta (subject-specific recurrence – initial estimates) SCNA burden and delta promoter epimutation burden in longitudinally profiled IDHwt tumors (*n* = 98 subjects, Klughammer et al.) Spearman correlation coefficient and p-value are presented. (E) Kaplan-Meier curve depicting time to second surgery in subjects where the change in epimutation burden between initial and recurrent disease was above (high, red) and below (low, blue) the median. Log-rank *p*-value for univariate analysis is presented within the figure. Hazard Ratio and *p*-value for change in epimutation burden are presented below for multi-variate Cox proportional hazard model including subject age and sex as predictors.

To validate the relationship between SCNA and epimutation burden in a larger cohort, we re-analyzed the bulk RRBS and copy number profiles of initial (*n* = 255 patients) and recurrent (*n* = 152 patients) IDH-wild-type gliomas, including a subset of longitudinally collected samples (n = 98 patients) (Klughammer et al., 2018). We confirmed our findings by demonstrating that SCNA burden was positively associated with epimutation burden at both initial and recurrent timepoints (Spearman correlation rho = 0.43, *p =* 3.5e-13 initial; rho = 0.36, *p* = 6.2e-06 recurrence, Figure 3C). A multivariable linear regression verified that this positive association between epimutation and SCNA burden was independent of subject age, tumor timepoint, and cellular proliferation (Figure S10B). To assess the relationship between longitudinal changes in SCNA burden and epimutation, we restricted our analysis to paired initial and recurrent samples and observed a positive association between increases in SCNA burden and epimutation (Spearman’s correlation rho = 0.37, *p* = 0.0002, Figure 3D). Furthermore, the highest increases in epimutation burden between initial and the recurrent tumor were associated with a shorter time to second surgery in both univariate (log-rank test *p* = 0.02, Figure 3F) and multivariate survival analyses (Cox proportional hazard model, HR = 1.69 95% CI (1.09 – 2.62), *p* = 0.02, Table S3) supporting that increased epigenetic instability is associated with accelerated disease progression. SCNA burden or aneuploidy results from mis-segregation during cell cycle, which can further perpetuate epimutations through aneuploid-induced metabolic and replication stress (Zhu et al., 2018a). The association between aneuploidy and epimutation uncovered here implicates that defective cell cycle checkpoints compromises genomic but also epigenetic integrity.

### Clonal evolution analyses highlight early somatic copy number evolution followed by epigenetic and transcriptomic diversification

The processes resulting in genetic, epigenetic, and transcriptomic heterogeneity may act at different times during tumor development. To evaluate somatic alteration timing and delineate intratumoral heterogeneity in the 11 glioma specimens, we reconstructed each tumor’s evolutionary history from bulk tumor whole genome sequencing data. Briefly, we determined the clonality of SCNAs and somatic point mutations assigning each genomic alteration to a tumor subclone (Methods). One to four genetic subclonal populations were detected per tumor, with linear and branched evolutionary patterns consistent with those previously observed in glioma (Kim et al., 2015; Korber et al., 2019). Assessment of the timing of genetic events revealed that chromosomal arm-level SCNA events were more likely to be classified as clonal (Fisher’s exact *p* = 0.03), while mutations at genes significantly mutated in glioma were more evenly distributed across subclones (56.1% classified as clonal in non-hypermutant tumors) (Methods, Figure 4A, Figure S11A-H). To identify any copy number alterations associated with DNA methylation states we compared phylogenetic and phyloepigenetic trees derived from single-cell DNA methylation data using an entanglement coefficient where a low value (i.e., 0 entanglement) indicates complete single cell alignment across the tree structures (Methods, Figure S12). We did not observe strong alignments between phylogenetic and phyloepigenetic trees (median value = 0.46, Figure 4B); instead DNA methylation profiles grouped by similar cell states, showing that, like transcriptomic cell states, epigenetic clones are distributed across genetic clones. Moreover, these results suggest that many large-scale copy number alterations occur as early events which propagate broad epigenetic diversification rather than simply affecting methylation state in proximal regions (Figure 4B, Figure S12A-K).

**Figure 4.**
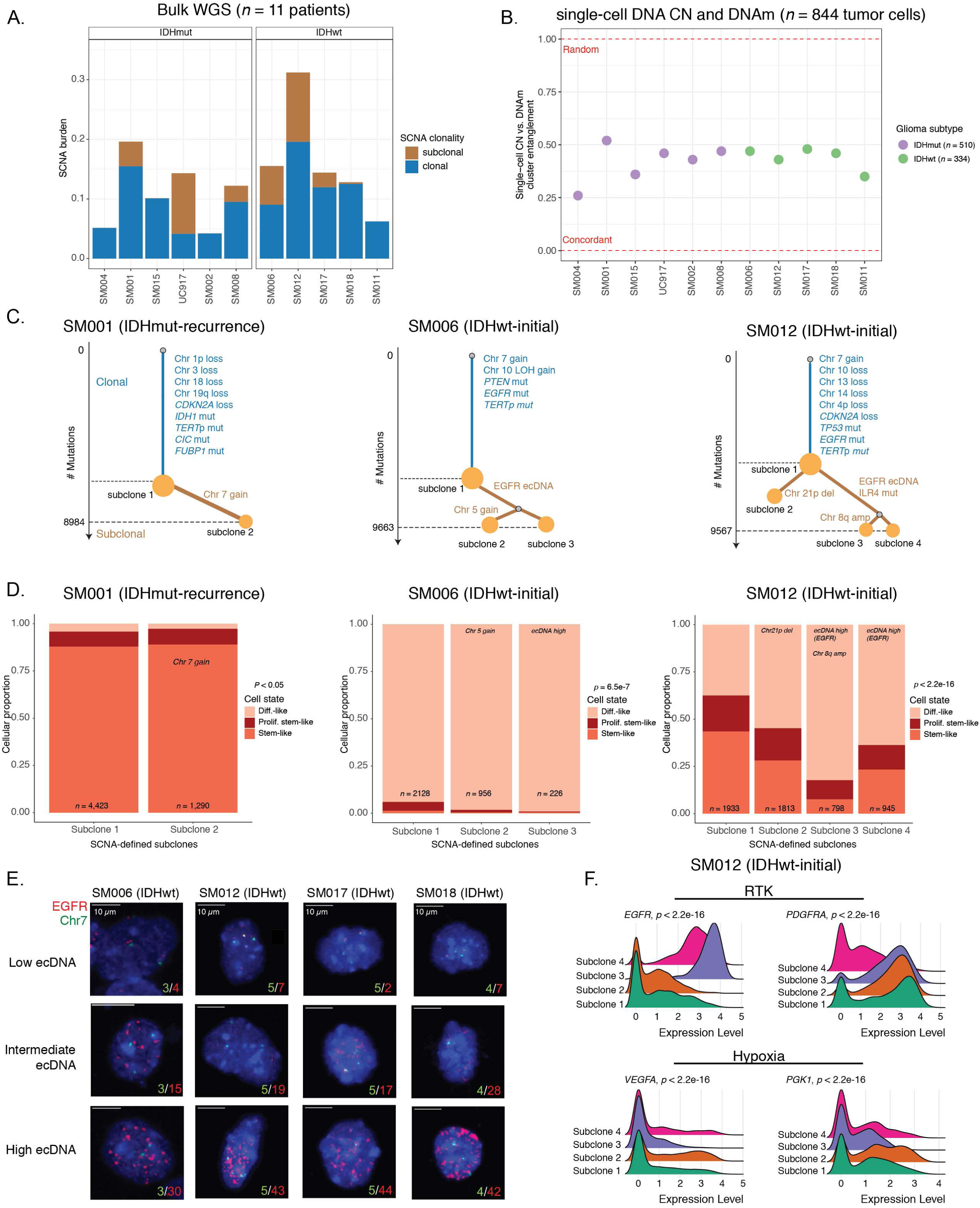
Clonal evolution analyses highlight early copy number evolution followed by epigenetic and transcriptomic diversification. (A) Stacked bar plots representing the proportion of whole-genome sequencing (WGS) derived somatic copy number alteration (SCNA) burden attributed to clonal vs. subclonal events. (B) Scatterplot depicting entanglement coefficients for tanglegrams comparing cluster dendrograms of scRRBS derived copy number and DNA methylation profiles. A coefficient of 0 indicates complete alignment of the tree structures, whereas a 1 indicates random association. Color indicates *IDH1* mutation status. (C) Examples of phylogenetic trees constructed from whole genome sequencing data (mutations and SCNAs) and further annotated using single-cell inferred copy number alterations (scRRBS + scRNAseq). Tree nodes represent alterations specific to the given clone, with node size corresponding to the fraction of cells with the associated alterations. Branch length scales with the number of mutations attributed to that clone. Clonal alterations are colored in blue, with subclonal alterations colored in gold. Genes considered significantly mutated in TCGA analyses (Ceccarelli et al., 2016) and chromosomal arm-level events are presented. (D) Single-cell RNAseq-derived cellular proportions separated by copy number-defined tumor subclone (Figure S3). Reported *p*-values represent Fisher’s exact test comparing the cellular state distributions across tumor subclones. (E) Representative Fluorescence *in* situ hybridization (FISH) images for IDHwt tumors computationally predicted to harbor *EGFR* extrachromosomal DNA (ecDNA) by whole genome sequencing (*n* = 4 patients). FISH images show *EGFR* amplifications (red) that occur distal to control chromosome 7 probes (green) indicating extrachromosomal status and high variability in copy number status across tumor cells. Scale bars = 10 microns. (F) Ridge plots of SM012 single-cell expression of receptor tyrosine kinase and hypoxia-associated genes, grouped by copy number-defined subclones. Reported p-values represent Wilcoxon Rank Sum tests comparing the gene expression of cells across tumor subclones.

We next asked whether genetic subclones within a tumor were associated with transcriptional diversity. We first used single-cell transcriptome inferred copy number profiles and found that three of eleven tumors (SM001, SM006, and SM012) had at least two distinct clones with chromosome arm-level alterations (Figure 4C, Figure S3). These tumors demonstrated significant shifts in cell state distributions across clones suggesting that the genetic heterogeneity also increases transcriptomic heterogeneity (per sample Fisher’s Exact test, *p* < 0.05, Figure 4D). Changes in transcriptional states may reflect cellular behaviors required to adapt to the varied microenvironmental niches, such as hypoxia, within a tumor.

Previous studies have demonstrated that *EGFR-*amplifying extrachromosomal DNA (ecDNA) elements are common in IDH-wild-type gliomas and enable widespread genomic heterogeneity through both the amplification of oncogenes as well as enhancer elements (deCarvalho et al., 2018; Morton et al., 2019; Wu et al., 2019). Therefore, we hypothesized that ecDNA may represent a particularly potent contributor to genomic heterogeneity whose impact extends to epigenetic and transcriptomic diversity (Verhaak et al., 2019; Wu et al., 2019). We detected ecDNAs by analyzing whole-genome sequencing data for our cohort and validated the variable distribution of extrachromosomal *EGFR* elements within a tumor using fluorescence *in situ* hybridization for *EGFR* (Figure 4E, Figure S13A-D). *EGFR* ecDNAs, like chromosomal arm level events (e.g., chr7 amplification in SM001) were able distinguish subsets of tumor cells (e.g. EGFR ecDNA in SM006) (Figure 4C, Figure S11F-G). We classified both single-cell DNA methylation and RNA profiles as ecDNA+ or ecDNA-based on *EGFR* copy number level (Figure S13E). We observed ecDNA+ cells had increased genome-wide DNA methylation in 3 of 4 cases (Wilcoxon *p* < 0.05, Figure S13F) and greater transcriptional diversity using gene count signatures compared with ecDNA-cells (Wilcoxon *p* < 0.05, Figure S13G, Methods) (Gulati et al., 2020). The tumor with the highest number of genetic subclones and epimutation burden (SM012) contained an *EGFR* amplifying ecDNA assigned to subclones 3 and 4 which were marked by differential expression of a receptor tyrosine kinase gene signature. The ecDNA(-) subclone 2 most closely associated with hypoxia gene expression (Wilcoxon *p* < 2.2e-16, Figure 4F), providing an example of how genetic heterogeneity may influence epigenetic and transcriptional reprogramming.

Taken together, our evolutionary analyses show that genetic evolution largely precedes epigenomic and transcriptomic diversification, and that intratumoral genetic heterogeneity influences but does not determine cell states.

### Integrated molecular trajectories supports adaptive DNA methylation changes under microenvironmental and therapeutic pressures

Our observation that genetic events likely precede epigenetic and transcriptomic diversification led us to ask whether epigenetic diversity accelerates tumor evolution by promoting cell survival in resource-deprived tumor environments (e.g., hypoxia or therapeutic exposures). To address this question and extend the generalizability of our findings, we sought to determine variable intratumoral DNA methylation levels in large-scale bulk glioma studies (Barthel et al., 2019; Ceccarelli et al., 2016; Verburg et al., 2020). Since these datasets were generated using DNA methylation microarrays, we used our single-cell DNA methylation data to define a microarray metric that quantified the DNA methylation instability of gene regions prone to epimutation (Figure 1E and Figure 5A). We reasoned that regions most susceptible to DNA methylation changes would reflect this stochasticity in bulk data by taking on intermediate DNA methylation values (Figure 5A). We confirmed that this epimutation, or DNA instability metric, approximated that of single-cell epimutation averages from the same tumor by comparing to microarray-derived profiles across the 11 tumors in our cohort (Spearman correlation rho = 0.65 *p* = 0.02, Figure S14A). We first applied this metric to The Cancer Genome Atlas (TCGA) data and found that the DNA methylation instability metric demonstrated differences across the TCGA-defined subtypes (Ceccarelli et al., 2016), with IDH-wild-type tumors displaying the highest levels (Kruskal Wallis *p* < 2.2e-16, Figure 5B). Integrating matching DNA methylation and RNAseq samples from 568 TCGA samples, we found that samples with higher levels of DNA methylation instability levels showed increased transcriptional activity of oxidative stress response genes, which corroborated our earlier finding of stronger positive associations between epigenetic instability and stress response regulation than randomly selected genes (Spearman rho = 0.47, *p* < 2.2e-16, *n* = 516 IDH-mutant initial tumors, rho = 0.31, *p* = 0.03, *n* = 52, IDH-wild-type initial tumors, Figure S14B-C).

**Figure 5.**
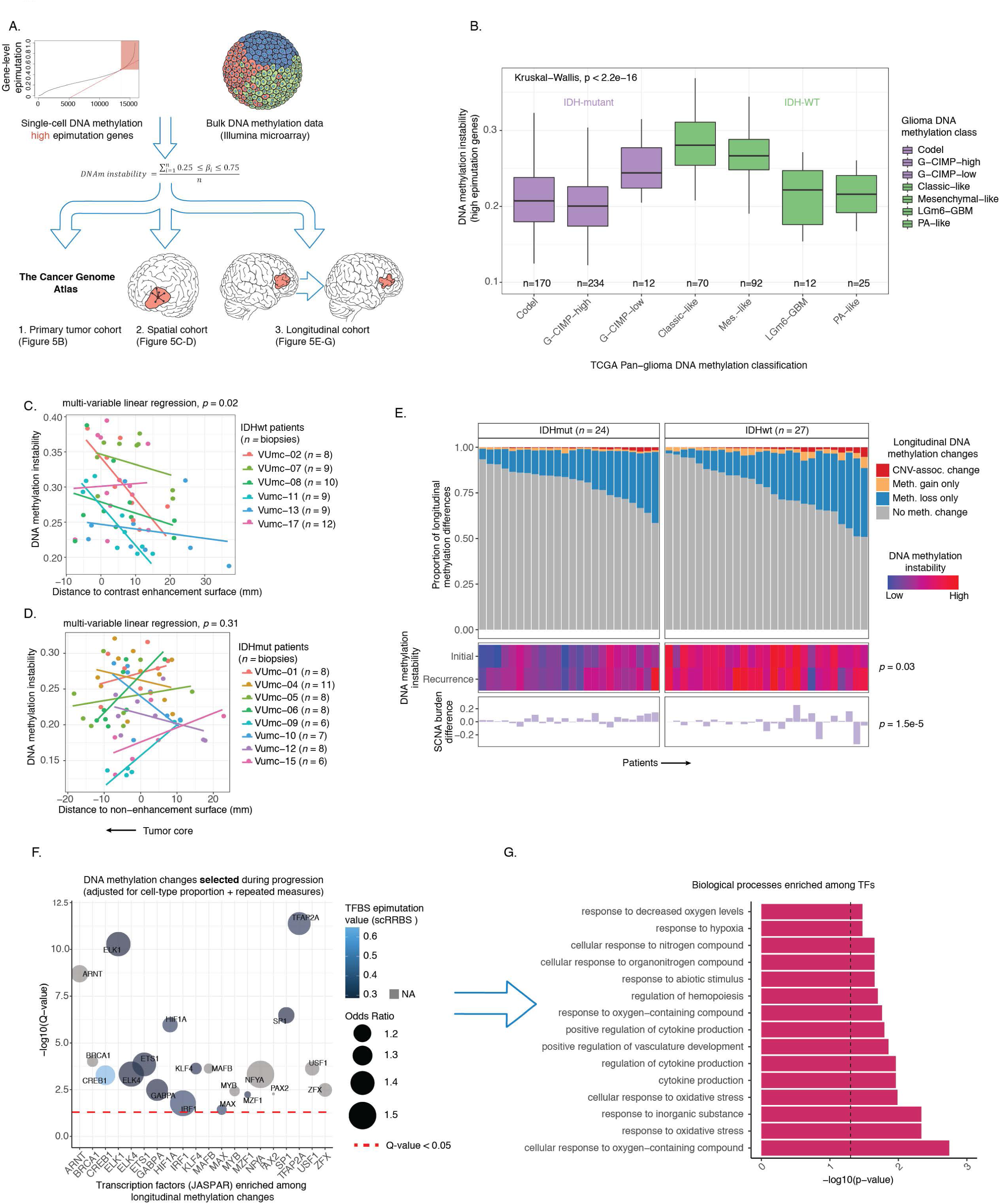
Integrated molecular trajectories supports adaptive DNA methylation changes under microenvironmental and therapeutic pressures. (A) Schematic workflow for construction of a DNA methylation instability metric in bulk cohorts informed by regions of high epimutation in single-cell DNA methylation data. The DNA methylation instability metric was calculated across bulk DNA methylation microarray data in a primary tumor cohort (TCGA), a cohort of multiple, spatially distinct biopsies from the same tumor (Verburg et al.), and a longitudinal cohort with accompanying genomic sequencing data (Glioma Longitudinal AnalySiS (GLASS), Barthel et al.). (B) Boxplots displaying the bulk DNA methylation instability metric calculated across previously described DNA-methylation based TCGA tumor classifications (Ceccarelli et al). Colors represent *IDH1*/*2* mutation status, and Kruskal-Wallis *p*-value testing for differences in distributions across classification is reported (*n* = 615 primary gliomas, *p* < 2.2e-16). (C-D) Scatterplots depicting distance from radiographic features plotted against the DNA methylation instability metric. Colors represent spatially separated biopsies from a single patient at initial clinical timepoint for (C) IDHwt tumors (*n* = 57 biopsies, *n* = 6 subjects) and (D) IDHmut tumors (*n* = 62 biopsies, *n* = 8 subjects). Linear regression lines colored by patient demonstrate the relationship between DNA methylation instability and radiographic features (i.e., contrast enhancement surface). The *p*-value reported from a multivariable linear regression model adjusting for subject represents the subtype-specific association between DNA methylation instability and radiographic feature. Biopsies taken closer to the tumor’s center (i.e., core) have the lowest value (left hand side of plot). (E) Each column represents an individual patient sampled across initial and recurrent timepoints and is separated into IDHmut (*n* = 24 subjects) and IDHwt (*n* = 27 subjects). Top panel, stacked bar plot represents the proportion of CpGs sites that experienced DNA methylation change associated with a subject-specific copy number change (defined by DNA sequencing data) between primary and recurrent disease (red), DNA methylation gain not associated with a CNV change (orange), DNA methylation loss not associated with a CNV change (blue), and no longitudinal DNA methylation change (gray). Middle panel, heatmap of DNA methylation instability metric in primary and recurrent disease (blue = low, red = high). Bottom panel, differences in SCNA burden between primary and recurrent tumor. All associated *p*-values represent Spearman correlations between absolute change in associated metric and the fraction of longitudinal DNA methylation differences. (F) Enrichment analysis for differentially methylated CpGs between primary and recurrent timepoints when adjusting for cellular composition, glioma subtype, and subject included as a random effect. Individual points represent enrichment of specific TFs in differentially methylated positions, color indicates the average TFBS epimutation burden from single-cell RRBS data (Figure 1G), point size indicates log-odds ratio, and dotted line represents the statistical significance threshold (*Q*-value < 0.05). (G) Gene Ontology enrichment of transcription factors associated with longitudinal DNA methylation changes. Dotted line represents threshold for statistical significance (Fisher’s exact test, *p* < 0.05).

We next applied the DNA methylation instability metric to 119 image-guided stereotactic biopsies taken from spatially distinct regions across IDH-wild-type (*n* = 57 biopsies, 6 patients) and IDH-mutant (*n* = 62 biopsies, *n* = 8 patients) tumors (Verburg et al., 2020). This enabled us to quantify the physical distance between a biopsied sample and specific radiographic features that delineate the tumor’s center (e.g., magnetic resonance imaging contrast-enhanced region, Figure S14D). We found an increase in DNA methylation instability closer to the tumor’s center across IDH-wild-type tumors while adjusting for patient (multivariable linear regression *p* = 0.02, Figure 5C), a region frequently characterized by hypoxia. The link between radiographic features and epigenetic shifts supports the association between cellular fitness and increased epigenetic plasticity. We did not observe a consistent relationship between tumor location and DNA methylation instability in IDH-mutant tumors (multivariable linear regression *p* = 0.31, Figure 5D) where hypoxia is less common.

The environmental pressures that tumors face may vary over time. To assess DNA methylation instability dynamics and its relationship with genetic alterations, we analyzed initial and recurrent tumor samples from the Glioma Longitudinal AnalySiS (GLASS) consortium for which DNA sequencing and DNA methylation data were available (*n* = 102 tumors, *n* = 51 patients). For each sample, we catalogued the specific copy number and DNA methylation alterations at individual CpG sites that changed between an initial tumor and its matched recurrence. Overall, we observed that DNA methylation changes were mostly decreases in DNA methylation consistent with previous findings (de Souza et al., 2018; Mazor et al., 2015), and that DNA methylation changes mainly occurred in regions that remained copy number stable between timepoints (Figure 5E). We then tested for DNA methylation changes following treatment while accounting for differences in cellular composition of the tumor microenvironment (Methods, Figure S14E). We discovered that regions with consistently altered DNA methylation independent of changes in microenvironment cell type distribution were enriched for the binding sites of transcription factors that regulate cellular stress response, particularly hypoxia (Methods, Figure 5F-G). We also observed the enrichment for differential DNA methylation among TFs that differed between stem-like and differentiated-like states in our single-cell data (e.g., SP1 and TFAP2A, Figure 5F and Figure 2H). These observations support our single-cell findings that regions with greatest epimutation levels are involved with processes regulating cellular differentiation and stress signaling. In summary, these results suggest that stochastic DNA methylation alterations can provide the variability necessary to enable transition to adaptive epigenetic phenotypes that are responsive to cellular stress (Figure 6).

**Figure 6.**
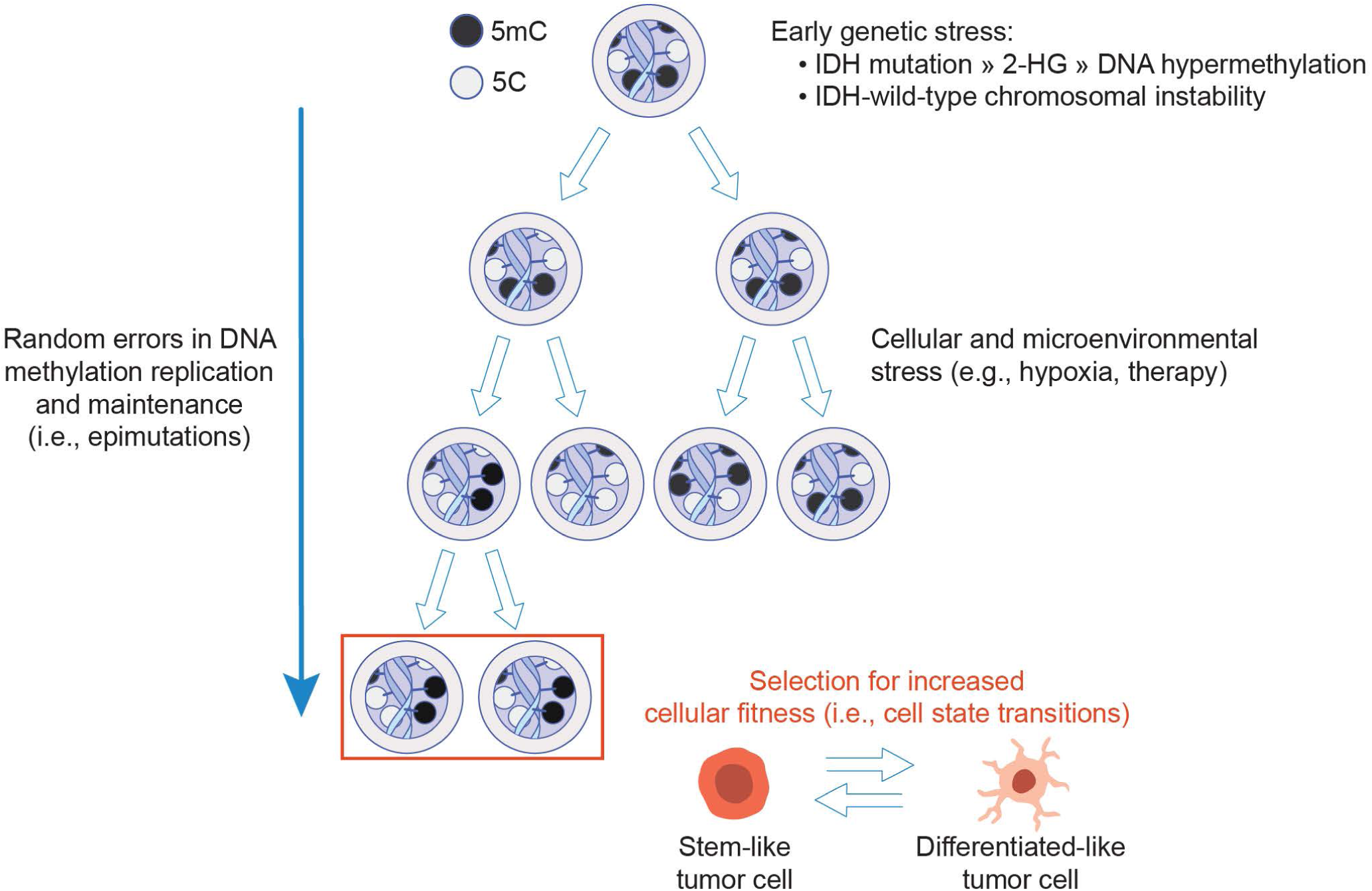
Model of epigenetic heterogeneity and tumor evolution. Schematic depiction of tumor evolution with general DNA methylation patterns represented by methylated (5-methylcytosine, 5mC) and unmethylated (5C) regions of the epigenome. Initiating genetic events such as *IDH1* and other driver mutations as well as somatic copy number alterations represent early stresses in glioma evolution that precipitate epigenetic heterogeneity. Both mutations in epigenetic enzymes and SCNAs can increase the likelihood of heritable DNA methylation alterations (i.e., epimutations). *IDH1* mutations result in the production of the oncometabolite 2-Hydroxyglutarate (2-HG) that leads to failure to remove aberrant DNA methylation while SCNAs can generate mitotic stress leading to the erosion of ordered DNA methylation. Non-genetic determinants further shape epigenetic heterogeneity as tumors evolve by exposing cells to spatially distinct microenvironmental stresses that impact the DNA methylation replication machinery. The subsequent epigenetic diversity provides an additional layer on which clonal evolution acts to select those cells with fitness-conferring epigenetic alterations. Ultimately, the loosened epigenetic control allows tumor cells to transition to cell states responsive to different selective pressures.

## DISCUSSION

Here, we integrated multimodal single-cell DNA methylation and transcriptomic profiles along with bulk profiles to interrogate the association between genetic tumor subclones, cellular states and epigenetic heterogeneity of glioma. We found that early genetic alterations largely precede epimutations, or stochastic changes in DNA methylation, whose accumulation throughout the genome led to dysregulated transcription and altered cellular states. Despite extensive intertumoral heterogeneity, we found recurrent epimutations localized to cellular differentiation genes and higher epimutation levels were associated with environmental pressures, such as hypoxia, highlighting a mechanism to overcome cell stress and enhance treatment resistance. Taken together, epigenetic intratumoral heterogeneity provides a plastic intermediate between genetic subclones and adaptive phenotypic cell states.

Epimutations increase a tumor population’s epigenetic diversity through random errors in the DNA methylation replication machinery (Klughammer et al., 2018; Landan et al., 2012; Landau et al., 2014). We found that genetic and environmental stimuli further induce epigenetic variability likely through altered cellular metabolism. Deregulated metabolism is a hallmark of glioma characterized by somatic mutations in the metabolic isocitrate dehydrogenase (IDH) genes and a hypoxic microenvironment in IDH-wild-type tumors. IDH-mutant glioma cells produce the oncometabolite 2-hydroxyglutarate (2HG) that interferes with DNA demethylation (Ceccarelli et al., 2016; Dang et al., 2009; Losman and Kaelin, 2013; Noushmehr et al., 2010; Turcan et al., 2012) leading to the observed high promoter epimutation levels at cellular differentiation genes and the predominance of a stem-like cell state. Across both subtypes, epimutation level was positively associated with broad chromosomal alterations, such as arm-level gains and losses, but not mutational burden. Copy number alterations occur during replicative crises that originate early in a tumor’s life history through punctuated evolution (Gao et al., 2016; Gerstung et al., 2020). We used a multimodal approach to link genetic clones across platforms and found that chromosomal alterations precede epigenetic and transcriptomic heterogeneity. The chromosomal imbalances may potentiate non-genetic diversity by accelerating cell proliferation (Taylor et al., 2018) and generating metabolic disruption via reactive oxygen species (Zhu et al., 2018a), thereby increasing the likelihood of aberrant DNA methylation. We also found that environmental stimuli, such as hypoxia, increase the rate of epimutation and is supported by a previous study that demonstrated hypoxia reduced the enzymatic activity of DNA methylation regulators (Thienpont et al., 2016). Tumor hypoxia is common across many cancers and could more broadly shape the phenotype of cells resistant to therapy through epimutation (Heddleston et al., 2010). Collectively, increased chromosomal alterations and more adverse microenvironments may explain the greater cell state diversity that exists in IDH-wild-type compared with IDH-mutant gliomas.

In a non-tumor setting, a cell’s epigenome reflects the tissue of origin and serves to stabilize cell state-specific gene expression (Roadmap Epigenomics et al., 2015). Epimutations may occur when this homeostasis is disrupted, enabling cells to acquire a de-differentiated malignant cell state or create an altered epigenetic landscape permissive to cell state transitions (Flavahan et al., 2017). Glioma cell states have been described to fall along axes of differentiation and proliferating potential (Bhaduri et al., 2020; Neftel et al., 2019; Venteicher et al., 2017; Wang et al., 2019). In accordance with prior reports, we observed pan-glioma malignant cell states that were found within each tumor. Our epigenetic single-cell profiles revealed that cell state-defining transcription factor activity can be perturbed by epimutation. Thus, diverse DNA methylation marks help to sustain multiple cell states that each confer their own fitness advantages and together accelerate disease progression.

Intratumoral heterogeneity in glioma reflects the Darwinian process of subclonal competition driven by limited nutrient access. While single-cell transcriptome-based phenotypes have investigated glioma transcriptomic heterogeneity (Bhaduri et al., 2020; Neftel et al., 2019; Tirosh et al., 2016; Venteicher et al., 2017; Wang et al., 2019), we have only limited knowledge on the degree of epigenetic variability. The intratumoral epigenetic variation defined here provides a link between Darwinian clone wars and phenotypic state changes by enabling diverse responses to selective pressures such as hypoxia and treatment. A better understanding of how to reprogram the glioma epigenome toward a more therapeutically vulnerable cell state will be needed to develop more effective interventions. In summary, single-cell epigenetic profiles show that each cell contains a unique set of methylation marks with distinct patterns regulating cellular states and reflecting variable levels of environmental stress.

## ACKNOWLEDGEMENTS

We would like to thank the patients and their families for their generous donation to biomedical research. We would also like to thank the staff in the following groups at The Jackson Laboratory for Genomic Medicine: single cell biology laboratory, flow cytometry core, and genomic technology core for assistance in data generation. We thank Matt Wimsatt for assistance in graphic design. We thank the University of Texas MD Anderson Epigenomics Profiling Core for their assistance in helping troubleshoot the scRRBS protocol. We thank Sheng Li and Samirkumar B. Amin for helpful feedback that strengthened the work. This work was supported by NIH grants R01 CA237208, R21 NS114873 and Cancer Center Support Grant P30 CA034196; Department of Defense W81XWH1910246 (R.G.W.V). F.S.V. is supported by a postdoctoral fellowship from The Jane Coffin Childs Memorial Fund for Medical Research. F.P.B. is supported by the National Cancer Institute (K99 CA226387). E.Y. is a fellow of the American Brain Tumor Association. K.C.J. is the recipient of an American Cancer Society Fellowship (130984-PF-17-141-01-DMC).

## AUTHOR CONTRIBUTIONS

K.C.J. and R.G.W.V. conceived the project and designed the experiments.

K.B. and S.D. curated patient samples and patient annotation.

K.C.J., M.R.H.E., M.T., N.N., R.M., C.N., M.S, and P.R performed single-cell library optimization and sequencing.

K.C. J. led data production and performed experiments with D.L., E.Y., and E.T.C.

K.C.J and K.J.A. led data analysis in collaboration with F.P.B., F.S.V., M.S., E.Y., and H.K. K.C.J., K.J.A., and R.G.W.V. wrote the manuscript with input from all authors.

## DECLARATIONS OF INTEREST

R.G.W.V. is a co-founder of and has received research support from Boundless Bio, Inc.

---

**Figure S1.**
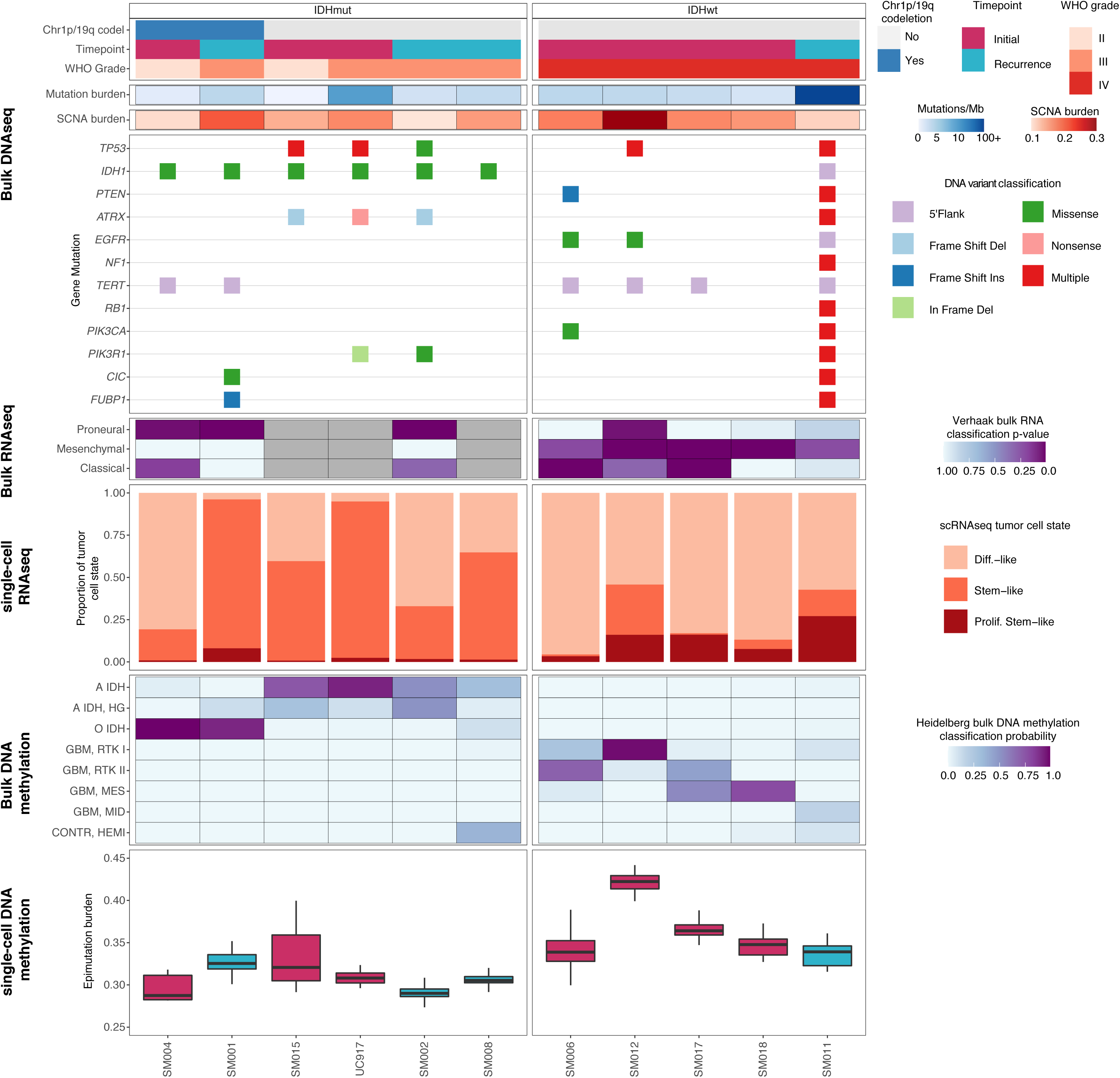
**Integrated molecular profiles of patient samples. Related to Figure 1.** Each patient is in a single column with data presented to indicate clinical features (top), followed by genetic alterations defined from bulk whole genome sequencing data, bulk RNA sequencing based subtype classification probabilities (Wang et al., *n* = 8 available), single-cell RNA tumor cellular state proportions, bulk DNA methylation microarray subtype classification probabilities (Capper et al.), and boxplots of single-cell epimutation burden with samples colored by clinical timepoint.

**Figure S2.**
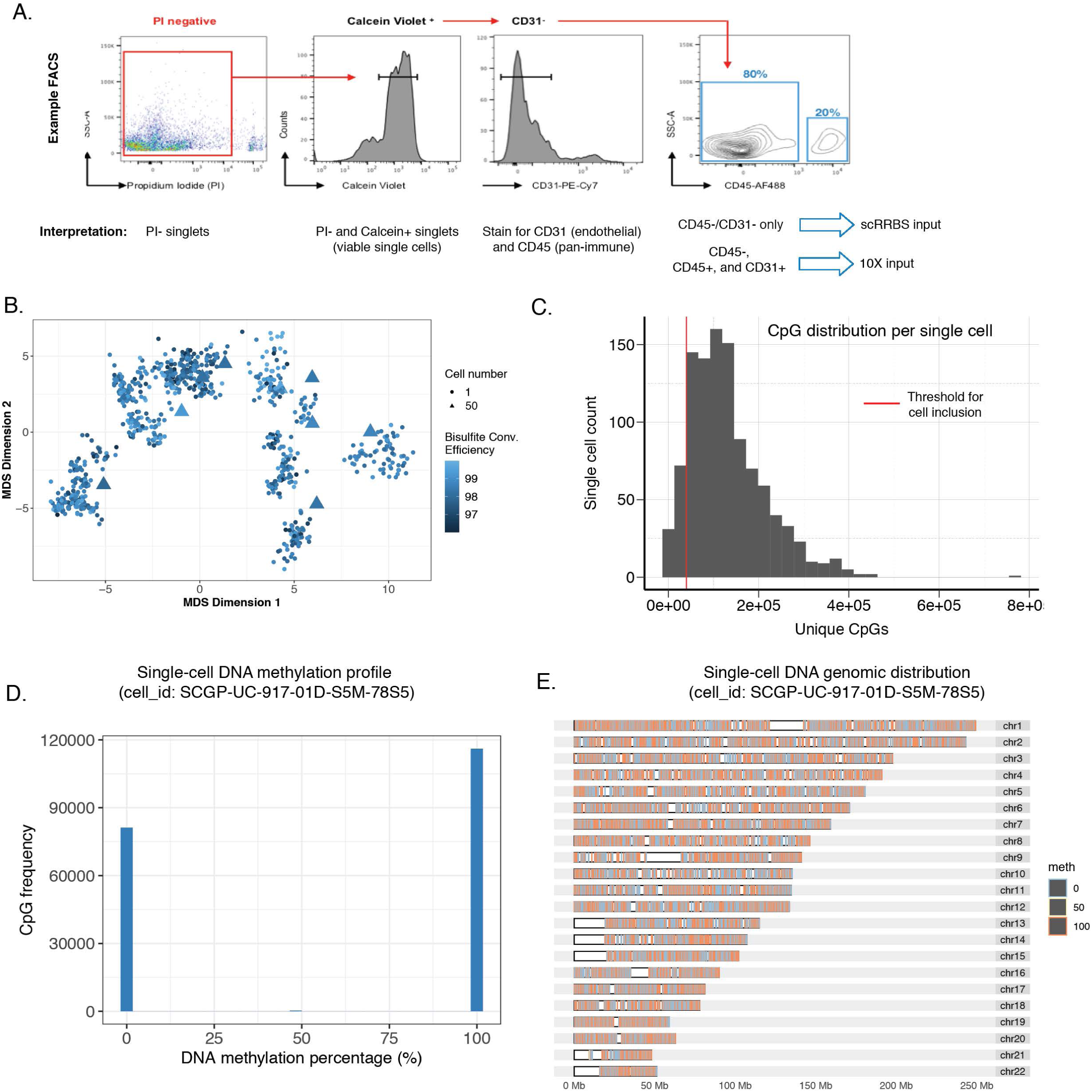
**Sample pre-processing and metrics related to single-cell DNA methylation data assessment. Related to Figure 1.** (A) Representative fluorescence activated cell sorting (FACS) data and strategy for viable cell enrichment for both single-cell protocols, and tumor cell enrichment in scRRBS. (B) The same multidimensional scaling (MDS) analysis using pairwise distance metrics calculated between individual cells as in Figure 1B, except colored by bisulfite conversion efficiency. (C) The number of unique CpGs detected per single cell, with the red line indicating the threshold (minimum 40,000 unique CpGs) for inclusion in the dataset presented herein. (D) Representative distribution of single locus DNA methylation estimates for a single cell. DNA methylation percentage of 0 represents an unmethylated locus, while a percentage of 100 represents a methylated locus. (E) Representative genomic distribution of DNA methylation values within a single cell.

**Figure S3.**
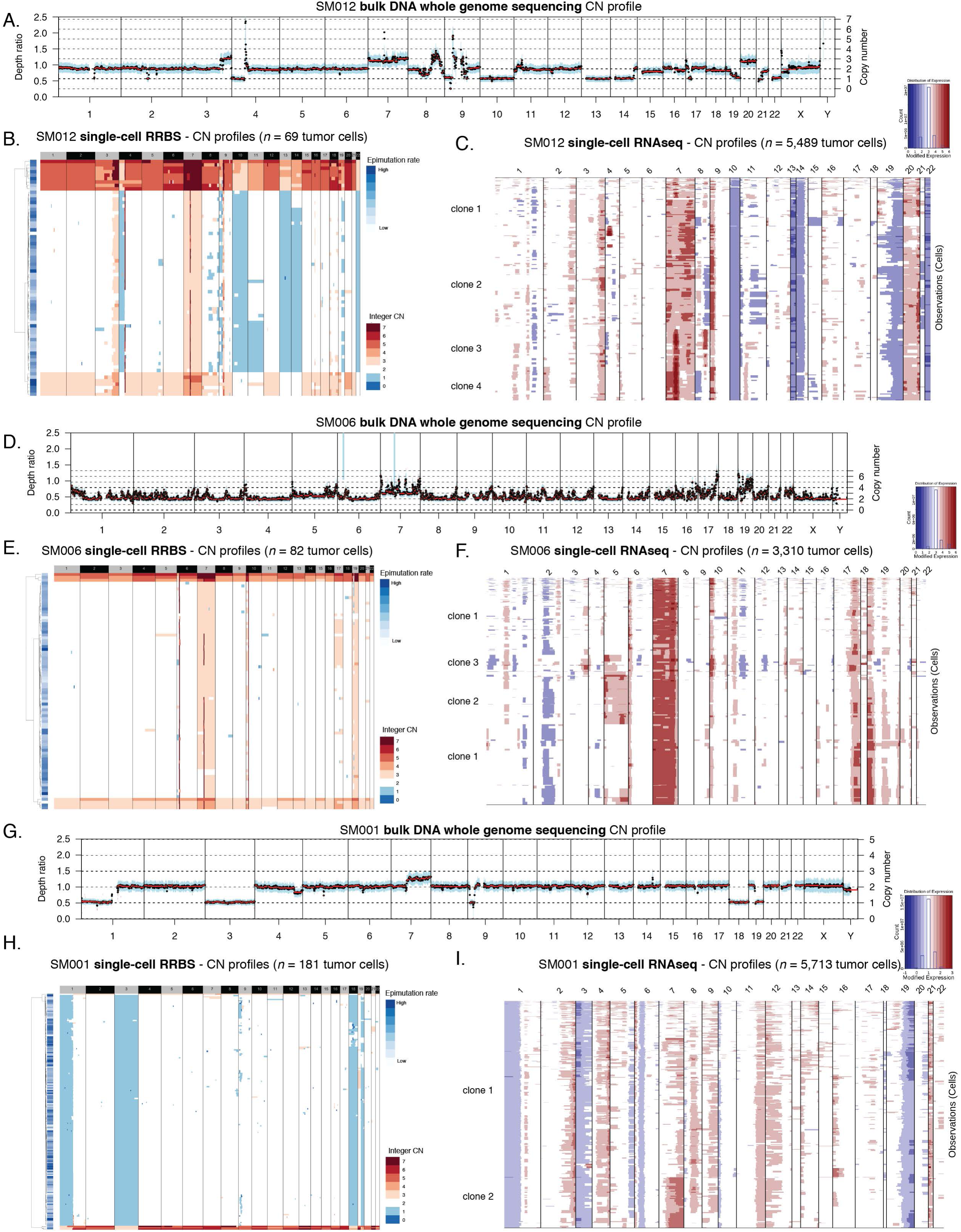
**Somatic copy number alteration examples estimated from whole genome sequencing, single-cell Reduced Representation Bisulfite Sequencing, and single-cell RNA-sequencing. Related to Figure 1.** (A-C) Representative images of copy number alterations derived from SM012 (IDHwt initial) whole genome sequencing (WGS) data. (A) Depth ratio for each segment with copy number status determined as compared with germline (normal blood) WGS data. (B) SM012 Single-cell DNA methylation-based copy number estimates (*n* = 69 tumor cells) with copy number integer depicted by color (blue = CN loss, white = neutral CN, and red = CN gain). Each row is a single cell with annotation for epimutation burden provided. (C) SM012 Single-cell RNAseq based copy number inference (*n* = 5,489) identifying major copy number events found in WGS with labelled subclones as presented in Figure 4D. (D-F) Similar example profiles as presented in (A-C) for tumor sample SM006 (IDHwt initial, *n* = 82 scRRBS cells, *n* = 3,310 scRNAseq cells). (G-I) Similar example profiles as presented in (A-C) for tumor sample SM001 (IDHmut recurrence, *n* = 181 scRRBS cells, *n* = 5,713 scRNAseq cells).

**Figure S4.**
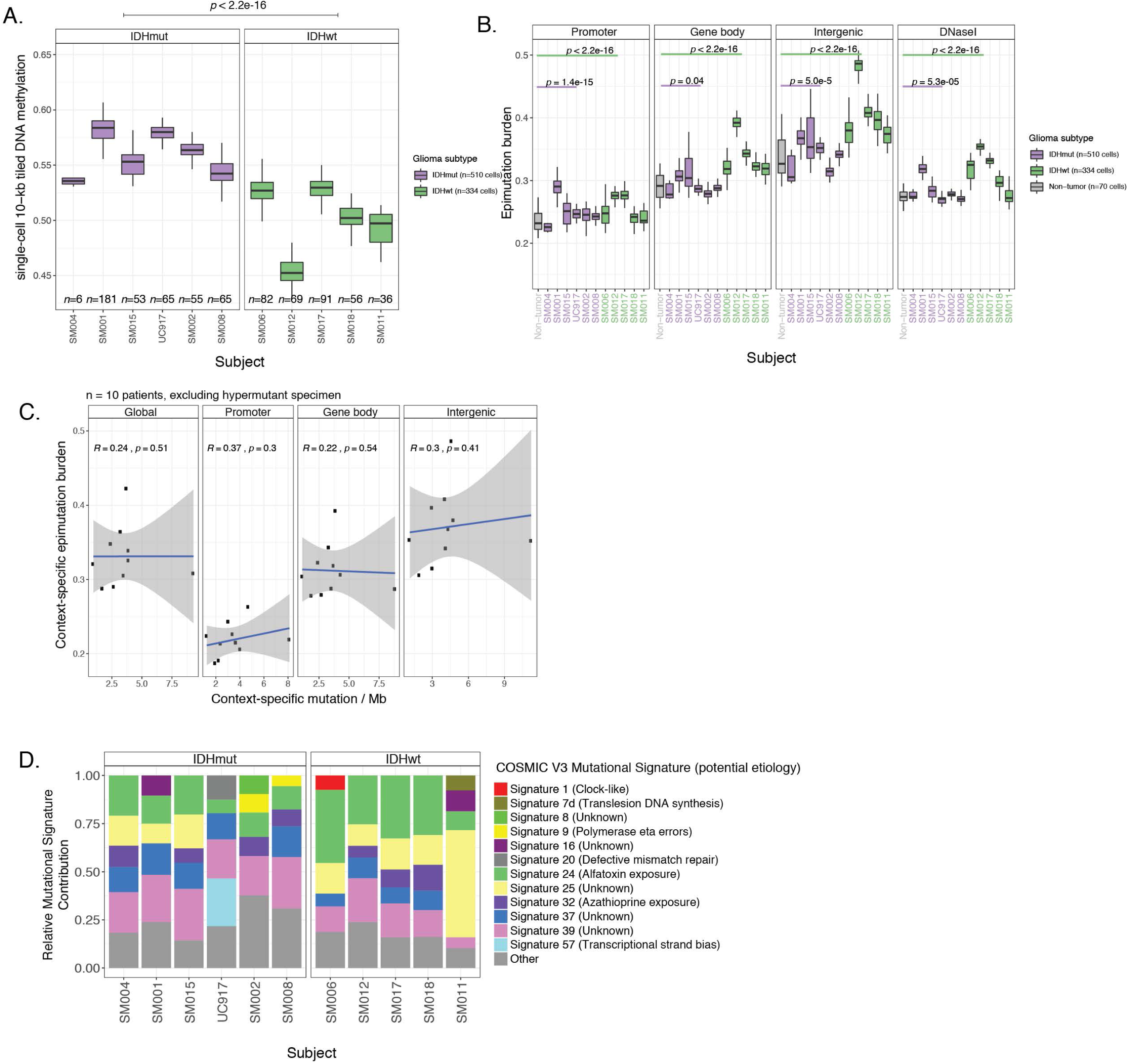
**Distribution and relationship of DNA methylation and epimutation throughout the glioma genome. Related to Figure 1.** (A) Boxplots representing average 10-kb tiled DNA methylation values per single tumor cell. (B) Boxplots highlighting the single-cell epimutation burden estimates calculated across different genomic contexts. (C) Scatterplots showing the relationship between genomic context-specific single-cell epimutation burden (sample-specific scRRBS average) and genomic context-specific mutation burden derived from whole genome sequencing (*n* = 10 excluding hypermutant sample). Panels are separated into global (i.e., all regions), promoter, gene body, and intergenic regions (Spearman correlations *p* > 0.05 for all comparisons). (D) The dominant Catalogue of Somatic Mutations in Cancer (COSMIC v3) mutational signatures are presented for each subject. The stacked bar plots represent the relative contribution of each mutational signature to the tumor’s mutational burden. Colors indicate distinct mutational signatures, which are further annotated with their proposed etiology.

**Figure S5.**
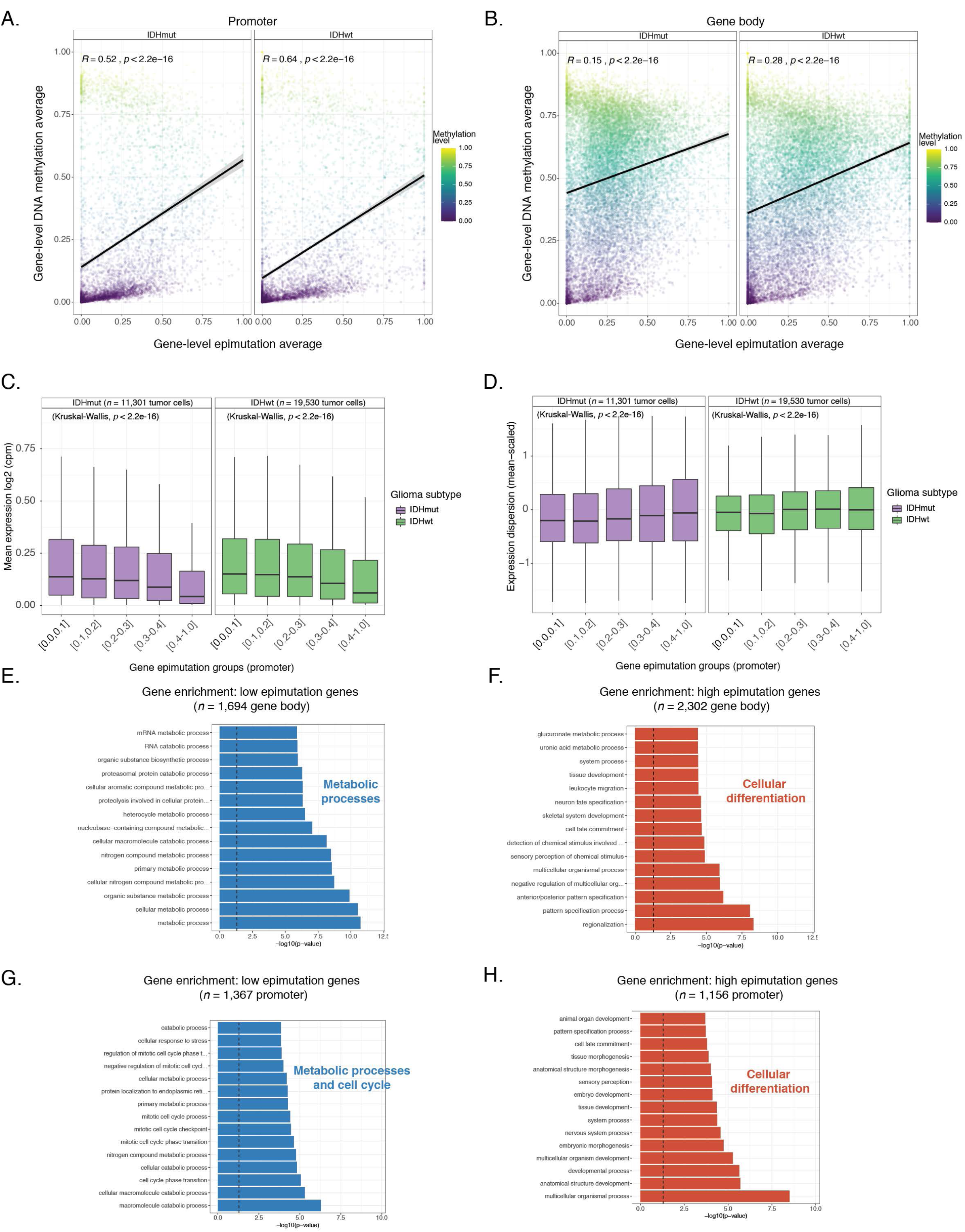
**Association between epimutation and disrupted transcriptional programs. Related to Figure 1.** (A-B) Scatterplots depicting single-cell gene-level epimutation average plotted against the gene-level methylation estimates in both (A) promoter regions and (B) gene body regions. (C) Boxplots of gene expression values, in log2 (counts per million), from single-cell RNAseq data across different sets of promoter regions defined by gene-derived epimutation groups. Gene epimutation groups are defined by the determining the mean epimutation value across a single gene. Color indicates *IDH1* mutation status. (D) Boxplots of gene expression dispersion that were mean-expression scaled to account for expression level-dependent variability across the same promoter-based gene epimutation groups defined in panel C. (E-F) Gene Ontology enrichment analyses for low epimutation genes (Figure 1E, mean epimutation across all tumor cells: 0.0 - 0.1) and high epimutation genes (Figure 1F, mean epimutation across all tumor cells: > 0.5) using gene body estimates. A meta biological process is placed next to significant Gene Ontology terms. (G-H) Same analyses presented in panels E-F, but for gene-level epimutation estimates determined in promoters.

**Figure S6.**
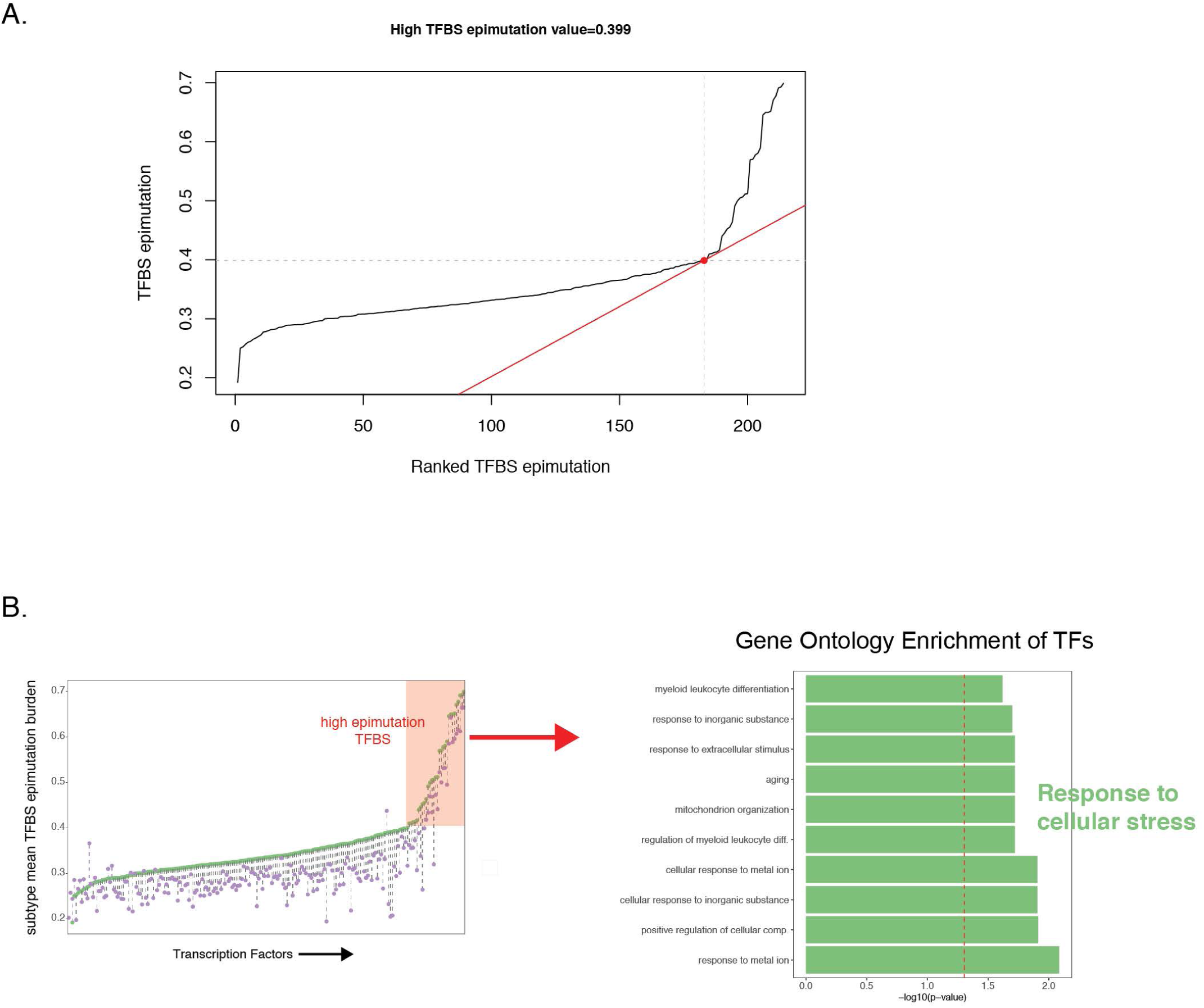
**Enrichment of high epimutation transcription factors and association with environmental stress response. Related to Figure 1.** (A) Computational approach to defining TFBSs with high epimutation burden (red tangent line at 0.399 TFBS epimutation burden). X-axis represents each TF ordered by mean epimutation burden in IDHwt single-cells (*n* = 334 cells). (B) Gene Ontology enrichment analysis of TFs with high epimutation burden in their binding sites. A meta biological process is placed next to significant Gene Ontology (GO) terms.

**Figure S7.**
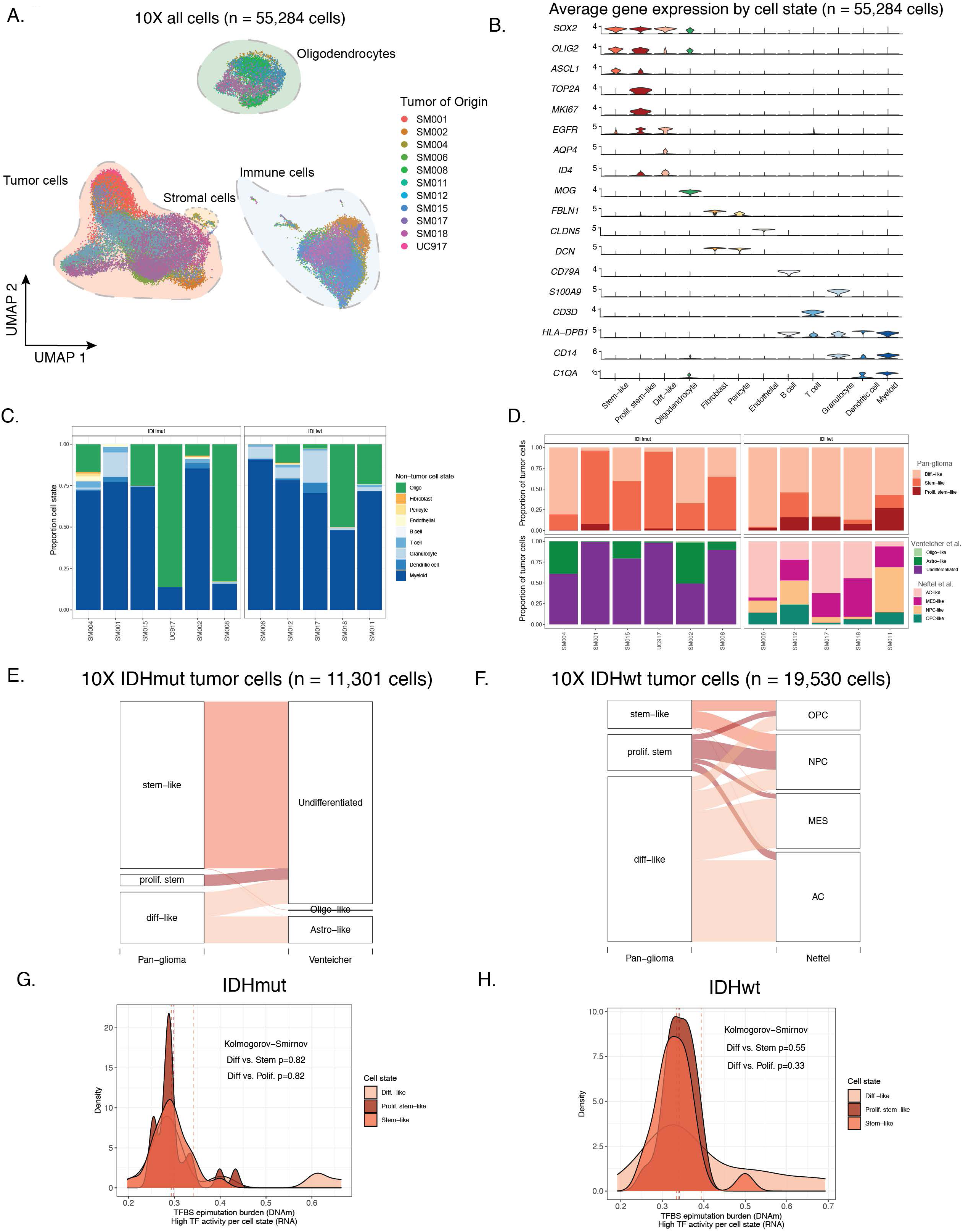
**Pan-glioma cell state assignment and characteristics. Related to Figure 2.** (A) UMAP dimensionality reduction plot of all scRNAseq data, including tumor and non-tumor cells (*n* = 55,248 cells). Each dot depicts a single cell and colors represents the tumor of origin. Shaded regions represent cell state classification. (B) Stacked violin plots of average single-cell gene expression for cells presented in Figure S7A. Selected genes presented are informative for cell state classification. (C) Stacked bar plots representing the proportion of non-tumor cellular states (D) Stacked bar plots representing the proportion of tumor cellular states per tumor for pan-glioma classification (top row) and previously published classifications (middle row; Venteicher et al. and Neftel et al.) (E) Sankey plot representing the proportion of IDHmut tumor cells with pan-glioma classification and associated classification described in Venteicher et al. (F) Sankey plot representing the proportion of IDHwt tumor cells with pan-glioma classification and associated classification described in Neftel et al. (G) Density plots representing TFBS epimutation burden (scRRBS data) in IDHmut single-cell DNA methylation data for TFs whose activity (scRNAseq based SCENIC analysis) characterizes a specific cell state (*n* = 20 TFs per cell state). Kolmogorov-Smirnov *p*-value tests for differences in TFBS epimutation burden across the cellular states. (H) Density plots representing TFBS epimutation burden (scRRBS data) in IDHwt single-cell DNA methylation data for TFs whose activity (scRNAseq based SCENIC analysis) characterizes a specific cell state (*n* = 20 TFs per cell state). Dotted lines represent the median TFBS value for cell state defining TFs. The Kolmogorov-Smirnov *p*-value corresponds to differences in TFBS epimutation burden across the cellular states.

**Figure S8.**
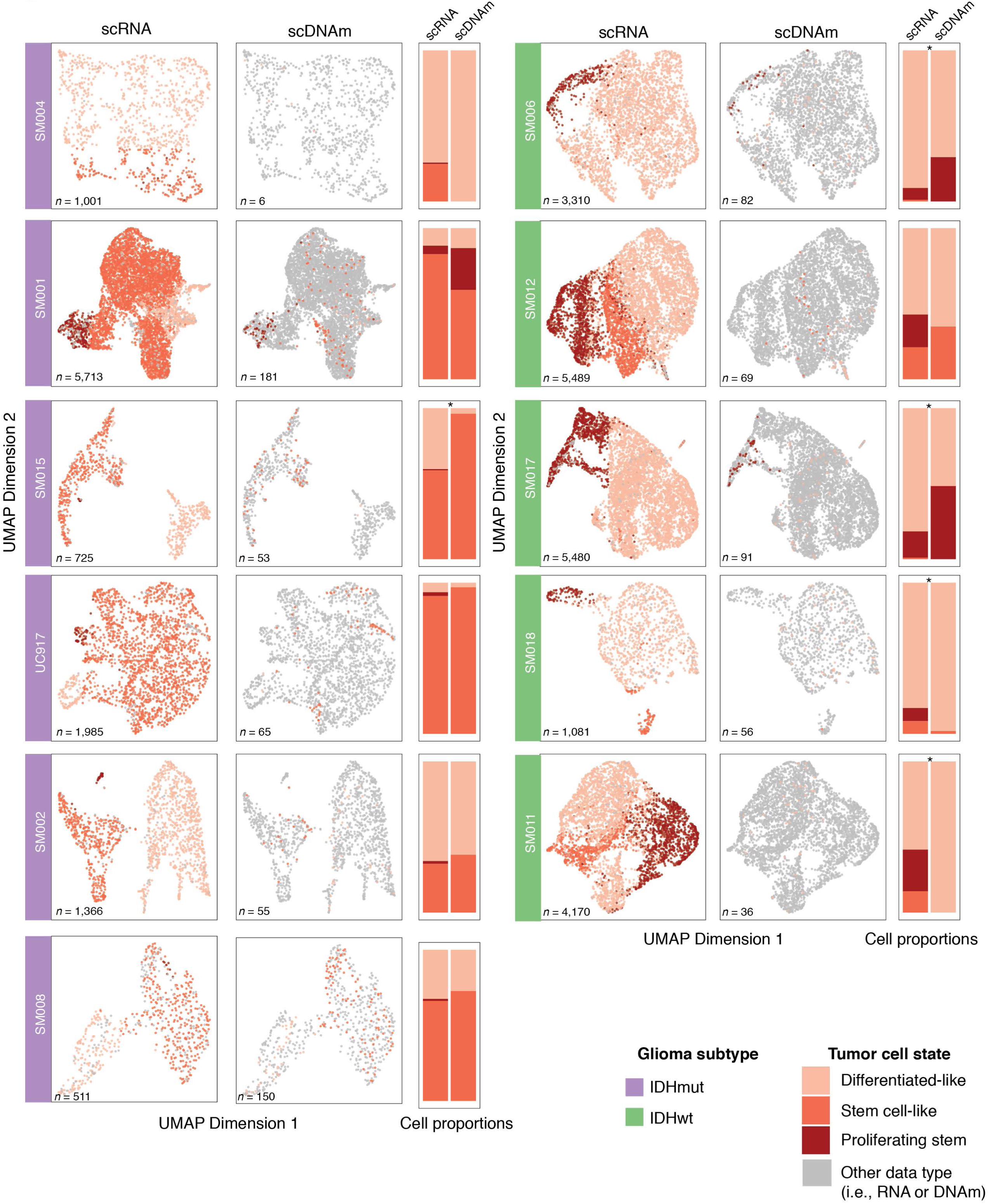
**LIGER integrated tumor-specific clustering of single-cell RNA and single-cell DNA methylation data. Related to Figure 2.** Joint single-cell RNAseq (scRNA) and single-cell DNA methylation (scDNAm) clustering and UMAP projections highlighting similar cellular state distributions across platforms. Sample annotation is presented on the left of each paired UMAP plot, each dot is an individual single cell, and cell number for each technology is presented in the lower-left hand corner. UMAP coordinate space remains the same for both scRNA and scDNAm visualizations with cellular states for that platform represented by a colored dot and data for the other platform represented by a gray dot. Stacked bar plots enumerating the proportion of cellular states detected by each platform are presented to the right of each paired UMAP plot. ‘*’ indicate specimens in which the cellular proportions across the two platforms are significantly different (Fisher’s Exact test, *p* < 0.05).

**Figure S9.**
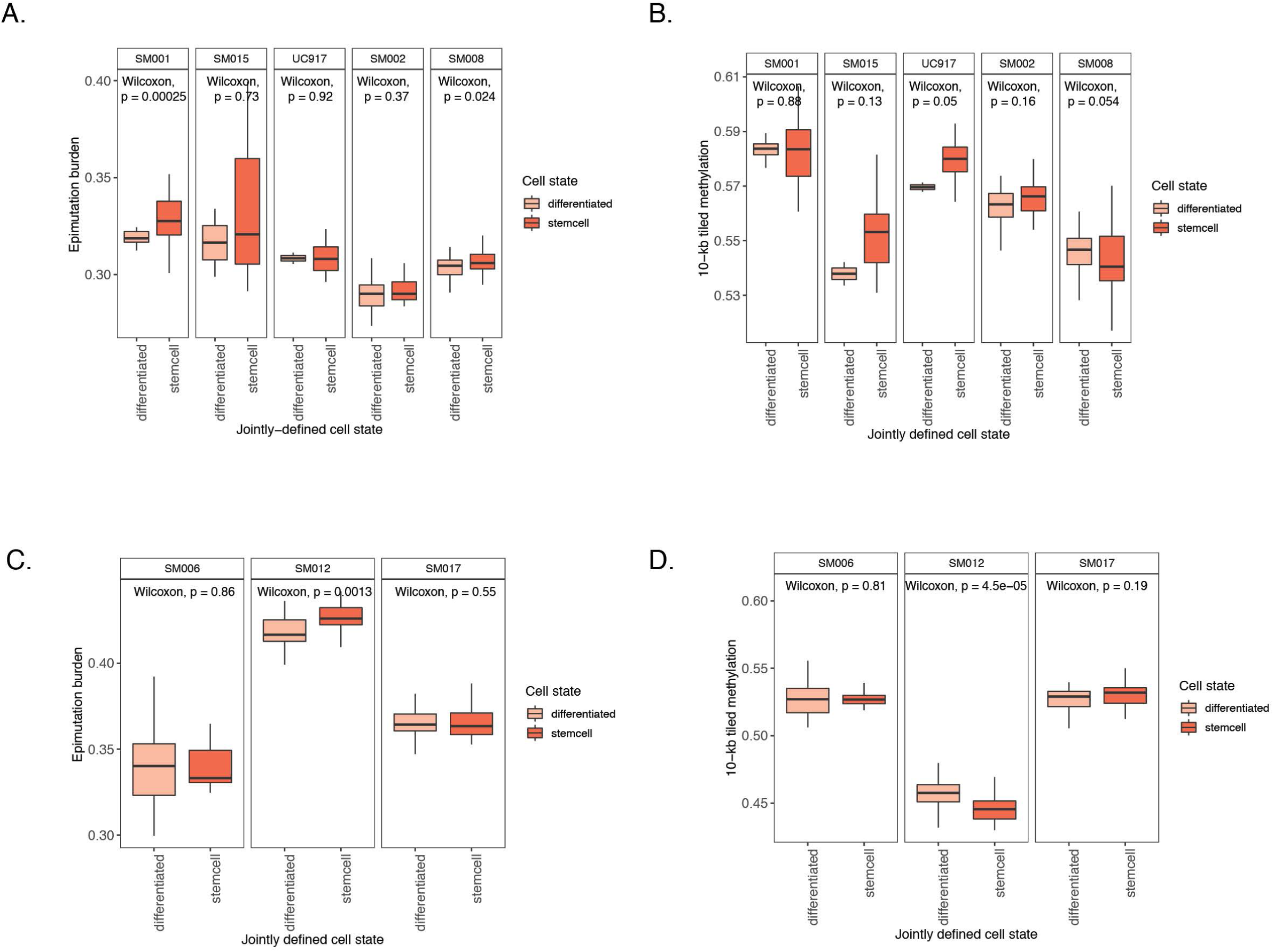
**Sample-specific differences in DNA methylation and epimutation burden across different cellular states. Related to Figure 2.** (A-B) Boxplots showing sample-specific differences in (A) epimutation burden and (B) 10-kb tiled DNA methylation across LIGER-defined cellular states in IDHmut tumors. Wilcoxon Rank Sum *p*-values are presented comparing cells from a given tumor. (C-D) Boxplots showing sample-specific differences in (C) epimutation burden and (D) 10-kb tiled DNA methylation across LIGER-defined cellular states in IDHwt tumors. Wilcoxon Rank Sum *p*-values are presented comparing cells from a given tumor. Samples with only one defined cell state are not visualized.

**Figure S10.**
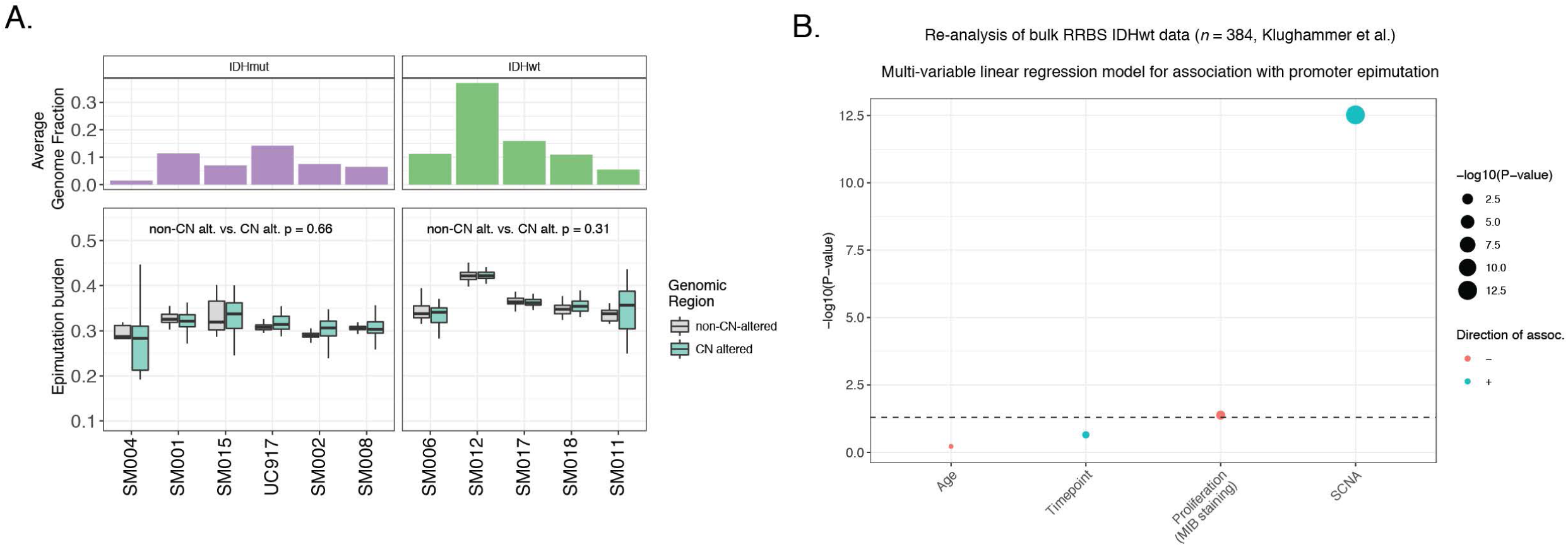
**Relationships between epimutation burden and genetic alterations. Related to Figure 3.** (A) Single-cell epimutation burden estimates were calculated across genomic regions with (teal) and without (gray) copy number alterations. The paired-sample Wilcoxon test *p*-value for each subtype represents the statistical difference of epimutation burden across these two regions. (B) Visualized results from multi-variable linear regression model testing for association with epimutation burden. Dot size indicates −log10 (*p*-value) for each predictor and color represents direction of association with epimutation burden (red = negative association, blue = positive association). Explanatory variables included subject age, timepoint (pre- and post-treatment), level of cellular proliferation determined by histological marker (MIB staining), and somatic copy number alteration burden (SCNA, total number of bases altered / total number of bases measured).

**Figure S11.**
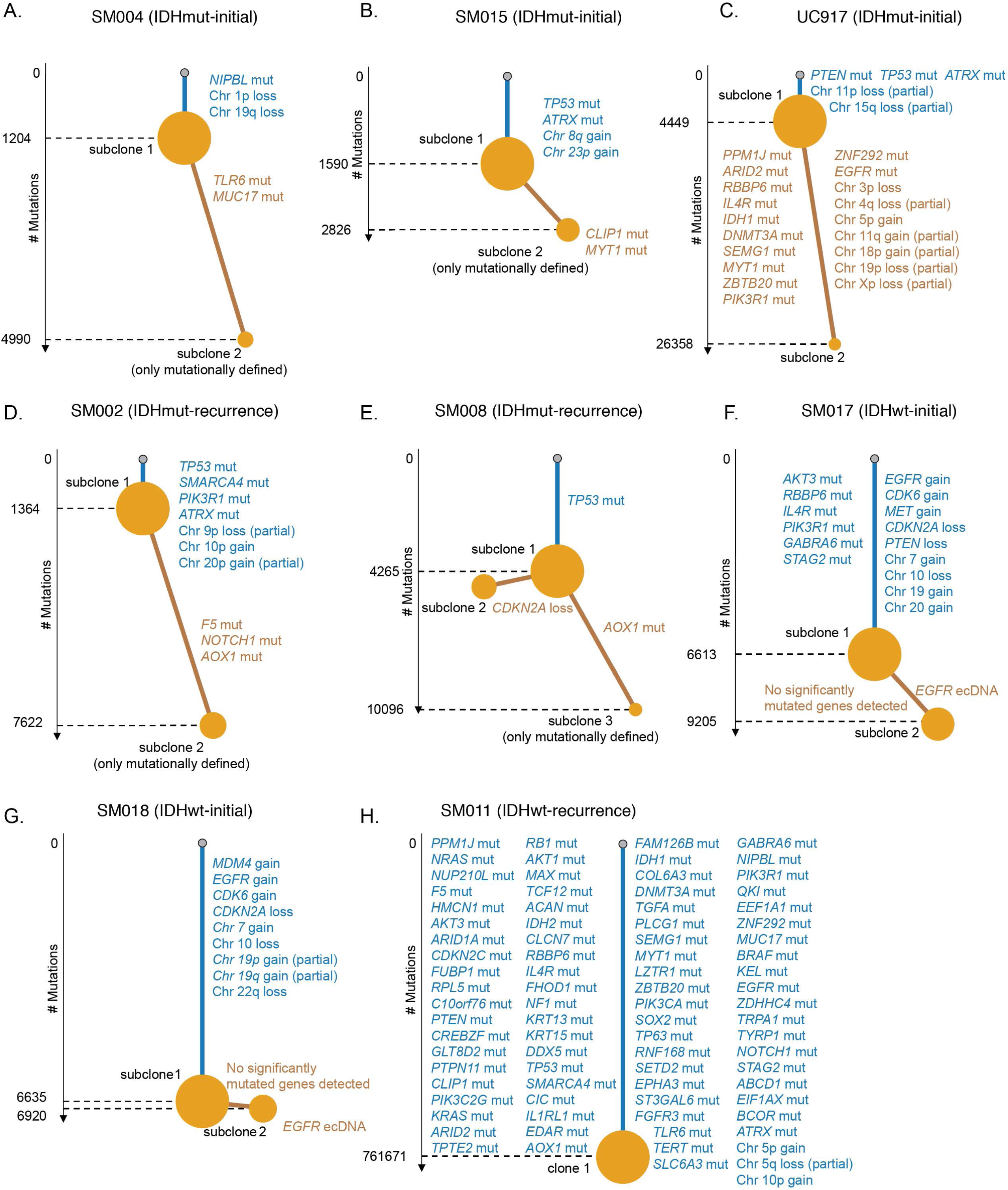
**Whole genome sequencing phylogenetic inference of tumor samples. Related to Figure 4.** (A-H) Phylogenetic trees constructed from whole genome sequencing data (mutations and somatic copy number alterations) using phyloWGS and further annotated using single-cell inferred copy number alterations (scRRBS + scRNAseq). Tree nodes represent alterations specific to the given clone, with node size corresponding to the fraction of cells with the associated alterations. Branch length scales with the number of mutations attributed to that clone. Clonal alterations are colored in blue, with subclonal alterations colored in gold. Genes considered significantly mutated in TCGA analyses (Ceccarelli et al., 2016) and chromosomal arm-level events are presented. Arm-level events are defined as spanning at least 80 percent of the chromosome arm, while partial events span at least 40 percent.

**Figure S12.**
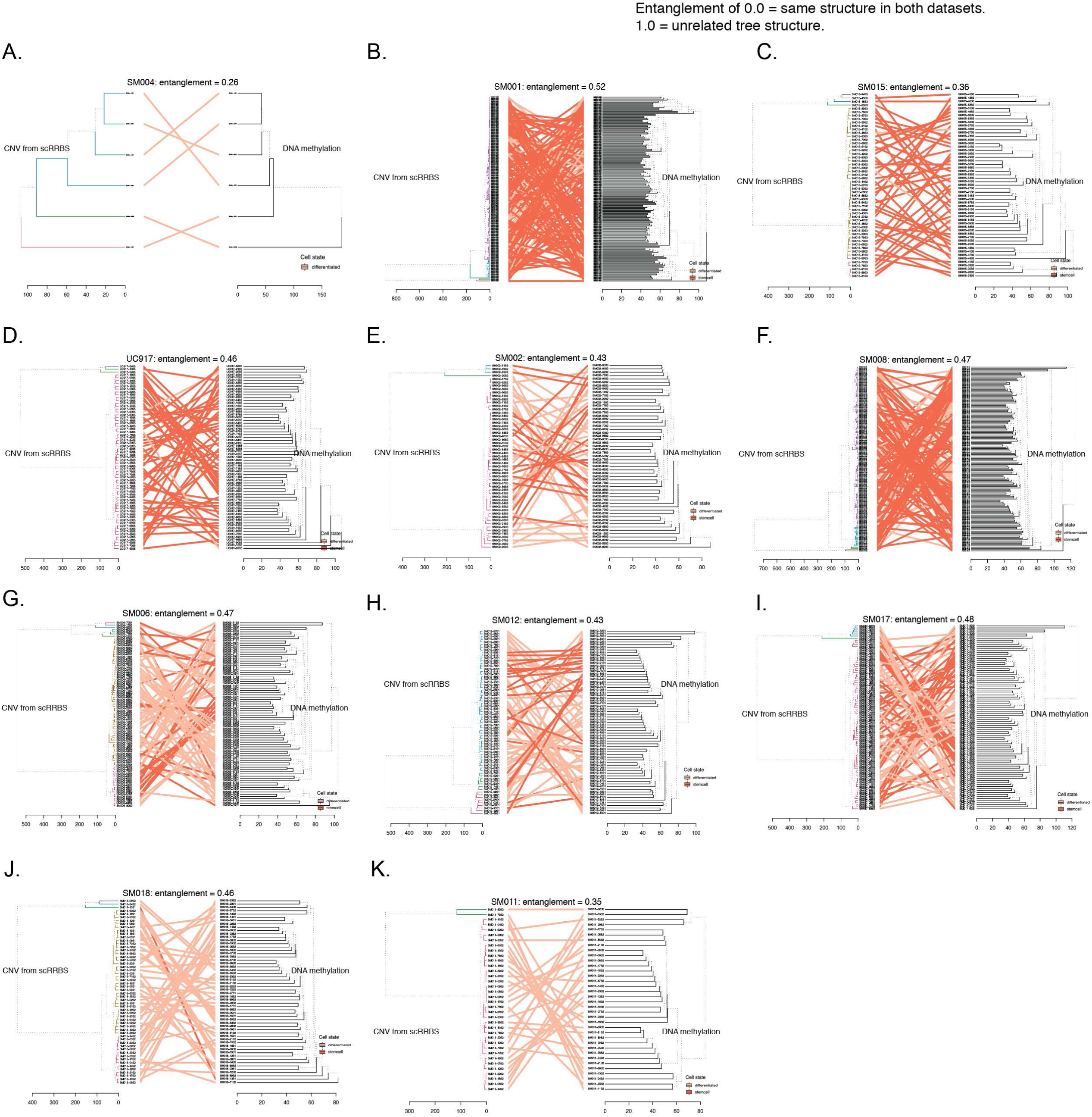
**Tumor-specific comparisons of phylogenetic and phyloepigenetic trees. Related to Figure 4.** (A-K) Tangelgrams highlighting the relationship between single-cell copy number and single-cell DNA methylation tree diagrams. Phylogenetic trees (left cluster) were calculated from copy number profiles (scRRBS data) and phyloepigenetic trees were constructed from the same cells across 10-kb tiled DNA methylation values. Cluster labels are connected with solid lines and are colored by cellular states determined by LIGER. Entanglement scores are listed above the phylogenetic and phyloepigenetic trees and indicate whether labels share the same structure (score = 0) or exhibit unrelated structures (score = 1).

**Figure S13:**
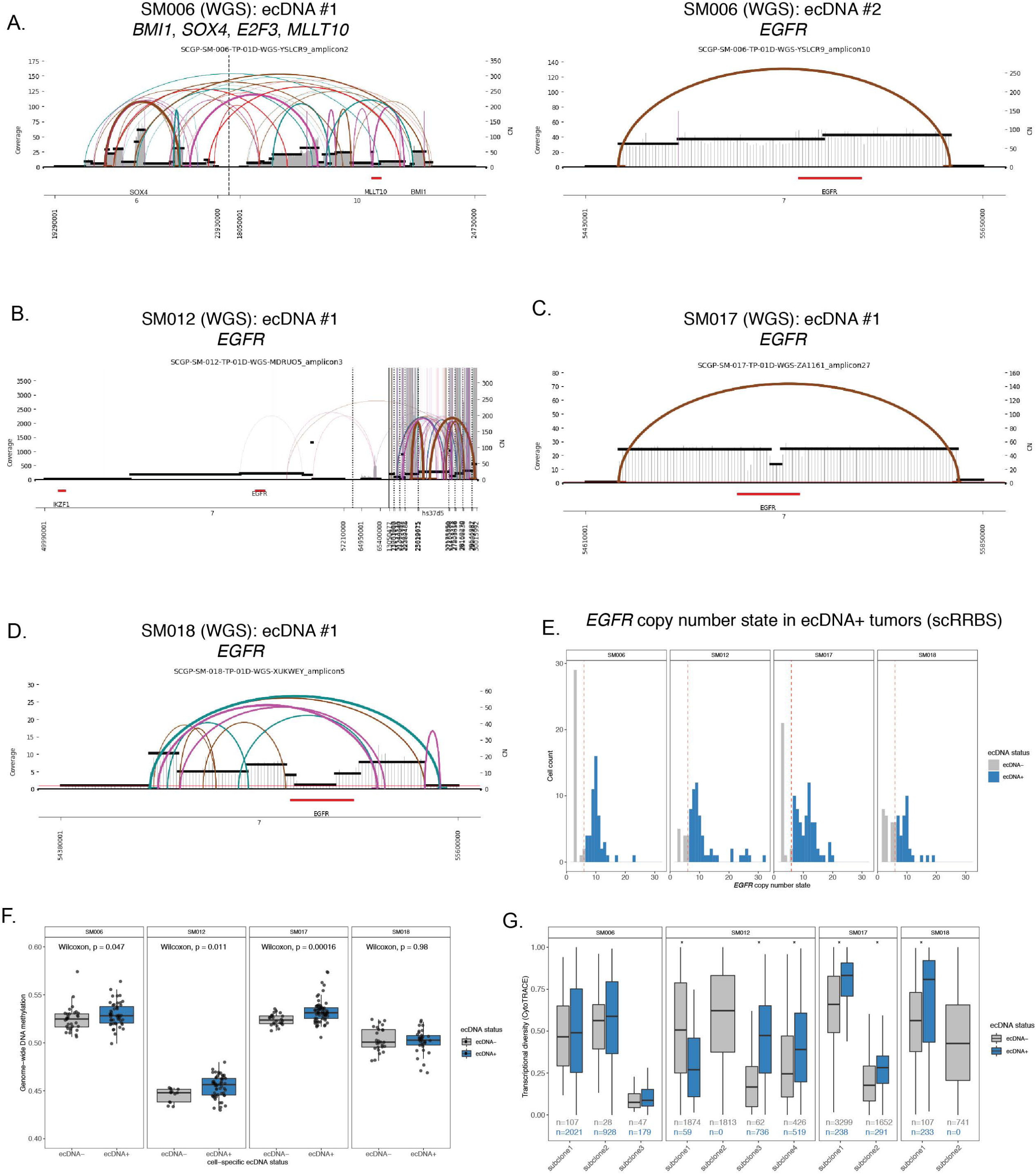
**Focal extrachromosomal DNA amplifications generate greater levels of epigenetic and transcript diversity in glioma single cells. Related to Figure 4.** (A-D) Extrachromosomal DNA circular amplicon reconstruction displaying genomic rearrangements predicted from whole genome sequencing. Coverage depth is represented as a histogram across a genomic interval with segment copy number (CN) estimation provided on the right y-axis. Discordant read pair clusters are indicated by arcs and colors highlight read pair orientation (e.g., brown = everted read pairs, (Deshpande et al., 2019). Amplicon intervals are provided at the bottom of the plot with annotation for known oncogenes (e.g., *EGFR*). (E) *EGFR* copy number estimation from single-cell RRBS data in ecDNA+ tumors. Cells with *EGFR* copy number greater than 7 were classified as *EGFR* ecDNA+ (blue). (F) Single-cell 10-kb tiled DNA methylation separated by *EGFR* ecDNA status. Single cells with inferred copy number status greater than 7 were classified as ecDNA+ (blue). Wilcoxon rank sum test *p*-values comparing DNA methylation across ecDNA status are reported for each patient tumor. (D) Boxplots depicting transcriptional diversity using gene count signatures calculated in scRNAseq data for each tumor, with cells separated based on inferred *EGFR* copy number status (gray = *EGFR* ecDNA-, blue = *EGFR* ecDNA+). Transcriptional diversity was compared based on predicted ecDNA status within each tumor subclone. Stars (*) indicate statistically significant differences based on Wilcoxon Rank Sum test (*p* < 0.05).

**Figure S14.**
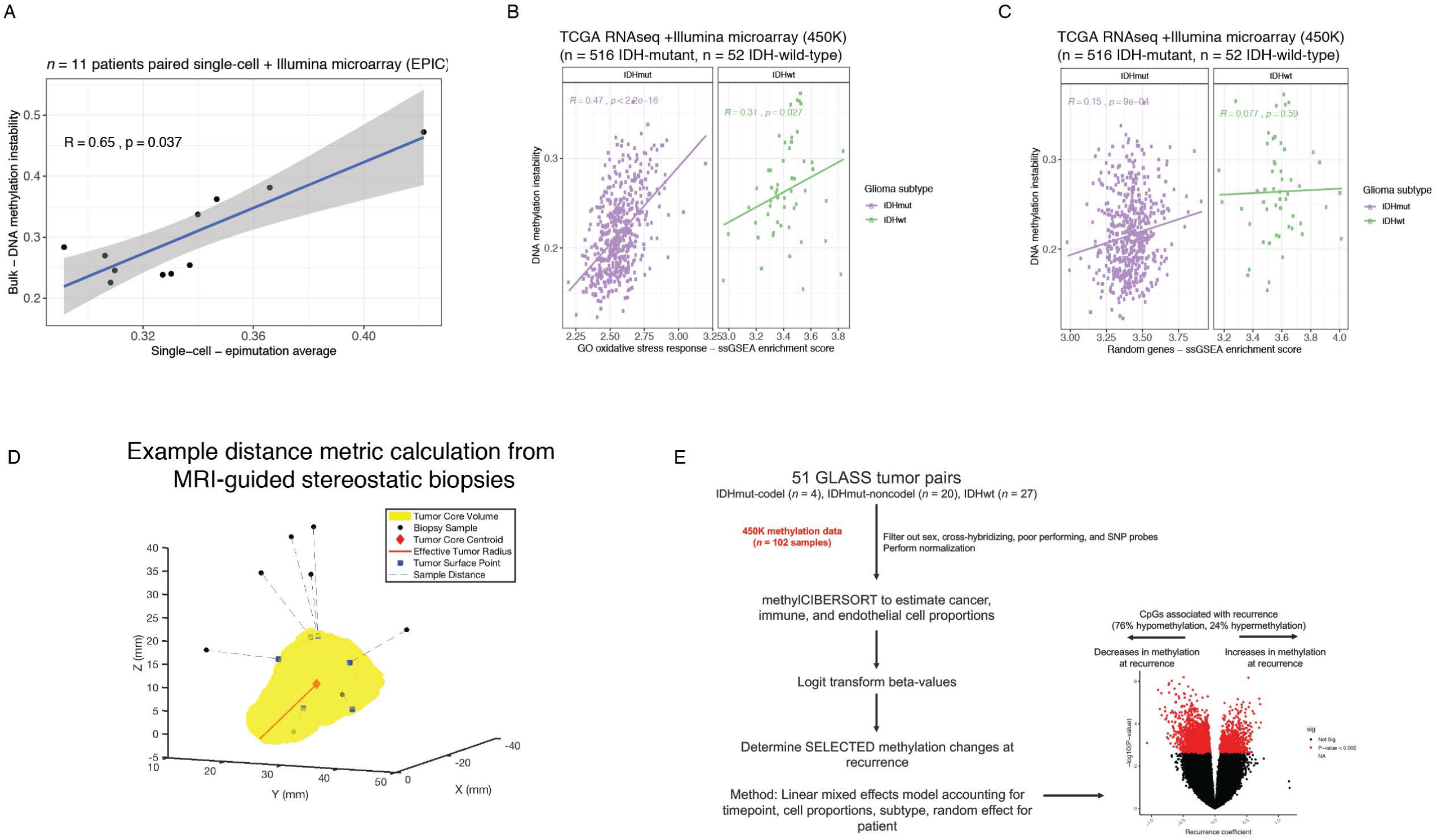
**DNA methylation instability metrics calculated in primary tumor, spatial, and longitudinal cohorts. Related to Figure 5.** (A) Scatterplot highlights the significant positive correlation between the single-cell epimutation burden metric and the bulk microarray-based DNA methylation instability metric (*n* = 11 tumors, Spearman correlation). (B-C) Scatterplot between DNA methylation instability and (bulk RNAseq) ssGSEA enrichment scores for (B) oxidative stress response genes and (C) a randomly selected gene set substantiates finding that epigenetic instability is associated with stress response. (D) Schematic depiction of magnetic resonance image-guided biopsies and radiographic features used in spatial cohort (Verburg et al.). (E) Workflow for linear-mixed effect model identifying differentially methylated CpG sites that are selected for during tumor evolution when adjusting for estimated cellular proportions, glioma subtype, and a random effect for patient (*n* = 102 tumor samples, *n* = 51 patients).

## METHODS

### EXPERIMENTAL METHODS

#### Description of human tumor specimens

Human glioma resection specimens were obtained from the University of Connecticut Health Center and from St. Michael’s Hospital. All tissue donations were approved by the Institutional Review Board of the Jackson Laboratory and clinical institutions involved. This work was performed in accordance with the Declaration of Helskinki principles. Initial pathological diagnosis was confirmed with tumor DNA methylation classification according to the Molecular Neuropathology Tool (Capper et al., 2018). Clinical characteristics for this population are provided in Table S1.

#### Sample preparation, partitioning, and fluorescence activated cell sorting for single-cell experiments

Tumor specimens were collected directly from the operating room and immediately placed into MACS tissue storage solution at 4C (Miltenyi, Cat. no. 130-100-008). Tumor specimens from the same spatial region were then minced and partitioned into single-cell and bulk fractions (Figure 1A). Any remaining tumor tissue was deposited into freezing media consisting of 90% heat-inactivated fetal bovine serum (FBS) (Invitrogen) and 10% dimethyl sulfoxide (Sigma-Aldrich), and gradually frozen in a freezing container (Mr. Frosty, Corning) over 24 hours before being stored in liquid nitrogen for future experiments (i.e., Fluorescence *in situ* hybridization). Bulk tissue specimens were immediately flash frozen for subsequent DNA and RNA extraction. The specimen fraction for single cell analyses was further mechanically and enzymatically dissociated using the Brain Tumor Dissociation Kit (P) according to the manufacturer’s protocol (Miltenyi Cat. No. 130-095-942) and as previously reported (Neftel et al., 2019; Tirosh et al., 2016; Venteicher et al., 2017).

Single cell suspensions were blocked with human BD Fc Block (BioLegend) for 5 min on ice, prior to antibody staining, and labelled via incubation with 1:100 dilution of Alexa Fluor 488 conjugated anti-CD45 antibody (Cat. no. 304017, BioLegend) and 1:100 dilution of PECy7-conjugated anti-CD31 antibody (Cat. no. 303117, BioLegend) for 30 minutes at 4C. Cells were washed with Hank’s buffered saline solution and resuspended in 2mM EDTA/ 2% BSA/ PBS buffer containing [2µg/mL] propidium iodide (PI) (BD Biosciences, Cat. No. 556364) and [1µM] Calcein violet (Invitrogen) for 20 minutes at 4C. Fluorescence activated cell sorting (FACS) was performed using a BD FACSAria Fusion instrument with an 130µm nozzle and using the lowest event rate. Single cell mode was selected to further ensure stringency of sorting. For the generation of 10X sequencing libraries, 50,000-150,000 PI^-^, Calcein+ viable single cells were collected in 20% FBS/HBSS buffer. CD45+ cells were limited to no more than 20% of the total viable sort to enrich for tumor cells (Figure S2A). For the generation of single-cell DNA methylation libraries, we sorted viable (PI^-^ and Calcein+), non-immune (CD45^-^), and non-endothelial (CD31^-^) cells into 96-well plates that were pre-loaded with 5 µL of 1X Tris-EDTA buffer (Figure S2A). Once the cells had been sorted, 96-well plates were either immediately processed through the single-cell DNA methylation protocol or flash frozen and stored at −80C.

#### scRRBS library preparation

Single-cell DNA methylation profiling was performed using a modified version of a previous scRRBS protocol (Guo et al., 2015; Guo et al., 2013). Single-cell DNA methylation experiments were performed with sorted viable, non-immune, non-endothelial (PI-, Calcein+, CD45^-^, CD31^-^) cells in a 96-well plate containing 5 µL pre-loaded Tris-EDTA buffer with an empty well control. For 9 out of 11 tumors, the protocol was also applied to a small population control of 50-cells (PI-, Calcein+, CD45^-^, CD31^-^). Sorted 96-well plates were frozen at −80 C until processing when cells were lysed with 0.2 µL 1 M KCl (Millipore Sigma), 0.2uL 10% Triton X-100 (Millipore Sigma), 0.3 µL 20mg/mL protease (Qiagen), and nuclease-free water in a total volume of 6 µL for 3 hours at 50 C. The protease was then heat-inactivated at 75 C for 15 minutes. The DNA was incubated with 50 units of MspI (NEB) and TaqI (NEB) with CutSmart buffer (NEB) for 3 hours at 37 C. 60fg of unmethylated Lambda bacteriophage DNA (Promega) was added to each well to serve as a control for bisulfite conversion efficiency assessment. The solution was heated to 80 C for 20 minutes to heat-inactivate the restriction enzymes and placed on ice. 5 units of Klenow Fragment (3’à 5’ exo-, NEB), CutSmart buffer (NEB), and end-repair dNTP mix (40uM dATP, 4uM dGTP, and 4uM dCTP; NEB) totaling 2 µL per reaction were added to perform end-repair and dA-tailing. 1:250X diluted NEXTflex methylated adapters (BiooScientific) were added to each quadrant of the 96-well plate (*n* = 24 unique adapters) with a ligation mixture of 40 Weiss U T4 ligase (NEB), 1mM ATP (ThermoFisher Scientific), and nuclease-free water to a final volume of 4 µL per reaction. TruSeq methylated adapters (Illumina) were also used in a single sample (SM001) using the same protocol. The ligation reaction proceeded at 16 C for 30 minutes followed by an incubation of 4 C for at least 8 hours. The ligation reaction was stopped by heat-inactivation at 65 C for 20 minutes. Post-adapter ligation, 24 individual cells with unique ligated adapters were pooled from each plate quadrant for the protocol’s remainder. Excess adapter was removed using a 1:1 volumetric ratio of Ampure beads (Beckman Coulter). Bisulfite conversion was performed using the EZ-DNA methylation kit (Zymo) according to the manufacturer’s instructions except with one-half volumes due to reduced DNA input. The solution was incubated at 98 C for 8 minutes, 64 C for 3.5 hours, and held at 4 C once the reaction was complete. 10ng of tRNA (Roche) was added prior to column elution to serve as a protective carrier. PCR enrichment was performed using the PfuTurbo Cx hotstart (Agilent), PfuTurbo Cx hotstart buffer (Agilent), primer mix (Bioo Scientific), dNTP mix (Promega), and nuclease-free water under the following conditions: 95 degrees Celsius for 2 minutes, 32 cycles of 95 C for 20 seconds, 60 C for 30 seconds, and 72 C for 60 seconds. The PCR reaction was terminated by incubating at 72 C for 5 minutes. The libraries were purified in a 1:1 volumetric ratio of Ampure beads (Beckman Coulter), Pippin size selection was performed between 200-1000bp (Sage Science), and quantified by qPCR (Kapa Biosystems / Roche). scRRBS libraries were paired-end sequenced alongside bulk whole genome libraries on an Illumina HiSeq4000 using 1% PhiX spike-in and 75bp reads.

#### Single-cell RNA library preparation

Sorted cells were washed and resuspended in 0.04% BSA/PBS buffer. Cells were counted on a Countess II automated cell counter and were loaded on a 10X Chromium chip with a target cell recovery of 6,000 cells per lane. Sequencing libraries were performed using the single-cell 3’ mRNA kit according to the manufacturer’s protocol (10X Genomics). cDNA and library quality were examined on an Agilent 4200 TapeStation and quantified by qPCR (Kapa Biosystems / Roche). Illumina sequencing was performed using a paired-end 100bp protocol. Libraries were sequenced to an average depth of 50,000 unique reads per cell.

#### Whole genome sequencing of tumors and matched normal blood

Genomic DNA was extracted from the same tumor region as the single-cell analyses using the Qiagen AllPrep kit and matched normal blood using DNeasy kit (Qiagen). Briefly, 400ng of DNA was sheared to 400bp using a LE220 focused-ultrasonicator (Covaris) and size selected using SPRI beads (Beckman Coulter). The fragments were treated with end-repair, A-tailing, and ligation of unique adapters (Illumina) using the KAPA HyperPrep Kit (Roche). This was followed by 5 cycles of PCR amplification when necessary. DNA sequencing was performed using paired-end 75bp protocol according to the manufacturer’s protocol (Illumina). The tumor samples were sequenced to an average depth of 44X and tumor-matched normal blood to 30X.

#### Bulk Illumina EPIC DNA methylation microarrays

250 ng of genomic tumor DNA was subject to bisulfite conversion using the EZ DNA Methylation kit (Zymo) and genome-wide DNA methylation was assessed by the Infinium MethylationEPIC kit according to the manufacturer’s protocol (Illumina).

#### Bulk RNA sequencing

Bulk tumor RNA was extracted from samples with sufficient tissue using the AllPrep kit (Qiagen). Samples with RIN values > 5 as assessed by TapeStation (Agilent Technologies) were prepared with KAPA mRNA HyperPrep kit (Roche). Libraries were sequenced using a paired-end 150bp protocol on a NovaSeq to 50 million reads according to the manufacturer’s protocol (Illumina).

#### Fluorescent in situ hybridization (FISH) analysis

Tissue slides were prepared by tumor touch prep method (deCarvalho et al., 2018). Positively charged glass slides were pressed against the surface of slightly thawed tissues. The slides were then immediately fixed by cold Carnoy’s fixative (3:1 methanol:glacial acetic acid, v/v) for 30 min and then air-dried. Slides were denatured in Hybridization buffer (Empire Genomics) mixed with EGFR-Chr7 probe (EGFR-CHR07-20-ORGR, Empire Genomics) at 75°C for 5 min and then incubated at 37°C overnight. The posthybridization wash was with 0.4x SSC at 75°C for 3 min followed by a second wash with 2x SSC/0.05% Tween20 for 1 min. The slides were then briefly rinsed by water and air dried. The VECTASHIELD mounting medium with DAPI (Vector Laboratories) was applied and the coverslip was mounted onto a glass slide. Tissue images were scanned under Leica STED 3X/DLS Confocal with 100x magnification.

### ANALYTICAL METHODS

#### Single-cell DNA methylation processing

Raw sequencing reads were trimmed to remove adapters and low-quality bases using trim_galore with the ‘--rrbs’ and ‘--paired’ parameters (version 0.4.0 https://github.com/FelixKrueger/TrimGalore). The trimmed reads were then aligned to the GRCh37 (hg19) genome using Bismark (version 0.19.1) with parameters ‘-N 1 -- bowtie2 -- score_min L,0,-0.4’ (Krueger and Andrews, 2011). PCR duplicates were removed with the ‘deduplicate_bismark’ command. Bisulfite conversion efficiency was determined using the spike-in unmethylated lambda DNA. Cells with fewer than 40,000 unique CpGs detected and bisulfite conversion rates below 95% were removed from analysis. 914 single cells were retained for downstream analysis (*n* = 914 / 1,076 total cells sequenced) with a mean of 145,000 CpGs per cell and mean bisulfite conversion rate of 98.4% (Table S4).

#### Unsupervised clustering of scRRBS data

Unsupervised clustering of the DNA methylation data was performed using pairwise comparisons of individual CpGs across all cell-to-cell comparisons as previously described (PDclust) (Hui et al., 2018). Briefly, this method performs pairwise comparisons of single-CpG methylation measurements to create a pairwise dissimilarity (PD) value that reflects the average absolute difference in methylation values at CpGs covered in any two cells. The pairwise dissimilarity values were used as input features for the Multidimensional Scaling (MDS) analysis for which visualization of cells in close proximity reflects greater similarity than cells further apart (Figure 1B).

#### Epimutation burden as a measure of epigenome instability

Epimutation burden was determined by identifying DNA methylation concordance of nearby CpGs on a single sequencing read as previously described for bulk and single-cell DNA methylation data (Gaiti et al., 2019; Landan et al., 2012; Landau et al., 2014). Briefly, in order for a sequencing read to be considered for this analysis it required a minimum of 4 CpGs located on the same sequencing read. These sequencing reads are referred to as “epialleles” and represent a subset of a cell’s total sequencing reads. An epiallele read is considered discordant if any of the CpGs on that sequencing read have different methylation states (e.g., three methylated CpGs and an unmethylated CpG). The epimutation burden metric reflects the sum of discordant epialleles divided by the total number of epialleles considered for analysis as previously described (Gaiti et al., 2019; Landan et al., 2012; Landau et al., 2014). The epimutation burden metric can be calculated across the entire genome (i.e., “epimutation burden”) or restricted to specific genomic regions where the metric considers only the epialleles overlapping that particular genomic context. A linear regression model was used to assess the impact of the total number of epialleles considered for analysis on the epimutation burden. The epimutation metric was very weakly associated with epiallele count in that an additional 10,000 epialleles was associated with an 0.001 increase in the epimutation burden metric. For analyses associating epimutation burden with metrics derived from bulk WGS data, sample-level epimutation burden was calculated as the median of single-cell epimutation values.

#### DNA methylation and epimutation over genomic annotations

To determine region-specific DNA methylation or epimutation burden, each cell’s measured CpGs or epialleles were intersected with the genomic coordinates of interest before methylation value or epimutation burden calculation, respectively. All coordinates were mapped against the hg19 human genome assembly. Regions of interest considered for analyses included promoter, gene body, intergenic, and DNaseI regions, TF binding sites, replication timing domains, and 5kb and 10kb tiled regions. Promoters were defined as 1kb upstream and 500 bp downstream of FANTOM5 (Forrest et al., 2014) TSS that mapped to Ensembl genes. If multiple TSSs mapped to a given gene, the TSS with the lowest genomic coordinate was selected. Gene body annotations were obtained from the Ensembl Genome Browser (Hunt et al., 2018). Intergenic regions were annotated by selecting regions not overlapping Ensembl gene body coordinates. DNaseI hypersensitivity region annotations were obtained from the UCSC Genome Browser (Raney et al., 2013). TF binding sites were obtained from the JASPAR 2020 Core Vertebrate database (Fornes et al., 2020) of non-redundant TF binding motifs. Each binding site is assigned a score of 0-1000, which corresponds to the p-value for the relative position weight matrix score of a TF binding site prediction. For a given TF, all identified target binding site coordinates were aggregated, and binding sites were excluded if they had a relative score less than 400, corresponding to a p-value greater than 0.0001. Replication timing of genes was retrieved from MutSigCV (Lawrence et al., 2013), and annotations for replication timing domains were generated by binning gene coordinates into quartiles based on the replication timing score. Methylation values were also calculated for non-overlapping windows of 5kb or 10kb. Ranks of high epimutation levels were determined by applying the ROSE software (https://bitbucket.org/young_computation/rose) for both gene-level and transcription factor binding sites.

#### SCNA estimation from single-cell DNA methylation data

To provide evidence for somatic copy number alterations in single-cell DNA methylation sequencing data, the Gingko algorithm (Garvin et al., 2015) was applied to single cells that passed the scRRBS quality control filters mentioned above. Briefly, this method bins mapped reads by chromosomal location, performs Lowess normalization to correct for GC biases, adjusts for potential amplification artifacts, and segments the genome to determine chromosomal regions with consistent copy number states. Here, the genome for each sample was divided into 2,597 variable-length bins with a median length of 1Mb. Segmentation was performed using independent normalized read counts and the parameter ‘mask bad bins’ (i.e., bins with consistent pileups) was enabled. Cells were considered “non-tumor” if less than 1% of the genome had a copy number state that was not 2. Copy number plots were generated using the R package “gplots”. Phylogenetic and phyloepigenetic trees were constructed for the same cells (scRRBS data) using Euclidean distance between profiles and clustered with the R function hclust using “ward.D2” linkage. The concordance between these two trees for each sample was determined using the tanglegram function in the dendextend R package and 10 random tree rotations were used to minimize artificial branch crossing (Galili, 2015).

#### Single-cell RNA processing and analysis

The Cell Ranger pipeline (v3.0.2) was used to convert Illumina base call files to FASTQ files and align FASTQs to hg19 10X reference genome (version 1.2.0). Preprocessing was performed using the Scanpy package (1.3.7) (Wolf et al., 2018). The gene expression profiles of each cell at the 1500 most highly variable genes (as measured by dispersion (Satija et al., 2015)) were used for neighborhood graph generation (using 33 nearest neighbors) and dimensionality reduction with UMAP (Becht et al., 2018). Clustering was performed on this neighborhood graph using the Leiden community detection algorithm (Traag et al., 2019). The neighborhood graph was batch-corrected using the batch correction software BBKNN (Polanski et al., 2020). These defined clusters were then labelled with particular cell states based on marker gene expression and previously described cell states (Bhaduri et al., 2020; Neftel et al., 2019; Tirosh et al., 2016). Cell state classification of malignant cells was also performed using previously developed classifiers for both IDH-wild-type (Neftel et al., 2019) and IDH-mutant tumors (Venteicher et al., 2017). The Seurat R package was also used for downstream analyses and visualizations (Stuart et al., 2019). Inference of gene regulatory networks was performed using SCENIC for a random set of 5,000 cells per subtype to permit heatmap visualization (Aibar et al., 2017). SCNA estimation from single-cell RNAseq data was performed as previously reported (Neftel et al., 2019; Tirosh et al., 2016; Venteicher et al., 2017). Briefly, the InferCNV method provides evidence for large-scale somatic copy number alterations by comparing averaged gene expression intensity values compared with normal cells (based on marker gene expression) from the same specimen. Subclusters of cells were partitioned into clones on the basis of shared copy number patterns (https://github.com/broadinstitute/inferCNV). Single-cell RNA diversity comparisons using gene count signatures were performed using the R package CytoTRACE across cells from the same tumor clone (Gulati et al., 2020).

#### Joint scRNA and scDNA methylation integration

Single-cell RNA and DNA methylation malignant cell profiles were integrated within the same specimen based on the differentially expressed across the pan-glioma RNA cell states (Table S2). The single-cell RNA data were jointly clustered with the gene-level methylation features as previously reported (Welch et al., 2019) using the R package liger (linked inference of genomic experimental relationships).

#### Analysis of publicly available brain tumor DNA methylation data

Data re-analysis of longitudinal glioma resources was accessed for Klughammer et al. (http://www.medical-epigenomics.org/papers/GBMatch/) (Klughammer et al., 2018) and the Glioma Longitudinal AnalySiS consortium (GLASS, http://synapse.org/glass) (Barthel et al., 2019). Magnetic Resonance Imaging guided biopsies taken from spatially distinct regions and subjected to bulk DNA methylation Illumina methylation microarray collected by our group was also accessed (Verburg et al.). DNA methylation microarrays (450K) were also retrieved The Cancer Genome Atlas initial glioma samples (Ceccarelli et al., 2016). All Illumina methylation microarrays were processed using the R package minfi. The recurrent DNA methylation changes between the initial and recurrent tumors were determined by fitting a linear mixed effect model (R nlme package) to each individual CpG modeled as a logit transformed M-value with independent variables of timepoint, subtype, cancer cell proportion, immune proportion, and subject included as the random effect. The cancer and immune cell proportions in the GLASS bulk Illumina methylation microarray data were determined using the glioma signature in the R package MethylCIBERSORT as previously described (Chakravarthy et al., 2018).

#### Gene and genomic region enrichment analyses

Enrichment of genes were performed using the R package topGO. Enrichment of genomic regions were determined using the Locus Overlap Analysis (LOLA) R package (Sheffield and Bock, 2016). LOLA enrichment analyses used all features considered for analysis as the background sets.

#### Variant detection and copy number calling

Variant detection and bulk copy number determination was performed in accordance to the GATK Best practices using GATK 4.1.0.0 (Mutect2) and as previously described (Barthel et al., 2019).

#### Mutational signature identification

Mutational signatures were identified in bulk WGS samples using the MutationalPatterns R package (Blokzijl et al., 2018). The trinucleotide context of single base substitutions was extracted for each sample in order to construct a mutational profile. For each mutational profile, the contribution of mutational signatures from the Catalogue of Somatic Mutations in Cancer (COSMIC v3) was quantified. Known signatures were ranked by order of relative contribution to the sample mutational profile; for visualization the top 5 signatures per sample were listed, with the remaining signatures collapsed into an “Other” category.

#### Phylogenetic reconstruction copy number / mutation clonality

To reconstruct the evolutionary history and subclonal composition of tumors, PhyloWGS (Deshwar et al., 2015) was applied to bulk WGS samples. PhyloWGS incorporates SCNAs with simple somatic mutations (SSMs) in inferred phylogenies by converting SCNAs into pseudo SSMs prior to subclonal reconstruction. For input, phyloWGS requires VCF format variant calls, SCNA segments, and estimates of tumor purity, which were generated using Mutect2 (Cibulskis et al., 2013), TITAN (Ha et al., 2014), and Sequenza (Favero et al., 2015), respectively. If a tumor contained more than 5000 variants, input variants were subsampled to 5000, ensuring all variants overlapping previously identified significantly mutated genes were included (Barthel et al., 2019; Ceccarelli et al., 2016). For each phyloWGS run, multiple Markov chain Monte Carlo chains were initiated with differing start values, and the optimum solution was selected based on negative normalized log likelihood. Cancer cell fractions (CCF) were calculated for each tumor subpopulation as the cellular prevalence for a given subpopulation divided by the maximum cellular prevalence for that tumor, which corresponds to the estimated tumor purity. Events were defined as clonal if they have a CCF of 1 or subclonal otherwise. SCNA subpopulation assignments and cellular prevalence estimates derived from phyloWGS were further informed by scRNAseq and scRRBS-derived copy number profiles.

#### Bulk RNA sequencing processing

FASTQ files were pre-processed with fastp v0.20.0 to assess quality control and were aligned to the hg19 genome using kallisto v0.46.0 with default parameters (Bray et al., 2016). The bulk RNA Verhaak classification and simplicity scores were determined as previously reported (Wang et al., 2017). Single sample gene set enrichment analysis for particular pathways was performed using the GVSA R package (Hanzelmann et al., 2013).

#### Detection of extrachromosomal DNA

Amplicon architect was used to detect extrachromosomal DNA in tumor whole genome sequencing data as previously described (Deshpande et al., 2019). Briefly, this method characterizes the architecture of amplified regions that are larger than 10kb and have more than four copies greater than the median sample ploidy.

#### DNA methylation-based tumor classification

Probabilistic estimates of tumor classification were defined both by the Molecular Neuropathology classification tool (version 11b4) as previously reported (Capper et al., 2018).

#### Statistical methods

All data analyses were conducted in R 3.6.1. Statistical analyses are described in the respective Methods subsection and briefly described in the figure legends. No statistical methods were used to predetermine study sample size. *p*-values of < 0.05 were considered statistically significant.

