## Supplementary Figures for "Single-cell multimodal glioma analyses reveal epigenetic regulators of cellular plasticity and environmental stress response"

### Figure S1.

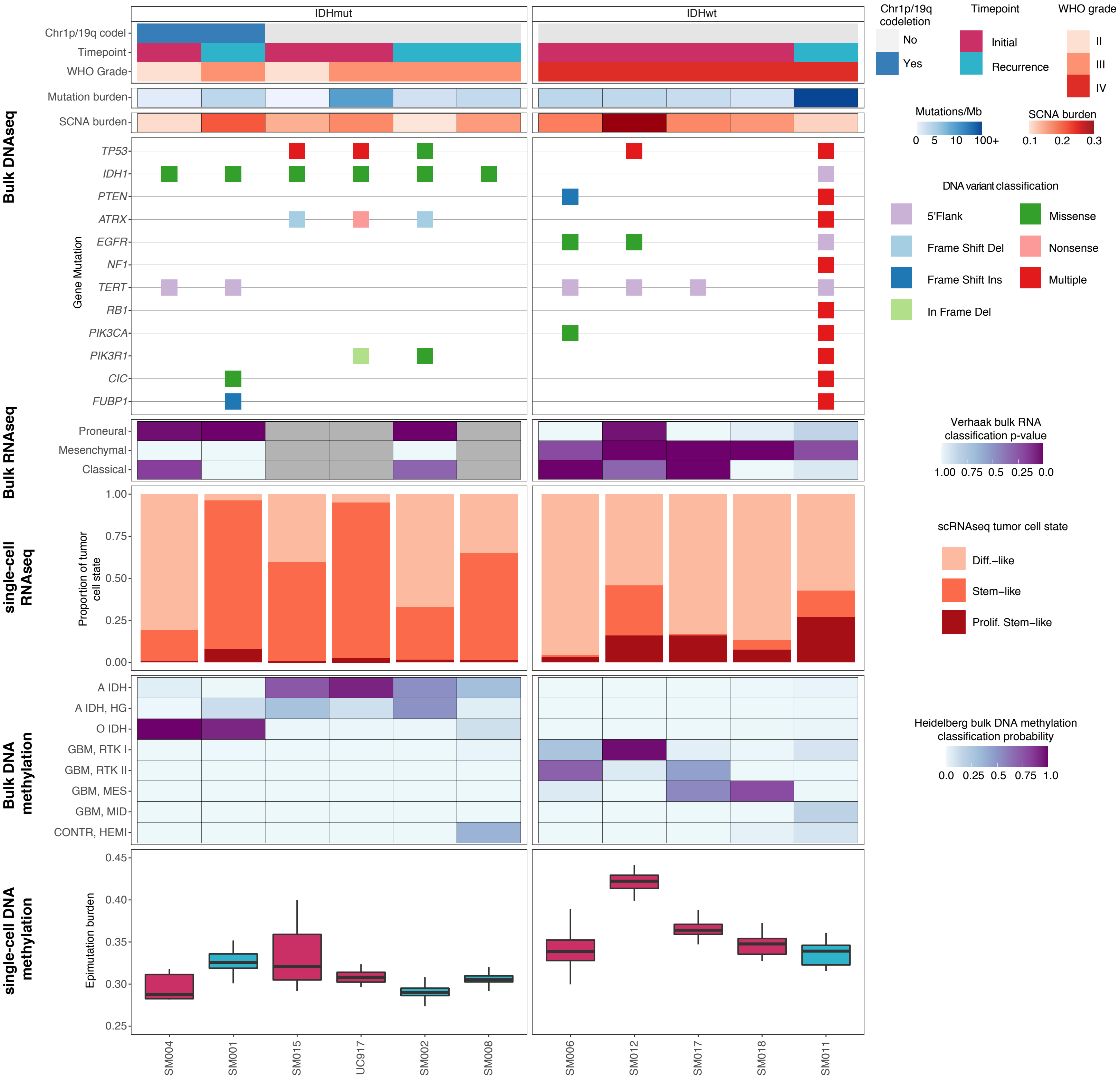
IDHmut
IDHwt
Bulk DNAseq
Chr1p/19q codel
Timepoint
WHO Grade
Mutation burden
SCNA burden
Gene Mutation
TP53
IDH1
PTEN
ATRX
EGFR
NF1
TERT
RB1
PIK3CA
PIK3R1
CIC
FUBP1
Bulk RNAseq
Proneural
Mesenchymal
Classical
single-cell RNAseq
Proportion of tumor cell state
1.00
0.75
0.50
0.25
0.00
Bulk DNA methylation
A IDH
A IDH, HG
O IDH
GBM, RTK I
GBM, RTK II
GBM, MES
GBM, MID
CONTR, HEMI
single-cell DNA methylation
Epimutation burden
0.45
0.40
0.35
0.30
0.25
SM004
SM001
SM015
UC917
SM002
SM008
SM006
SM012
SM017
SM018
SM011
Chr1p/19q codeletion
No
Yes
Timepoint
Initial
Recurrence
WHO grade
II
III
IV
Mutations/Mb
0
5
10
100+
SCNA burden
0.1
0.2
0.3
DNA variant classification
5'Flank
Frame Shift Del
Frame Shift Ins
In Frame Del
Missense
Nonsense
Multiple
Verhaak bulk RNA classification p-value
1.00
0.75
0.5
0.25
0.0
scRNAseq tumor cell state
Diff.–like
Stem–like
Prolif. Stem–like
Heidelberg bulk DNA methylation classification probability
0.0
0.25
0.5
0.75
1.0

**Figure S1. Integrated molecular profiles of patient samples. Related to Figure 1.**
Each patient is in a single column with data presented to indicate clinical features (top), followed by genetic alterations defined from bulk whole genome sequencing data, bulk RNA sequencing based subtype classification probabilities (Wang et al., $n$ = 8 available), single-cell RNA tumor cellular state proportions, bulk DNA methylation microarray subtype classification probabilities (Capper et al.), and boxplots of single-cell epimutation burden with samples colored by clinical timepoint.

**Figure S2.**

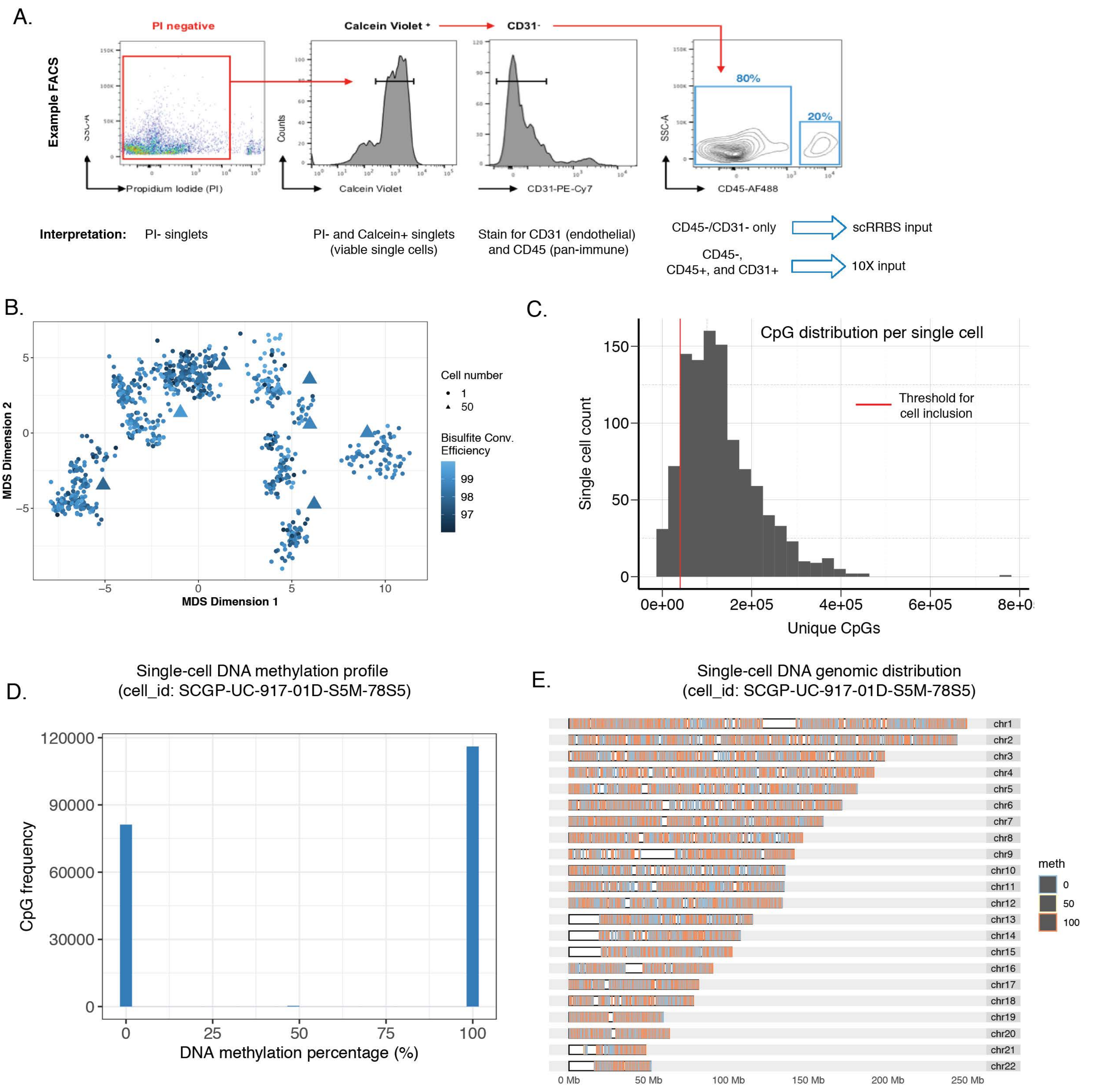


**Figure S2. Sample pre-processing and metrics related to single-cell DNA methylation data assessment. Related to Figure 1.**

(A) Representative fluorescence activated cell sorting (FACS) data and strategy for viable cell enrichment for both single-cell protocols, and tumor cell enrichment in scRRBS. (B) The same multidimensional scaling (MDS) analysis using pairwise distance metrics calculated between individual cells as in Figure 1B, except colored by bisulfite conversion efficiency. (C) The number of unique CpGs detected per single cell, with the red line indicating the threshold (minimum 40,000 unique CpGs) for inclusion in the dataset presented herein. (D) Representative distribution of single locus DNA methylation estimates for a single cell. DNA methylation percentage of 0 represents an unmethylated locus, while a percentage of 100 represents a methylated locus. (E) Representative genomic distribution of DNA methylation values within a single cell.

### Figure S3.

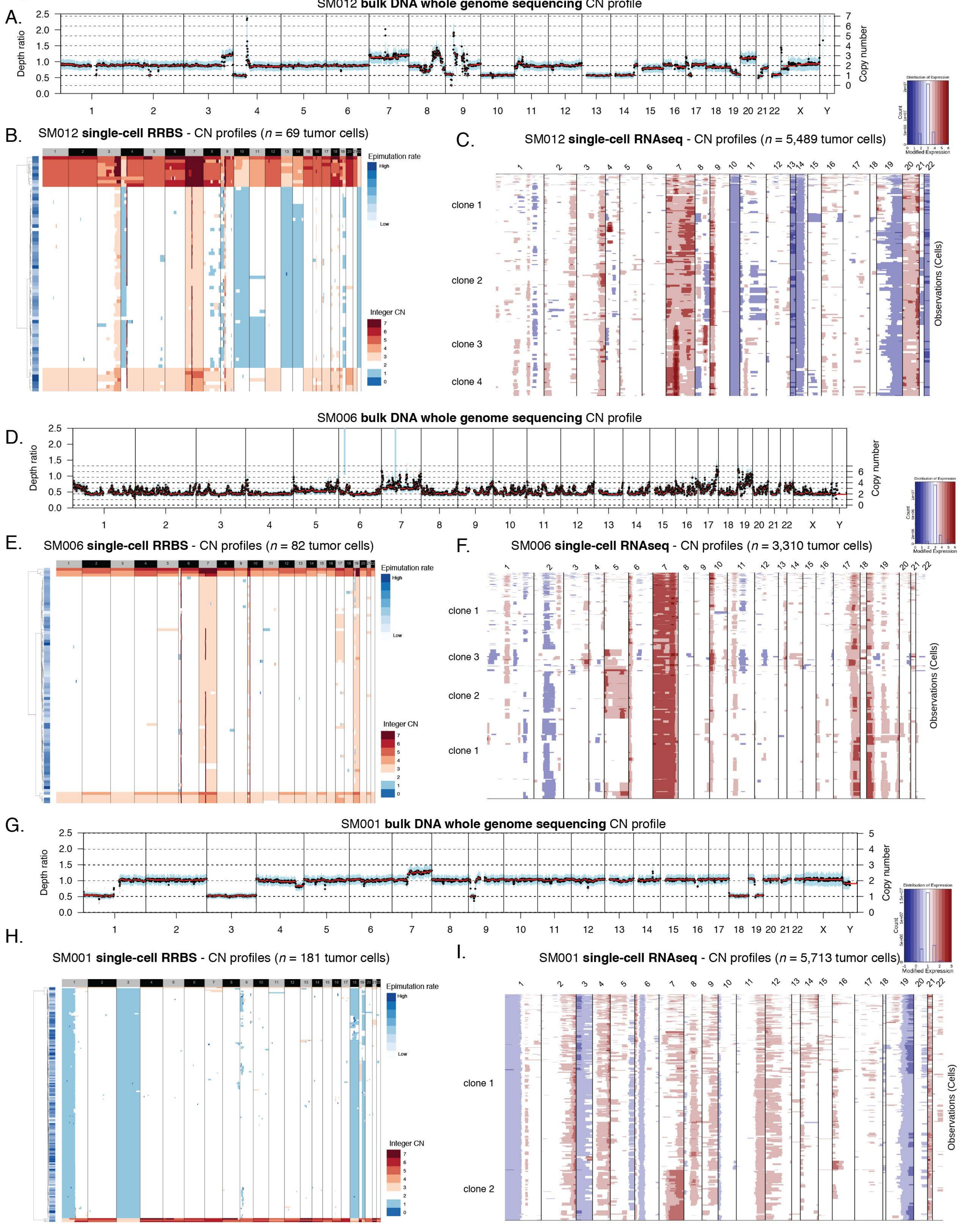
A.
SM012 bulk DNA whole genome sequencing CN profile
Depth ratio
Copy number
B.
SM012 single-cell RRBS - CN profiles (n = 69 tumor cells)
Epimutation rate
High
Low
Integer CN
C.
SM012 single-cell RNAseq - CN profiles (n = 5,489 tumor cells)
clone 1
clone 2
clone 3
clone 4
Observations (Cells)
Distribution of Expression
Count
Modified Expression
D.
SM006 bulk DNA whole genome sequencing CN profile
Depth ratio
Copy number
E.
SM006 single-cell RRBS - CN profiles (n = 82 tumor cells)
Epimutation rate
High
Low
Integer CN
F.
SM006 single-cell RNAseq - CN profiles (n = 3,310 tumor cells)
clone 1
clone 3
clone 2
clone 1
Observations (Cells)
G.
SM001 bulk DNA whole genome sequencing CN profile
Depth ratio
Copy number
H.
SM001 single-cell RRBS - CN profiles (n = 181 tumor cells)
Epimutation rate
High
Low
Integer CN
I.
SM001 single-cell RNAseq - CN profiles (n = 5,713 tumor cells)
clone 1
clone 2
Observations (Cells)

**Figure S3. Somatic copy number alteration examples estimated from whole genome sequencing, single-cell Reduced Representation Bisulfite Sequencing, and single-cell RNA-sequencing. Related to Figure 1.**

(A-C) Representative images of copy number alterations derived from SM012 (IDHwt initial) whole genome sequencing (WGS) data. (A) Depth ratio for each segment with copy number status determined as compared with germline (normal blood) WGS data. (B) SM012 Single-cell DNA methylation-based copy number estimates (*n* = 69 tumor cells) with copy number integer depicted by color (blue = CN loss, white = neutral CN, and red = CN gain). Each row is a single cell with annotation for epimutation burden provided. (C) SM012 Single-cell RNAseq based copy number inference (*n* = 5,489) identifying major copy number events found in WGS with labelled subclones as presented in Figure 4D. (D-F) Similar example profiles as presented in (A-C) for tumor sample SM006 (IDHwt initial, *n* = 82 scRRBS cells, *n* = 3,310 scRNAseq cells). (G-I) Similar example profiles as presented in (A-C) for tumor sample SM001 (IDHmut recurrence, *n* = 181 scRRBS cells, *n* = 5,713 scRNAseq cells).

**Figure S4.**

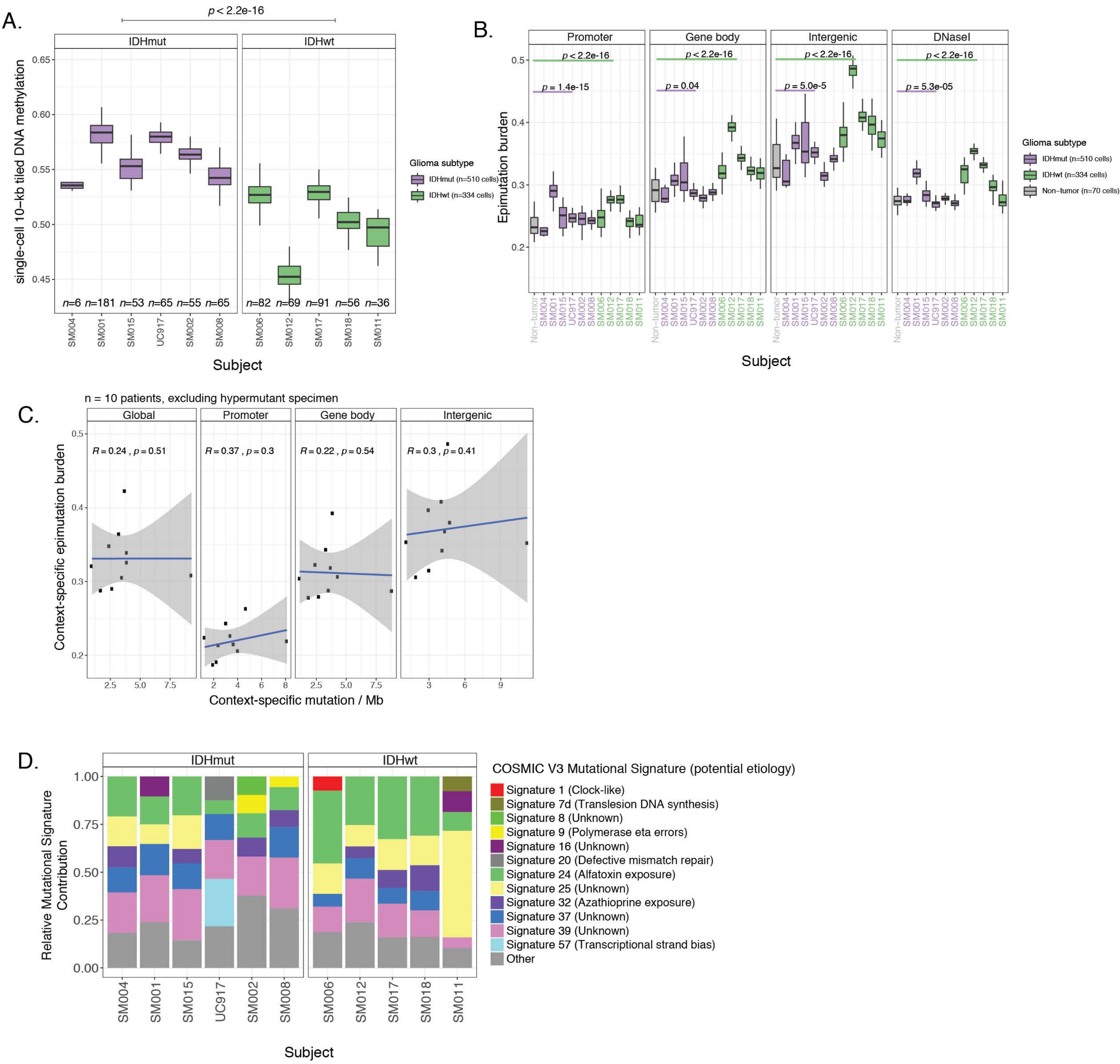


**Figure S4. Distribution and relationship of DNA methylation and epimutation throughout the glioma genome. Related to Figure 1.**
(A) Boxplots representing average 10-kb tiled DNA methylation values per single tumor cell. (B) Boxplots highlighting the single-cell epimutation burden estimates calculated across different genomic contexts. (C) Scatterplots showing the relationship between genomic context-specific single-cell epimutation burden (sample-specific scRRBS average) and genomic context-specific mutation burden derived from whole genome sequencing (n = 10 excluding hypermutant sample). Panels are separated into global (i.e., all regions), promoter, gene body, and intergenic regions (Spearman correlations p > 0.05 for all comparisons). (D) The dominant Catalogue of Somatic Mutations in Cancer (COSMIC v3) mutational signatures are presented for each subject. The stacked bar plots represent the relative contribution of each mutational signature to the tumor's mutational burden. Colors indicate distinct mutational signatures, which are further annotated with their proposed etiology.

**Figure S5.**

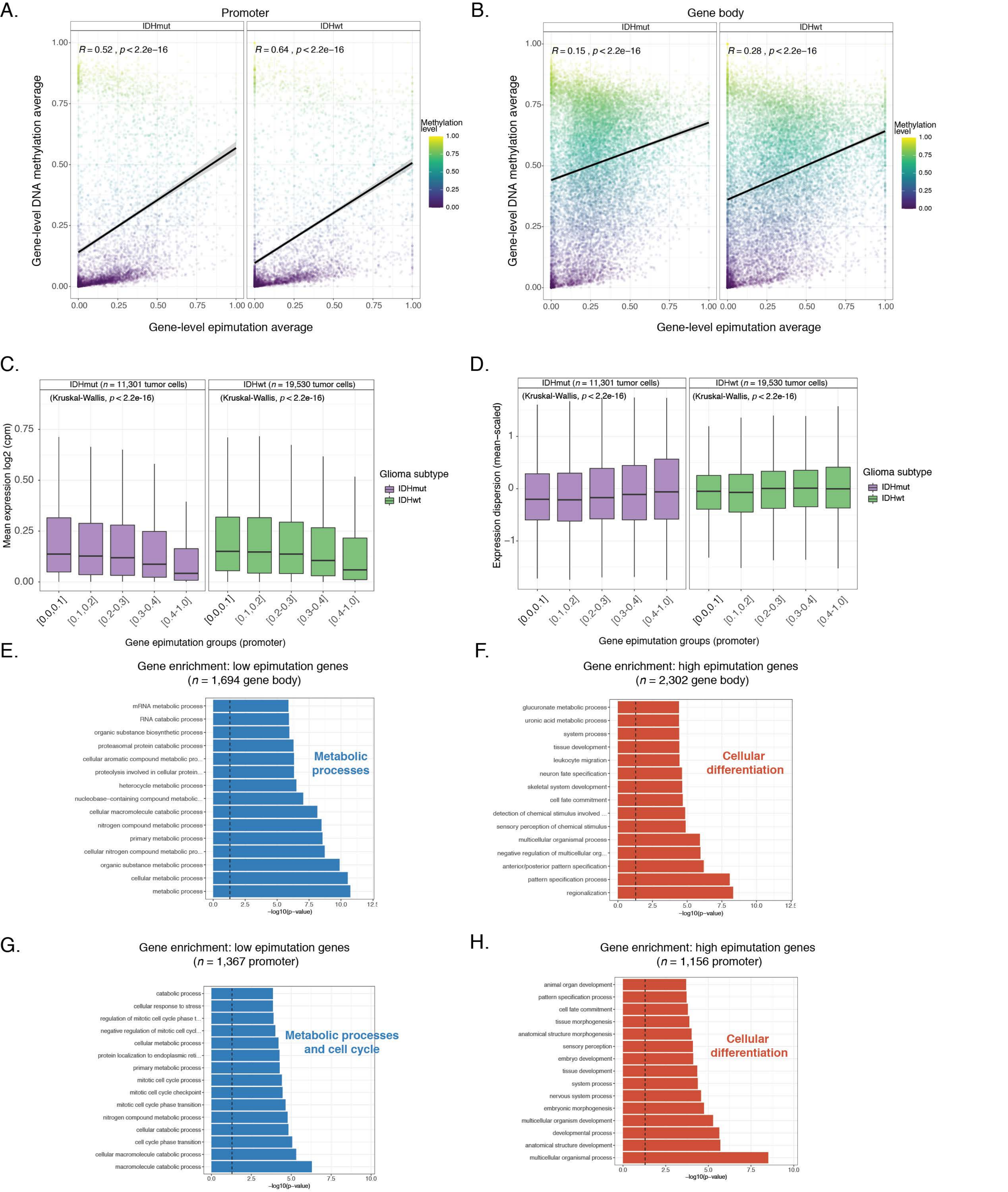
A.
Promoter
IDHmut
IDHwt
R = 0.52 , p < 2.2e−16
R = 0.64 , p < 2.2e−16
Gene-level DNA methylation average
Gene-level epimutation average
Methylation level
1.00
0.75
0.50
0.25
0.00
B.
Gene body
IDHmut
IDHwt
R = 0.15 , p < 2.2e−16
R = 0.28 , p < 2.2e−16
Gene-level DNA methylation average
Gene-level epimutation average
Methylation level
C.
IDHmut (n = 11,301 tumor cells)
IDHwt (n = 19,530 tumor cells)
(Kruskal-Wallis, p < 2.2e-16)
Mean expression log2 (cpm)
Glioma subtype
IDHmut
IDHwt
[0.0,0.1]
[0.1,0.2]
[0.2-0.3]
[0.3-0.4]
[0.4-1.0]
Gene epimutation groups (promoter)
D.
IDHmut (n = 11,301 tumor cells)
IDHwt (n = 19,530 tumor cells)
(Kruskal-Wallis, p < 2.2e-16)
Expression dispersion (mean−scaled)
Glioma subtype
IDHmut
IDHwt
Gene epimutation groups (promoter)
E.
Gene enrichment: low epimutation genes
(n = 1,694 gene body)
mRNA metabolic process
RNA catabolic process
organic substance biosynthetic process
proteasomal protein catabolic process
cellular aromatic compound metabolic pro...
proteolysis involved in cellular protein...
heterocycle metabolic process
nucleobase−containing compound metabolic...
cellular macromolecule catabolic process
nitrogen compound metabolic process
primary metabolic process
cellular nitrogen compound metabolic pro...
organic substance metabolic process
cellular metabolic process
metabolic process
Metabolic processes
−log10(p−value)
F.
Gene enrichment: high epimutation genes
(n = 2,302 gene body)
glucuronate metabolic process
uronic acid metabolic process
system process
tissue development
leukocyte migration
neuron fate specification
skeletal system development
cell fate commitment
detection of chemical stimulus involved ...
sensory perception of chemical stimulus
multicellular organismal process
negative regulation of multicellular org...
anterior/posterior pattern specification
pattern specification process
regionalization
Cellular differentiation
−log10(p−value)
G.
Gene enrichment: low epimutation genes
(n = 1,367 promoter)
catabolic process
cellular response to stress
regulation of mitotic cell cycle phase t...
negative regulation of mitotic cell cycl...
cellular metabolic process
protein localization to endoplasmic reti...
primary metabolic process
mitotic cell cycle process
mitotic cell cycle checkpoint
mitotic cell cycle phase transition
nitrogen compound metabolic process
cellular catabolic process
cell cycle phase transition
cellular macromolecule catabolic process
macromolecule catabolic process
Metabolic processes and cell cycle
−log10(p−value)
H.
Gene enrichment: high epimutation genes
(n = 1,156 promoter)
animal organ development
pattern specification process
cell fate commitment
tissue morphogenesis
anatomical structure morphogenesis
sensory perception
embryo development
tissue development
system process
nervous system process
embryonic morphogenesis
multicellular organism development
developmental process
anatomical structure development
multicellular organismal process
Cellular differentiation
−log10(p−value)

**Figure S5. Association between epimutation and disrupted transcriptional programs. Related to Figure 1.**
(A-B) Scatterplots depicting single-cell gene-level epimutation average plotted against the gene-level methylation estimates in both (A) promoter regions and (B) gene body regions. (C) Boxplots of gene expression values, in log2 (counts per million), from single-cell RNAseq data across different sets of promoter regions defined by gene-derived epimutation groups. Gene epimutation groups are defined by the determining the mean epimutation value across a single gene. Color indicates *IDH1* mutation status. (D) Boxplots of gene expression dispersion that were mean-expression scaled to account for expression level-dependent variability across the same promoter-based gene epimutation groups defined in panel C. (E-F) Gene Ontology enrichment analyses for low epimutation genes (Figure 1E, mean epimutation across all tumor cells: 0.0 - 0.1) and high epimutation genes (Figure 1F, mean epimutation across all tumor cells: > 0.5) using gene body estimates. A meta biological process is placed next to significant Gene Ontology terms. (G-H) Same analyses presented in panels E-F, but for gene-level epimutation estimates determined in promoters.

**Figure S6.**

A.

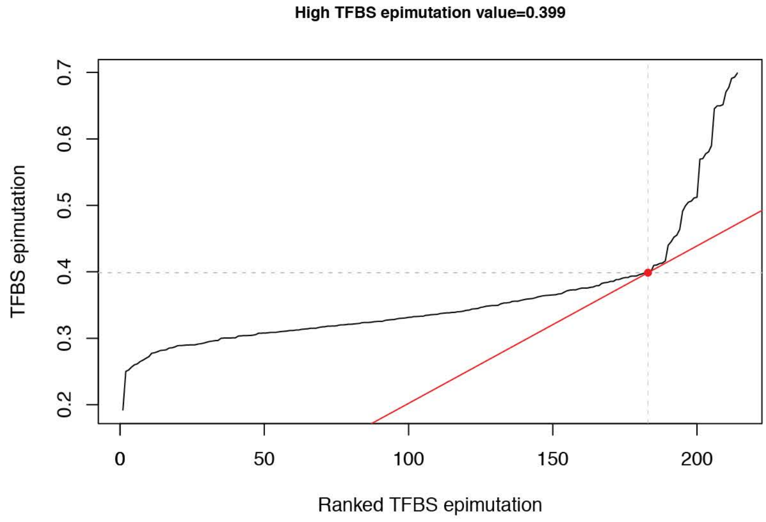


B.

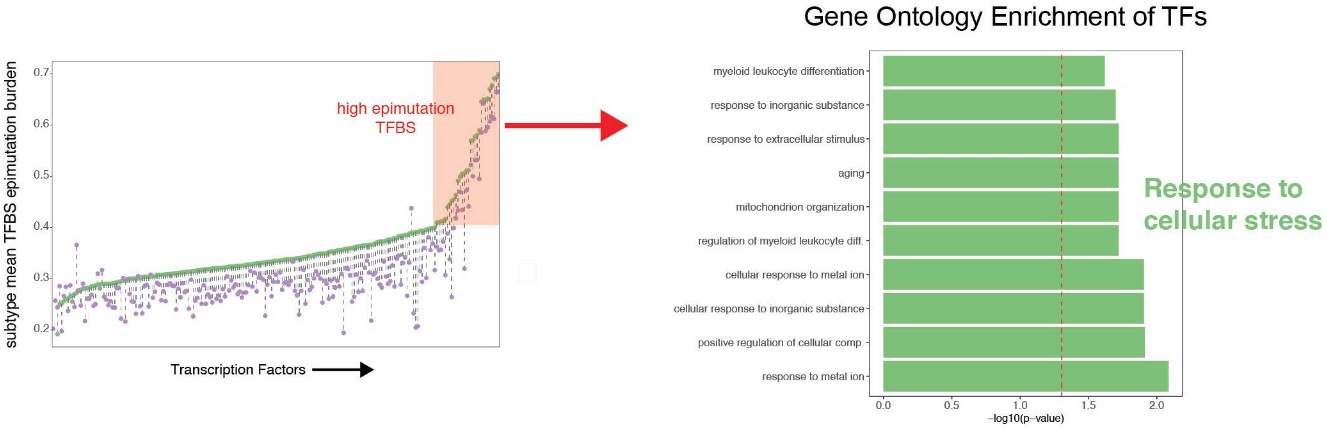


**Figure S6. Enrichment of high epimutation transcription factors and association with environmental stress response. Related to Figure 1.**

(A) Computational approach to defining TFBSs with high epimutation burden (red tangent line at 0.399 TFBS epimutation burden). X-axis represents each TF ordered by mean epimutation burden in IDHwt single-cells (n = 334 cells).

(B) Gene Ontology enrichment analysis of TFs with high epimutation burden in their binding sites. A meta biological process is placed next to significant Gene Ontology (GO) terms.

**Figure S7.**

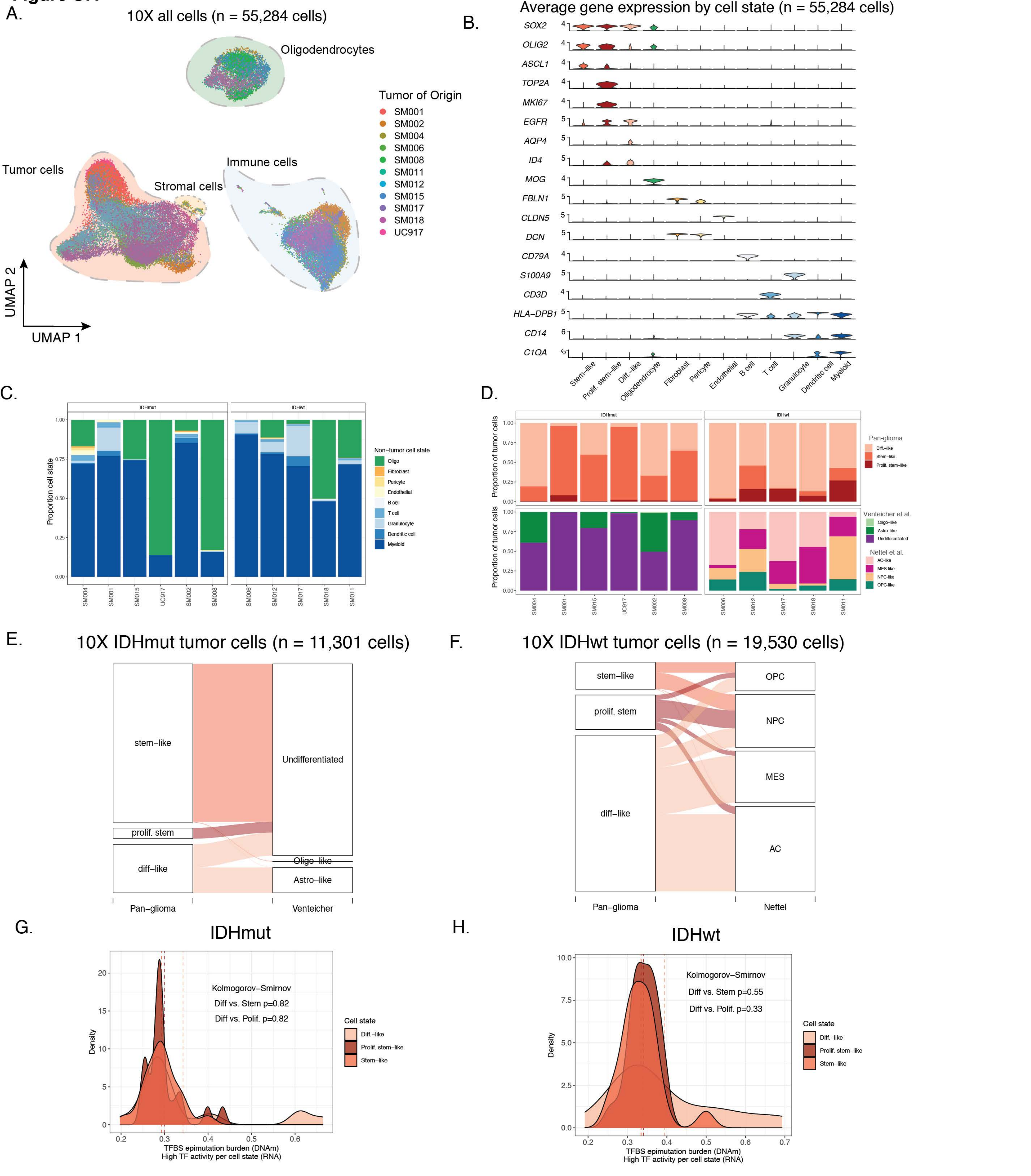
A.
10X all cells (n = 55,284 cells)
Oligodendrocytes
Tumor of Origin
SM001
SM002
SM004
SM006
SM008
SM011
SM012
SM015
SM017
SM018
UC917
Tumor cells
Stromal cells
Immune cells
UMAP 2
UMAP 1
B.
Average gene expression by cell state (n = 55,284 cells)
SOX2
OLIG2
ASCL1
TOP2A
MKI67
EGFR
AQP4
ID4
MOG
FBLN1
CLDN5
DCN
CD79A
S100A9
CD3D
HLA-DPB1
CD14
C1QA
Stem-like
Prolif. stem-like
Diff.-like
Oligodendrocyte
Fibroblast
Pericyte
Endothelial
B cell
T cell
Granulocyte
Dendritic cell
Myeloid
C.
IDHmut
IDHwt
Proportion cell state
Non-tumor cell state
Oligo
Fibroblast
Pericyte
Endothelial
B cell
T cell
Granulocyte
Dendritic cell
Myeloid
SM004
SM001
SM015
UC917
SM002
SM008
SM006
SM012
SM017
SM018
SM011
D.
Proportion of tumor cells
Pan-glioma
Diff.-like
Stem-like
Prolif. stem-like
Venteicher et al.
Oligo-like
Astro-like
Undifferentiated
Neftel et al.
AC-like
MES-like
NPC-like
OPC-like
E.
10X IDHmut tumor cells (n = 11,301 cells)
stem-like
prolif. stem
diff-like
Undifferentiated
Oligo-like
Astro-like
Pan-glioma
Venteicher
F.
10X IDHwt tumor cells (n = 19,530 cells)
stem-like
prolif. stem
diff-like
OPC
NPC
MES
AC
Pan-glioma
Neftel
G.
IDHmut
Kolmogorov-Smirnov
Diff vs. Stem p=0.82
Diff vs. Polif. p=0.82
Density
Cell state
Diff.-like
Prolif. stem-like
Stem-like
TFBS epimutation burden (DNAm)
High TF activity per cell state (RNA)
H.
IDHwt
Kolmogorov-Smirnov
Diff vs. Stem p=0.55
Diff vs. Polif. p=0.33
Density
Cell state
Diff.-like
Prolif. stem-like
Stem-like
TFBS epimutation burden (DNAm)
High TF activity per cell state (RNA)

**Figure S7. Pan-glioma cell state assignment and characteristics. Related to Figure 2.** (A) UMAP dimensionality reduction plot of all scRNAseq data, including tumor and non-tumor cells (*n* = 55,248 cells). Each dot depicts a single cell and colors represents the tumor of origin. Shaded regions represent cell state classification. (B) Stacked violin plots of average single-cell gene expression for cells presented in Figure S7A. Selected genes presented are informative for cell state classification. (C) Stacked bar plots representing the proportion of non-tumor cellular states (D) Stacked bar plots representing the proportion of tumor cellular states per tumor for pan-glioma classification (top row) and previously published classifications (middle row; Venteicher et al. and Neftel et al.) (E) Sankey plot representing the proportion of IDHmut tumor cells with pan-glioma classification and associated classification described in Venteicher et al. (F) Sankey plot representing the proportion of IDHwt tumor cells with pan-glioma classification and associated classification described in Neftel et al. (G) Density plots representing TFBS epimutation burden (scRRBS data) in IDHmut single-cell DNA methylation data for TFs whose activity (scRNAseq based SCENIC analysis) characterizes a specific cell state (*n* = 20 TFs per cell state). Kolmogorov-Smirnov *p*-value tests for differences in TFBS epimutation burden across the cellular states. (H) Density plots representing TFBS epimutation burden (scRRBS data) in IDHwt single-cell DNA methylation data for TFs whose activity (scRNAseq based SCENIC analysis) characterizes a specific cell state (*n* = 20 TFs per cell state). Dotted lines represent the median TFBS value for cell state defining TFs. The Kolmogorov-Smirnov *p*-value corresponds to differences in TFBS epimutation burden across the cellular states.

#### Figure S8.

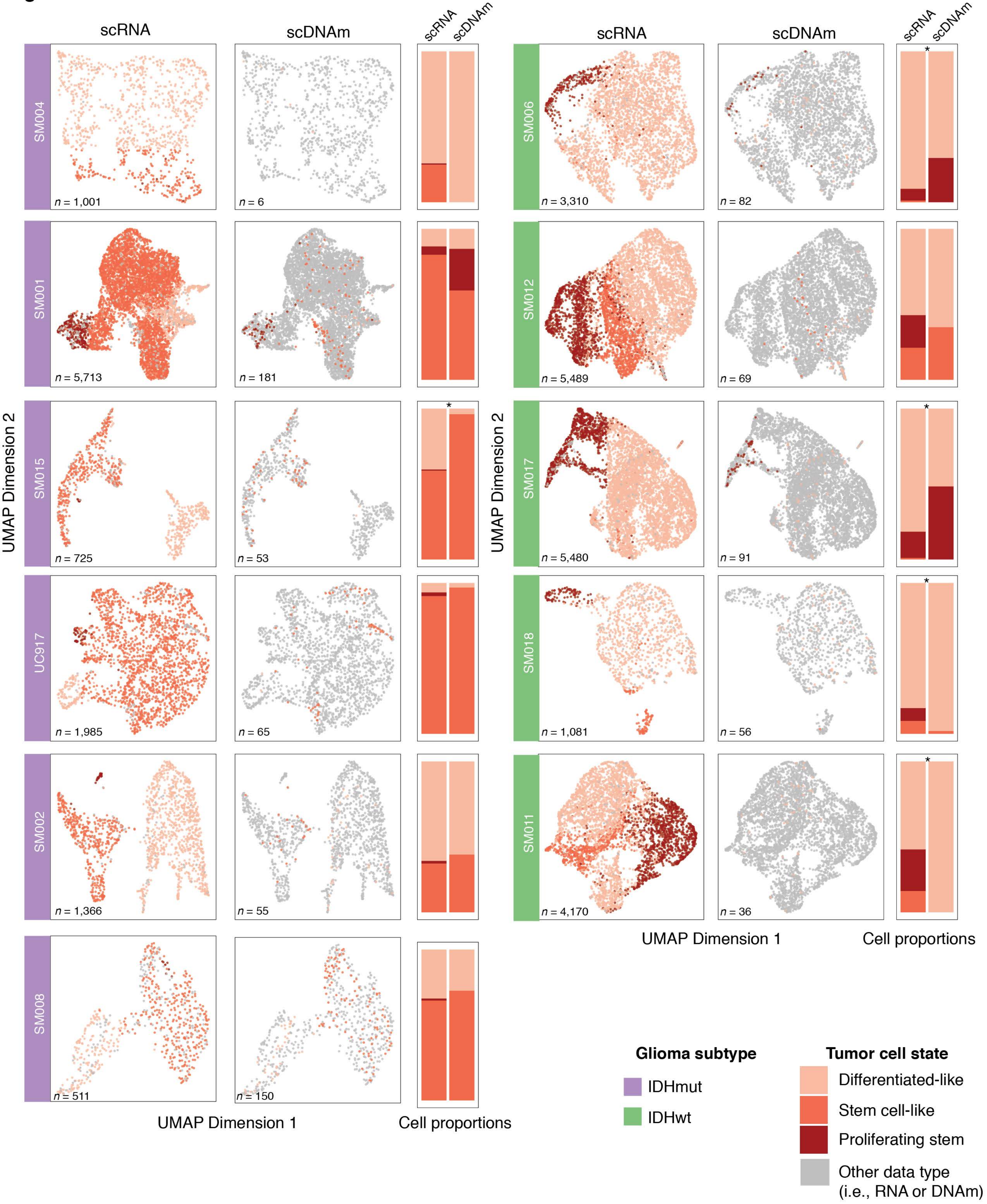

scRNA
scDNAm
scRNA
scDNAm
SM004
n = 1,001
n = 6
SM001
n = 5,713
n = 181
SM015
n = 725
n = 53
UC917
n = 1,985
n = 65
SM002
n = 1,366
n = 55
SM008
n = 511
n = 150
SM006
n = 3,310
n = 82
SM012
n = 5,489
n = 69
SM017
n = 5,480
n = 91
SM018
n = 1,081
n = 56
SM011
n = 4,170
n = 36
*
UMAP Dimension 2
UMAP Dimension 1
Cell proportions
Glioma subtype
IDHmut
IDHwt
Tumor cell state
Differentiated-like
Stem cell-like
Proliferating stem
Other data type
(i.e., RNA or DNAm)

**Figure S8. LIGER integrated tumor-specific clustering of single-cell RNA and single-cell DNA methylation data. Related to Figure 2.**
Joint single-cell RNAseq (scRNA) and single-cell DNA methylation (scDNAm) clustering and UMAP projections highlighting similar cellular state distributions across platforms. Sample annotation is presented on the left of each paired UMAP plot, each dot is an individual single cell, and cell number for each technology is presented in the lower-left hand corner. UMAP coordinate space remains the same for both scRNA and scDNAm visualizations with cellular states for that platform represented by a colored dot and data for the other platform represented by a gray dot. Stacked bar plots enumerating the proportion of cellular states detected by each platform are presented to the right of each paired UMAP plot. \`*\` indicate specimens in which the cellular proportions across the two platforms are significantly different (Fisher's Exact test, $p < 0.05$).

**Figure S9.**

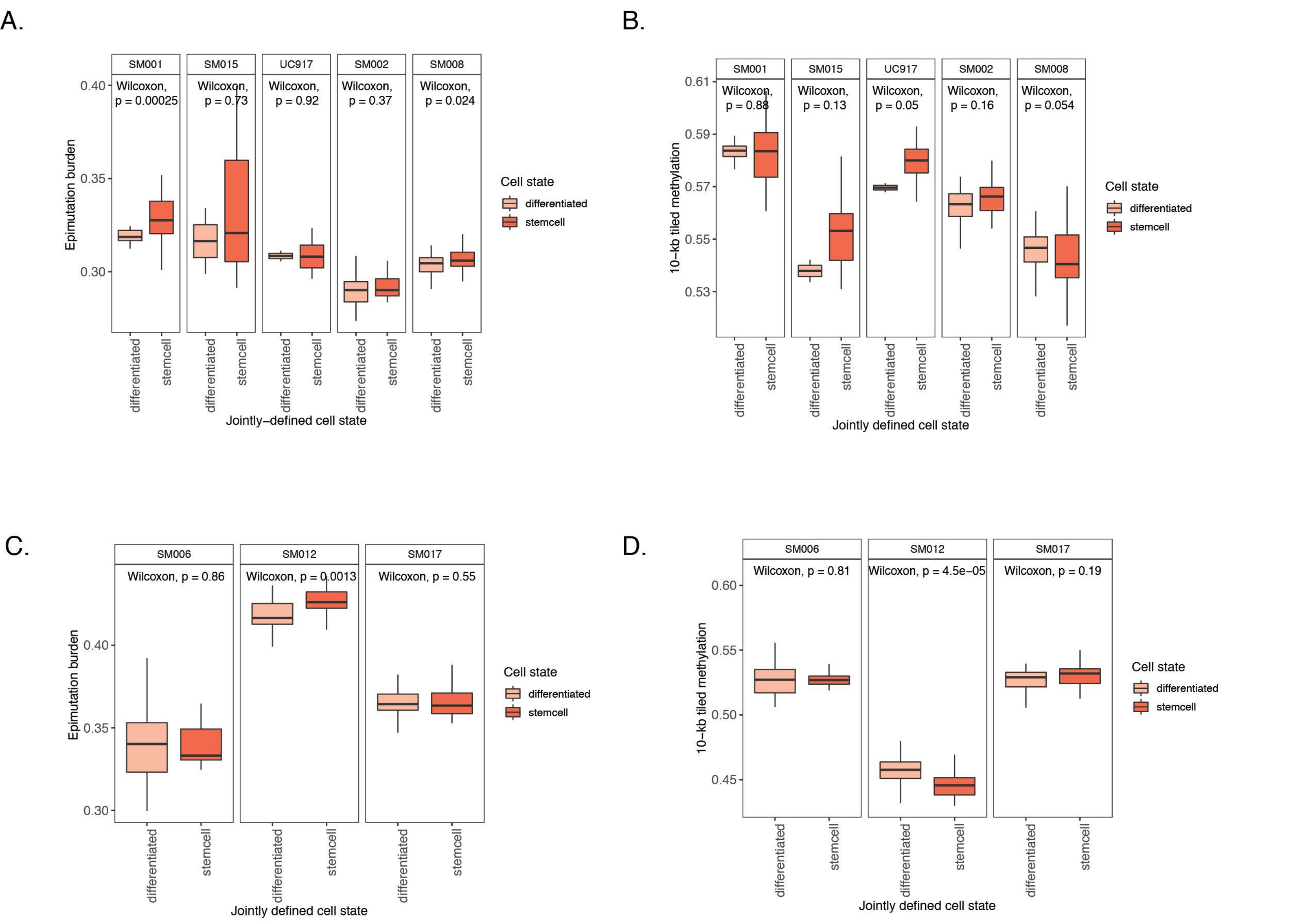


**Figure S9. Sample-specific differences in DNA methylation and epimutation burden across different cellular states. Related to Figure 2.**
(A-B) Boxplots showing sample-specific differences in (A) epimutation burden and (B) 10-kb tiled DNA methylation across LIGER-defined cellular states in IDHmut tumors. Wilcoxon Rank Sum p-values are presented comparing cells from a given tumor. (C-D) Boxplots showing sample-specific differences in (C) epimutation burden and (D) 10-kb tiled DNA methylation across LIGER-defined cellular states in IDHwt tumors. Wilcoxon Rank Sum p-values are presented comparing cells from a given tumor. Samples with only one defined cell state are not visualized.

**Figure S10.**

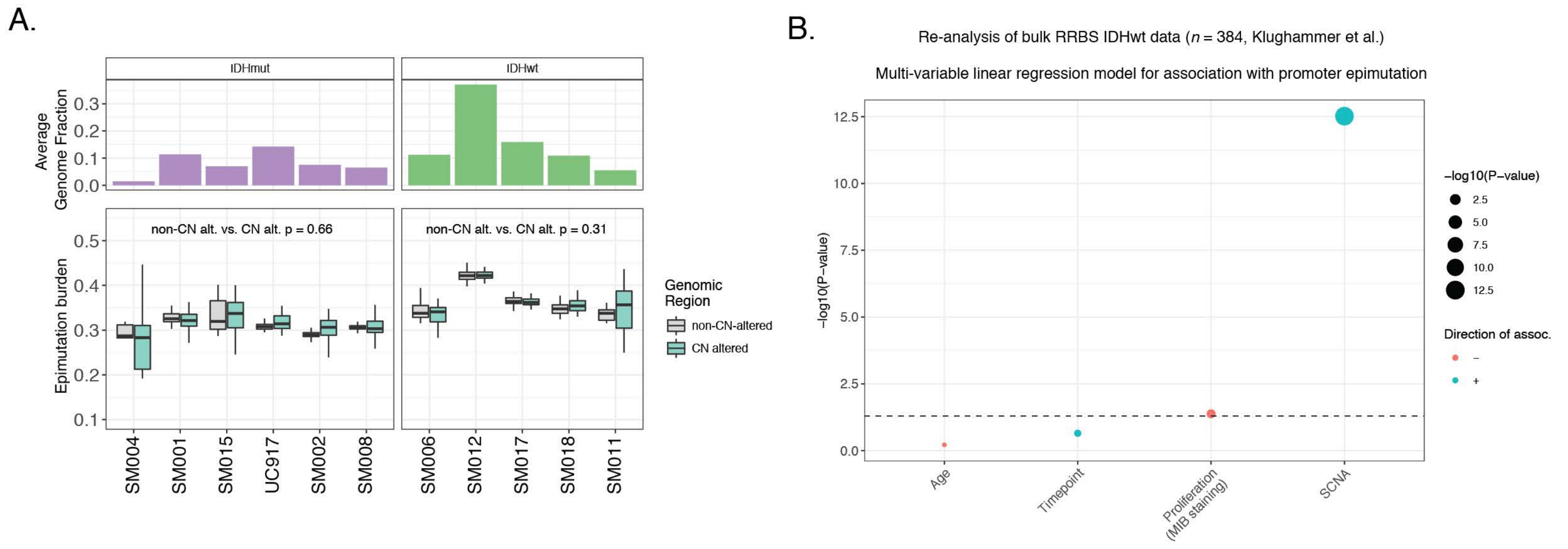


**Figure S10. Relationships between epimutation burden and genetic alterations. Related to Figure 3.**
(A) Single-cell epimutation burden estimates were calculated across genomic regions with (teal) and without (gray) copy number alterations. The paired-sample Wilcoxon test p-value for each subtype represents the statistical difference of epimutation burden across these two regions. (B) Visualized results from multi-variable linear regression model testingfor association with epimutation burden. Dot size indicates -log10 (p-value) for each predictor and color represents direction of association with epimutation burden (red = negative association, blue = positive association). Explanatory variables included subject age, timepoint (pre- and post-treatment), level of cellular proliferation determined by histological marker (MIB staining), and somatic copy number alteration burden (SCNA, total number of bases altered / total number of bases measured).

**Figure S11.**

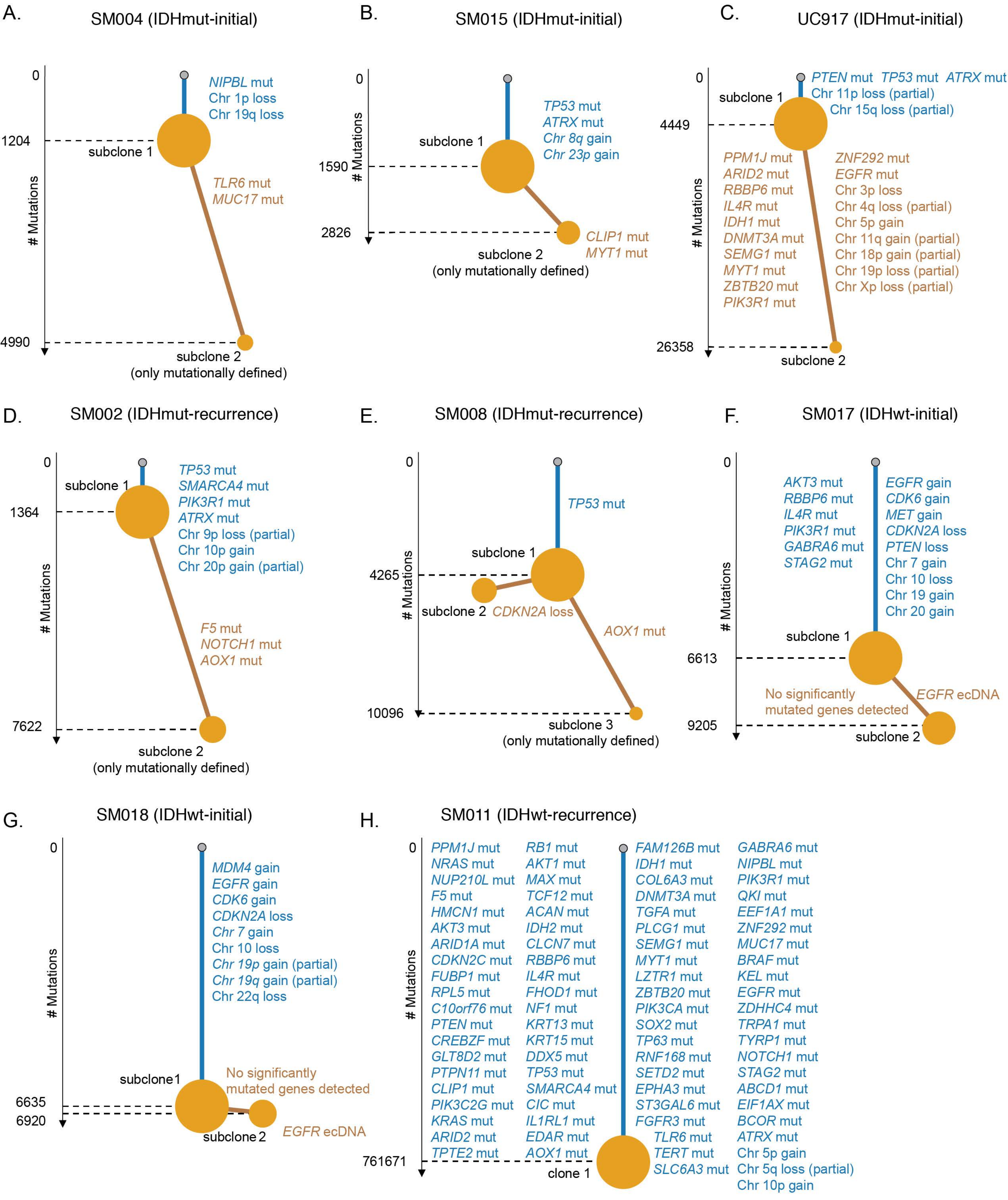
A.
SM004 (IDHmut-initial)
NIPBL mut
Chr 1p loss
Chr 19q loss
subclone 1
TLR6 mut
MUC17 mut
subclone 2
(only mutationally defined)
### Mutations
0
1204
4990
B.
SM015 (IDHmut-initial)
TP53 mut
ATRX mut
Chr 8q gain
Chr 23p gain
subclone 1
CLIP1 mut
MYT1 mut
subclone 2
(only mutationally defined)
### Mutations
0
1590
2826
C.
UC917 (IDHmut-initial)
PTEN mut TP53 mut ATRX mut
Chr 11p loss (partial)
Chr 15q loss (partial)
subclone 1
PPM1J mut
ARID2 mut
RBBP6 mut
IL4R mut
IDH1 mut
DNMT3A mut
SEMG1 mut
MYT1 mut
ZBTB20 mut
PIK3R1 mut
ZNF292 mut
EGFR mut
Chr 3p loss
Chr 4q loss (partial)
Chr 5p gain
Chr 11q gain (partial)
Chr 18p gain (partial)
Chr 19p loss (partial)
Chr Xp loss (partial)
subclone 2
### Mutations
0
4449
26358
D.
SM002 (IDHmut-recurrence)
TP53 mut
SMARCA4 mut
PIK3R1 mut
ATRX mut
Chr 9p loss (partial)
Chr 10p gain
Chr 20p gain (partial)
subclone 1
F5 mut
NOTCH1 mut
AOX1 mut
subclone 2
(only mutationally defined)
### Mutations
0
1364
7622
E.
SM008 (IDHmut-recurrence)
TP53 mut
subclone 1
subclone 2
CDKN2A loss
AOX1 mut
subclone 3
(only mutationally defined)
### Mutations
0
4265
10096
F.
SM017 (IDHwt-initial)
AKT3 mut
RBBP6 mut
IL4R mut
PIK3R1 mut
GABRA6 mut
STAG2 mut
EGFR gain
CDK6 gain
MET gain
CDKN2A loss
PTEN loss
Chr 7 gain
Chr 10 loss
Chr 19 gain
Chr 20 gain
subclone 1
No significantly
mutated genes detected
EGFR ecDNA
subclone 2
### Mutations
0
6613
9205
G.
SM018 (IDHwt-initial)
MDM4 gain
EGFR gain
CDK6 gain
CDKN2A loss
Chr 7 gain
Chr 10 loss
Chr 19p gain (partial)
Chr 19q gain (partial)
Chr 22q loss
subclone1
No significantly
mutated genes detected
subclone 2
EGFR ecDNA
### Mutations
0
6635
6920
H.
SM011 (IDHwt-recurrence)
PPM1J mut
NRAS mut
NUP210L mut
F5 mut
HMCN1 mut
AKT3 mut
ARID1A mut
CDKN2C mut
FUBP1 mut
RPL5 mut
C10orf76 mut
PTEN mut
CREBZF mut
GLT8D2 mut
PTPN11 mut
CLIP1 mut
PIK3C2G mut
KRAS mut
ARID2 mut
TPTE2 mut
RB1 mut
AKT1 mut
MAX mut
TCF12 mut
ACAN mut
IDH2 mut
CLCN7 mut
RBBP6 mut
IL4R mut
FHOD1 mut
NF1 mut
KRT13 mut
KRT15 mut
DDX5 mut
TP53 mut
SMARCA4 mut
CIC mut
IL1RL1 mut
EDAR mut
AOX1 mut
FAM126B mut
IDH1 mut
COL6A3 mut
DNMT3A mut
TGFA mut
PLCG1 mut
SEMG1 mut
MYT1 mut
LZTR1 mut
ZBTB20 mut
PIK3CA mut
SOX2 mut
TP63 mut
RNF168 mut
SETD2 mut
EPHA3 mut
ST3GAL6 mut
FGFR3 mut
TLR6 mut
TERT mut
SLC6A3 mut
GABRA6 mut
NIPBL mut
PIK3R1 mut
QKI mut
EEF1A1 mut
ZNF292 mut
MUC17 mut
BRAF mut
KEL mut
EGFR mut
ZDHHC4 mut
TRPA1 mut
TYRP1 mut
NOTCH1 mut
STAG2 mut
ABCD1 mut
EIF1AX mut
BCOR mut
ATRX mut
Chr 5p gain
Chr 5q loss (partial)
Chr 10p gain
clone 1
### Mutations
0
761671

**Figure S11. Whole genome sequencing phylogenetic inference of tumor samples. Related to Figure 4.**

(A-H) Phylogenetic trees constructed from whole genome sequencing data (mutations and somatic copy number alterations) using phyloWGS and further annotated using single-cell inferred copy number alterations (scRRBS + scRNAseq). Tree nodes represent alterations specific to the given clone, with node size corresponding to the fraction of cells with the associated alterations. Branch length scales with the number of mutations attributed to that clone. Clonal alterations are colored in blue, with subclonal alterations colored in gold. Genes considered significantly mutated in TCGA analyses (Ceccarelli et al., 2016) and chromosomal arm-level events are presented. Arm-level events are defined as spanning at least 80 percent of the chromosome arm, while partial events span at least 40 percent.

**Figure S12.**

Entanglement of 0.0 = same structure in both datasets.
1.0 = unrelated tree structure.

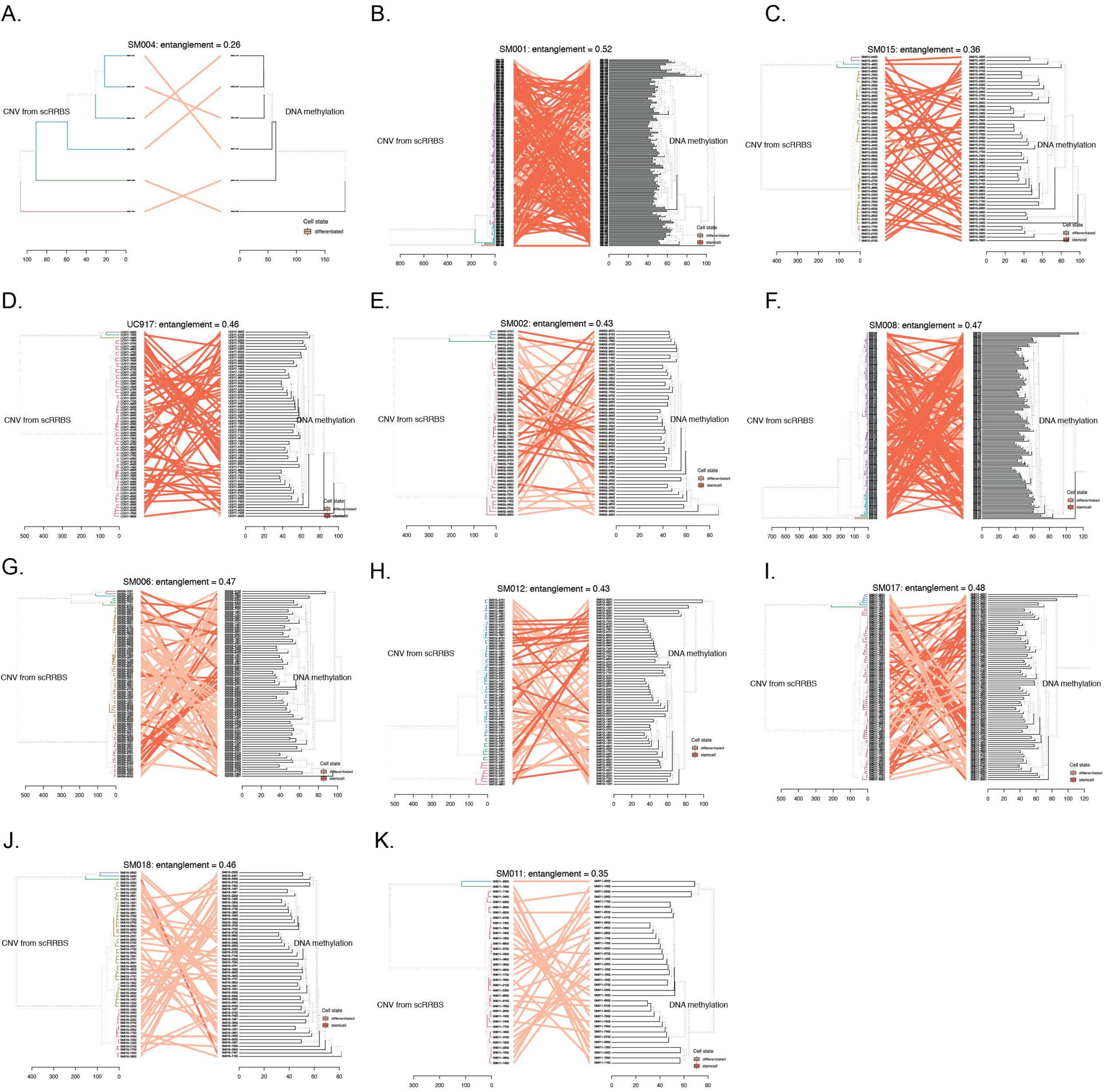


**Figure S12. Tumor-specific comparisons of phylogenetic and phyloepigenetic trees. Related to Figure 4.**
(A-K) Tangelgrams highlighting the relationship between single-cell copy number and single-cell DNA methylation tree diagrams. Phylogenetic trees (left cluster) were calculated from copy number profiles (scRRBS data) and phyloepigenetic trees were constructed from the same cells across 10-kb tiled DNA methylation values. Cluster labels are connected with solid lines and are colored by cellular states determined by LIGER. Entanglement scores are listed above the phylogenetic and phyloepigenetic trees and indicate whether labels share the same structure (score = 0) or exhibit unrelated structures (score = 1).

**Figure S13.**

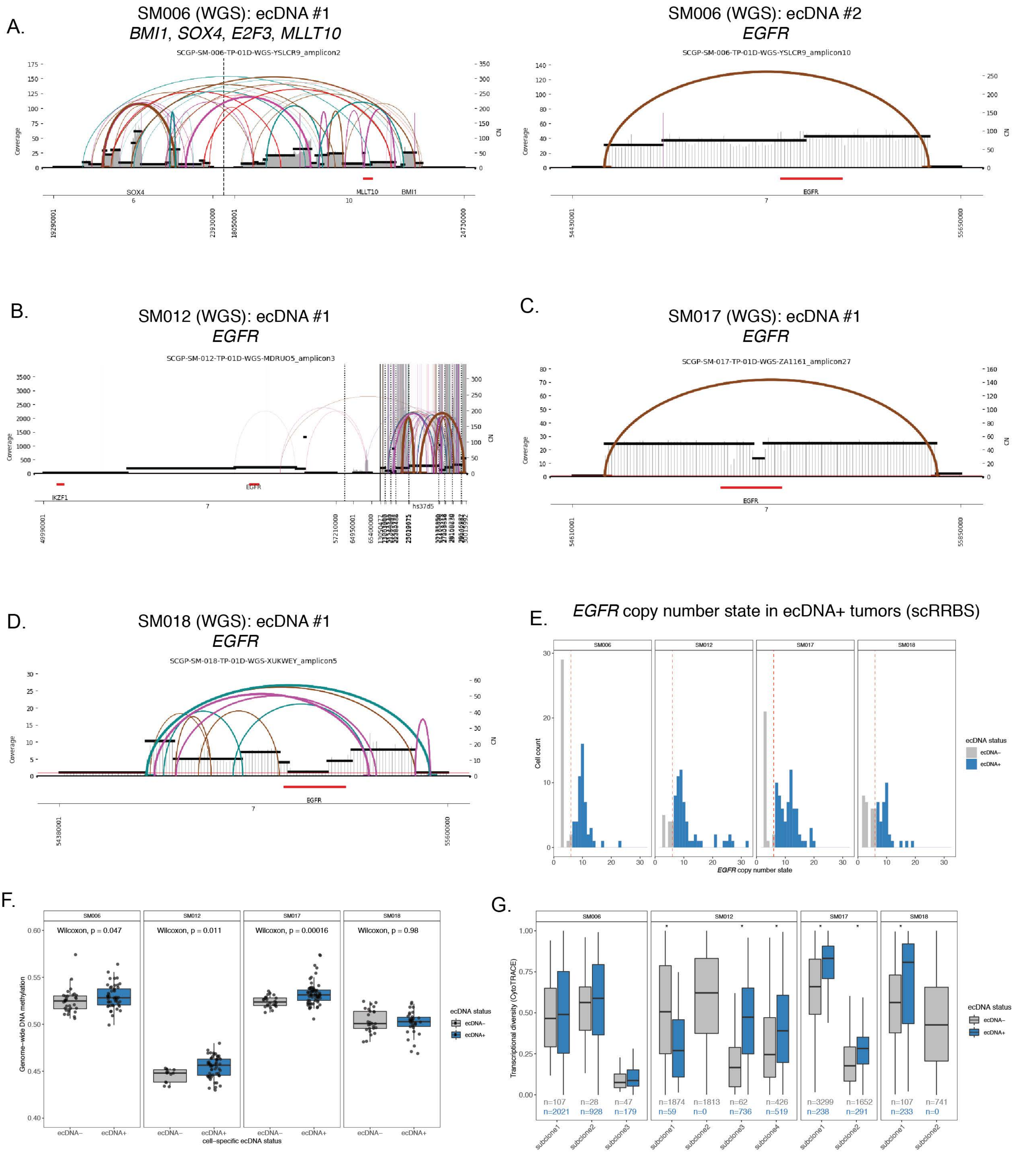
A.
SM006 (WGS): ecDNA #1
*BMI1*, *SOX4*, *E2F3*, *MLLT10*
SCGP-SM-006-TP-01D-WGS-YSLCR9_amplicon2
Coverage
CN
SOX4
MLLT10
BMI1
SM006 (WGS): ecDNA #2
*EGFR*
SCGP-SM-006-TP-01D-WGS-YSLCR9_amplicon10
EGFR
B.
SM012 (WGS): ecDNA #1
*EGFR*
SCGP-SM-012-TP-01D-WGS-MDRUO5_amplicon3
IKZF1
EGFR
hs37d5
C.
SM017 (WGS): ecDNA #1
*EGFR*
SCGP-SM-017-TP-01D-WGS-ZA1161_amplicon27
EGFR
D.
SM018 (WGS): ecDNA #1
*EGFR*
SCGP-SM-018-TP-01D-WGS-XUKWEY_amplicon5
EGFR
E.
*EGFR* copy number state in ecDNA+ tumors (scRRBS)
SM006
SM012
SM017
SM018
Cell count
*EGFR* copy number state
ecDNA status
ecDNA−
ecDNA+
F.
SM006
SM012
SM017
SM018
Wilcoxon, p = 0.047
Wilcoxon, p = 0.011
Wilcoxon, p = 0.00016
Wilcoxon, p = 0.98
Genome−wide DNA methylation
cell−specific ecDNA status
ecDNA status
ecDNA−
ecDNA+
G.
SM006
SM012
SM017
SM018
Transcriptional diversity (CytoTRACE)
n=107 n=2021
n=28 n=928
n=47 n=179
n=1874 n=59
n=1813 n=0
n=62 n=736
n=426 n=519
n=3299 n=238
n=1652 n=291
n=107 n=233
n=741 n=0
subclone1
subclone2
subclone3
subclone4
ecDNA status
ecDNA−
ecDNA+

**Figure S13: Focal extrachromosomal DNA amplifications generate greater levels of epigenetic and transcript diversity in glioma single cells. Related to Figure 4.**
(A-D) Extrachromosomal DNA circular amplicon reconstruction displaying genomic rearrangements predicted from whole genome sequencing. Coverage depth is represented as a histogram across a genomic interval with segment copy number (CN) estimation provided on the right y-axis. Discordant read pair clusters are indicated by arcs and colors highlight read pair orientation (e.g., brown = everted read pairs, (Deshpande et al., 2019). Amplicon intervals are provided at the bottom of the plot with annotation for known oncogenes (e.g., *EGFR*). (E) *EGFR* copy number estimation from single-cell RRBS data in ecDNA+ tumors. Cells with *EGFR* copy number greater than 7 were classified as *EGFR* ecDNA+ (blue). (F) Single-cell 10-kb tiled DNA methylation separated by *EGFR* ecDNA status. Single cells with inferred copy number status greater than 7 were classified as ecDNA+ (blue). Wilcoxon rank sum test *p*-values comparing DNA methylation across ecDNA status are reported for each patient tumor. (D) Boxplots depicting transcriptional diversity using gene count signatures calculated in scRNAseq data for each tumor, with cells separated based on inferred *EGFR* copy number status (gray = *EGFR* ecDNA-, blue = *EGFR* ecDNA+). Transcriptional diversity was compared based on predicted ecDNA status within each tumor subclone. Stars (*) indicate statistically significant differences based on Wilcoxon Rank Sum test ($p < 0.05$).

### Figure S14.

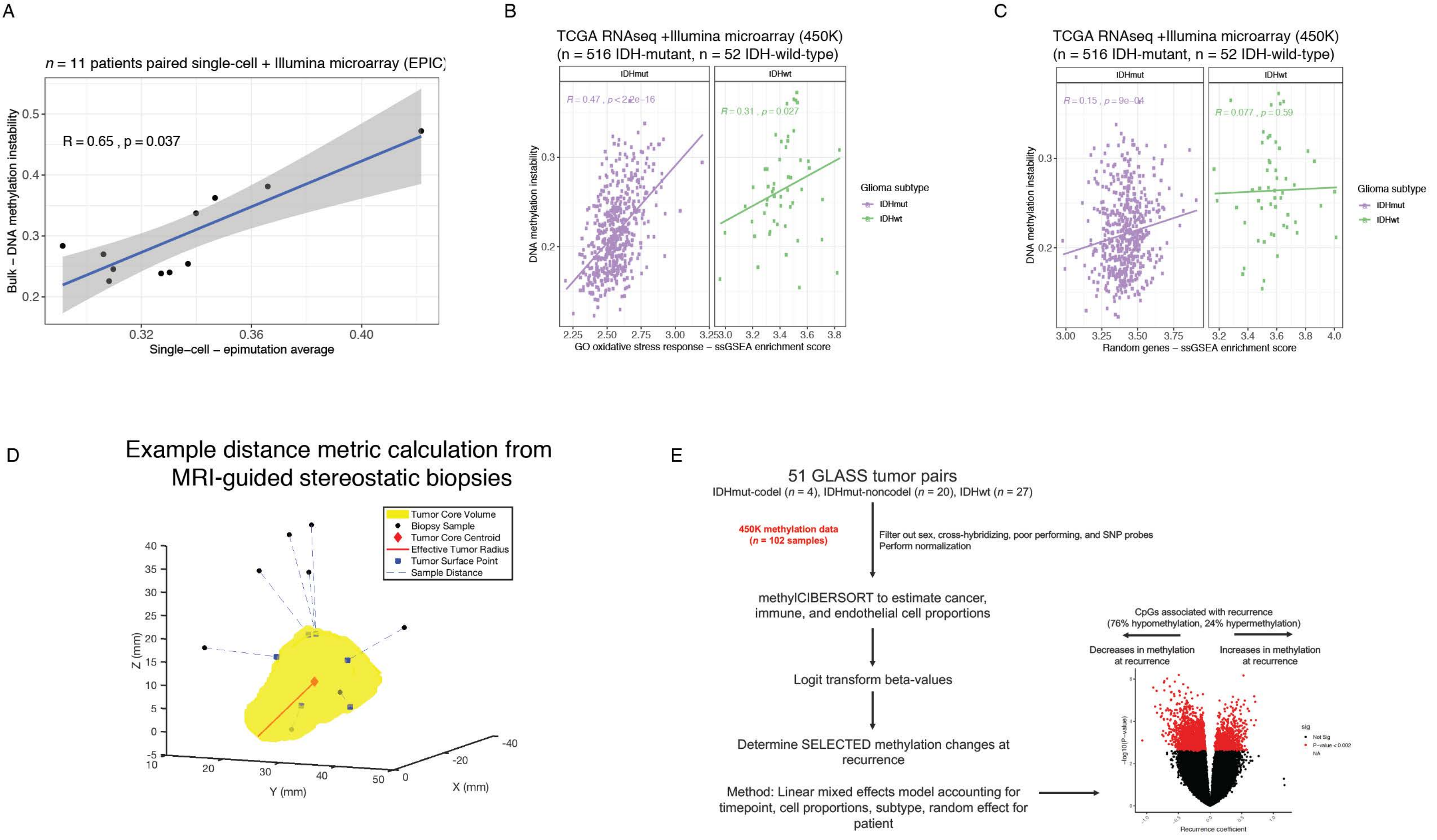


**Figure S14. DNA methylation instability metrics calculated in primary tumor, spatial, and longitudinal cohorts. Related to Figure 5.**

(A) Scatterplot highlights the significant positive correlation between the single-cell epimutation burden metric and the bulk microarray-based DNA methylation instability metric (n = 11 tumors, Spearman correlation). (B-C) Scatterplot between DNA methylation instability and (bulk RNAseq) ssGSEA enrichment scores for (B) oxidative stress response genes and (C) a randomly selected gene set substantiates finding that epigenetic instability is associated with stress response. (D) Schematic depiction of magnetic resonance image-guided biopsies and radiographic features used in spatial cohort (Verburg et al.). (E) Workflow for linear-mixed effect model identifying differentially methylated CpG sites that are selected for during tumor evolution when adjusting for estimated cellular proportions, glioma subtype, and a random effect for patient (n = 102 tumor samples, n = 51 patients).
